# A cargo receptor entrapment complex is a therapeutic node for genetically and clinically distinct proteinopathies

**DOI:** 10.64898/2026.07.10.737576

**Authors:** Magdalena Riedl Khursigara, Alissa C. Goss, Maria Kost-Alimova, Keith Keller, Christine Danielle De Mata, Ranjithmenon Muraleedharan, Edward Ryan Collantes, Matthew Brown, Elizabeth Grinkevich, Felichi Mae Arines, John Lin, Patrick Byrne, Silvana Bazua Valenti, Elizabeth Morici, Julie Roignot, Jason Zavras, Brianna R. Silverman, John Carlos Ignacio, Yoochan Myung, Seulki Kwon, Andrew Nelson, Hyery Yoo, Michelle Melanson, Matthew Racette, Valeria Padovano, Seth L. Alper, Dominique Carey, Namrata D. Udeshi, Steven A. Carr, Moran Dvela-Levitt, Toshio Narimatsu, Gustavo Sakuno, Victor San Martin Carvalho Correa, Nikolaos E. Efstathiou, Dimitrios P. Ntentakis, Tian Cao, Zhiqian Dong, Kim Thien Nguyen, Carolline Rodrigues Menezes, Ravi Kurumbail, Steven Kazmirski, Le Xiao, Hao Wu, David Liu, Sumaiya Iqbal, Eric S. Lander, Demetrios G. Vavvas, Krzysztof Palczewski, Juan Lorenzo B. Pablo, Anna Greka

**Affiliations:** Broad Institute of MIT and Harvard, Cambridge, MA, USA; Harvard/MIT MD-PhD Program, Harvard Medical School, Boston, MA, USA; Department of Medicine, Mass General Brigham and Harvard Medical School, Boston, MA, USA; Department of Medicine, Beth Israel Deaconess Medical Center and Harvard Medical School, Boston, MA, USA; The Mina and Everard Goodman Faculty of Life Sciences, Bar-Ilan University, Ramat-Gan, Israel; Retina Service, Department of Ophthalmology, Massachusetts Eye and Ear, Harvard Medical School, Boston, MA; Gavin Herbert Eye Institute - Brunson Center for Translational Vision Research, Department of Ophthalmology, University of California, Irvine, Irvine, CA, USA; Department of Physiology and Biophysics, University of California, Irvine, Irvine, CA, USA; Department of Biological Chemistry and Molecular Pharmacology, Blavatnik Institute, Harvard Medical School, Boston, MA, USA; Program in Cellular and Molecular Medicine, Boston Children’s Hospital, Boston, MA, USA; Department of Chemistry, University of California, Irvine, Irvine, CA, USA. Department of Molecular Biology and Biochemistry, University of California, Irvine, Irvine, CA, USA

## Abstract

Severe proteinopathies—such as retinitis pigmentosa, a form of inherited blindness—are driven by genetic mutations that overwhelm the quality control of the post-endoplasmic reticulum (post-ER) secretory pathway, causing toxic protein accumulation. Here, we identify a therapeutic node defined by a hetero-oligomeric cargo receptor complex consisting of TMED7, 2, 9, and 10. This “entrapment complex” anchors structurally and functionally diverse mutant clients within the early secretory pathway via TMED7 binding to the integral Golgi protein GRASP55. Disruption of the entrapment complex results in the clearance of accumulated protein cargoes. *In vivo* ablation of the entrapment node via inducible genetic deletion or via the small molecule BRD7635 reverses histopathological hallmarks and rescues functional deficits in clinically distinct proteinopathies of the kidney and the eye, including mitigating vision loss in a mouse model of retinitis pigmentosa.

## Introduction

More than a third of all human proteins rely on the secretory pathway for their proper folding and cellular localization^1–10^. Human proteinopathies are caused by mutations that overwhelm the cell’s proteostasis mechanisms^1–10^, leading to devastating and to-date incurable degenerative diseases such as retinitis pigmentosa^11^, an inherited form of blindness. While some misfolded proteins can be degraded by the proteasome^1,2,5,6^, less is known about how cells handle hundreds of disease-causing proteins that escape the ER and enter a post-ER secretory pathway. Addressing this knowledge gap may allow us to identify and target “nodes” that underlie multiple proteinopathies, thereby developing efficient and much needed therapies.

The transmembrane emp24 domain (TMED) proteins are known to monitor the folding state of secretory and membrane proteins and hold them back in the early secretory pathway if they are improperly folded^12–17^ – and yet, in an apparent contradiction, TMEDs have also been reported to be necessary for the canonical trafficking of secretory pathway cargoes as diverse as GPI-anchored proteins^13,16,18–26^, Wnt proteins^12,23,27,28^, toll-like receptors^14,29^, and G-protein coupled receptors^15,30^. A parsimonious explanation is that these seemingly opposing functions are served by different TMEDs or specific oligomeric TMED complexes. TMEDs comprise four subfamilies (α, β, γ, δ)^25,31–35^. In humans the α-subfamily comprises TMED9 and TMED4; the sole β is TMED2; the γ-subfamily is the largest comprising TMED1, TMED3, TMED5, TMED6, and TMED7; and the sole δ is TMED10^25,31–35^ (Figure S1A). However, (a) how specific TMEDs modulate the fates of mutant protein cargoes, and (b) whether targeting a distinct TMED complex could serve as a “nodal” therapeutic strategy for multiple clinically-diverse genetic proteinopathies are key questions that remained unexplored.

Here, we report that α, β, and δ TMEDs 9, 2 and 10 assemble with γ-TMED7 to form a hetero-oligomeric complex that specifically mediates misfolded cargo entrapment. TMED7 binds the Golgi protein GRASP55 via its cytoplasmic tail, which is essential for mutant cargo entrapment and accumulation in the early secretory pathway. *In vivo*, genetic deletion of TMED7 or the TMED-targeted small molecule BRD7635 ameliorated diseases caused by functionally and structurally distinct mutant proteins, including heretofore incurable genetic disorders that lead to kidney failure and blindness. Thus, the TMED7-containing hetero-oligomeric entrapment complex was revealed as a shared targetable “node” that may allow us to address multiple devastating proteinopathies with a single intervention.

## Results

### TMED depletion *in vivo* protects mice from blindness

We have previously shown that targeting TMED9 with the small molecule BRD4780 effectively removes intracellularly entrapped protein cargo^17^. Rhodopsin is a 7-transmembrane G-protein coupled receptor expressed in retinal rod photoreceptor cells^36^. Several rhodopsin (RHO) missense mutants, including the common RHO-P23H, accumulate intracellularly and cause premature photoreceptor cell death, leading to a form of retinitis pigmentosa (RP) characterized by progression to blindness^11,37,38^. TMED9 co-immunoprecipitated (Figure S1B) with three different disease-causing rhodopsin mutant proteins that accumulated intracellularly^38^, RHO-P23H, RHO-T17M and RHO-R135W, but not with the plasma membrane-localized mutant RHO-Q344X^38^ or the wild-type protein (Figure S1B-C). GFP-tagged RHO-P23H co-localized intracellularly with TMED9 whereas RHO-WT accumulated at the plasma membrane and did not colocalize with TMED9 (Figure S1D-E).

To explore the in vivo relevance of TMED9-mediated rhodopsin P23H entrapment, we crossed *Rho^P23H/+^* mice with mice engineered for inducible knockout of TMED9 (*Rho^P23H/+^*, *Tmed9^fl/fl^*, *CAGGCre–ER* mice; Figure 1A). Tamoxifen injection into these mice led to a decrease of TMED9 (TMED9 KO) by Western blot (Figure 1B). Inducible global deletion of TMED9 for up to 3 weeks in these adult mice was well tolerated; the TMED9 KO animals were grossly indistinguishable from their control littermates. Importantly, we observed decreased abundance of mutant RHO-P23H in TMED9 KO mice (Figure 1B). Since RP disease progression leads to the irreparable loss of photoreceptors and thinning of the outer nuclear layer (ONL), we obtained cross-sectional pictures of the retina and optic nerve using non-invasive Optical Coherence Tomography (OCT)^39,40^. OCT measurements showed a significantly thicker and preserved ONL in postnatal day 42 (P42) TMED9 KO mice (Figure 1C-D), indicating that disruption of TMED9 protected mice from photoreceptor loss. In the same mice, we also performed electroretinogram (ERG) recordings, a non-invasive test that measures light-induced retinal electrical activity^41^. We found that the b-wave amplitude, a functional measure of the inner retina processivity and a critical marker for photoreceptor synaptic health^42,43^, was significantly higher in TMED9 KO mice compared to controls (Figure 1E-F), consistent with preserved retinal function. In sum, we concluded that TMED9 KO leads to removal of mutant RHO-P23H from the retina and protection from retinal damage and vision loss.

**Figure 1.**
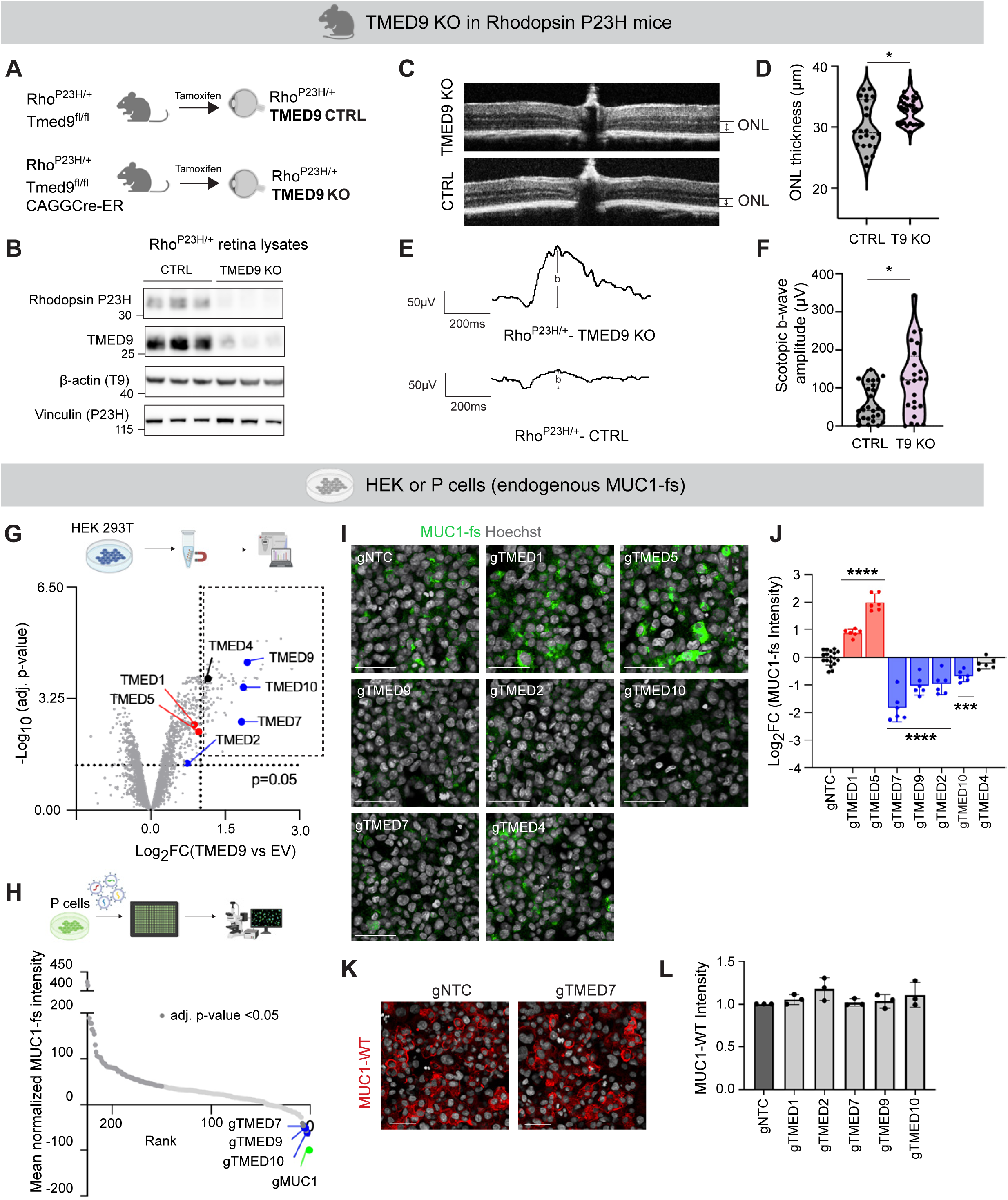
*In vivo*, inducible whole-body deletion of TMED9 mitigates progression to blindness, and *in vitro*, deletion of TMED9 and its interactors TMED2, 7 and 10 leads to clearance of mutant protein cargo. (**A**) Schematic of mouse model. *Rho^P23H/+^* mice were crossed with TMED9*^fl/fl^*mice either expressing CAGGCre-ER (TMED9 KO) or not (TMED9 CTRL). Whole-body Cre expression and genetic knockout was induced with Tamoxifen (3 doses, 96 mg/kg). (**B**) Western blot for RHO-P23H and TMED9 in retinal lysates from TMED9 KO or CTRL mice. Loading controls, ß-actin (for TMED9 (T9)) and vinculin (for RHO-P23H (P23H)). Each lane represents one animal. (**C**) Representative OCT images measuring the outer nuclear layer (ONL) taken at P42 in TMED9 KO and CTRL mice. (**D**) Violin plot depicting ONL thickness in both eyes of 14 TMED9 KO and 11 CTRL mice. ONL Median and quartiles shown. * p < 0.05. (**E**) Representative ERG waves from TMED9 KO and CTRL mice. Interval schematic indicates the b-wave distance measured. **(F**) Violin plot depicting b-wave amplitude in both eyes of 12 TMED9 KO and 13 CTRL mice. Median and quartiles shown. * p < 0.05. (**G**) IP-MS of TMED9-MYC from HEK293T cells. Volcano plot comparing −Log_10_ adj. p-values and Log_2_ fold change in protein abundance between TMED9-MYC and empty vector (EV) conditions. N = 3 technical replicates. Entrapment TMEDs in blue, non-entrapment TMEDs in red. (**H**) Arrayed CRISPR/Cas9 knockout screen of TMED9-MYC interactors. Mean normalized MUC1-fs intensity per gRNA shown. N = 4 technical replicates. 2,000-8,000 cells analyzed per gRNA. TMED gRNAs (blue) resulting in significant reduction of MUC1-fs cargo; MUC1 gRNA (green), positive control; gRNAs with adjusted p-value < 0.05 (dark gray). (**I**) Immunofluorescence (IF) images of endogenous MUC1-fs (green) in human P cells with stable depletion of each TMED as indicated (gTMED**X**). Scale = 50 μm. (**J**) Quantification of (I). Log_2_ fold change from NTC of mean MUC1-fs intensity/well is shown. N = 6 technical replicates. 540 - 1620 cells analyzed per stable P cell line. Significant increases (non-entrapment TMEDs, red) and decreases (entrapment TMEDs, blue) in mean MUC1-fs intensity are indicated. Means ± SD. *** = p < 0.001; **** = p < 0.0001. (**K**) Representative IF images of endogenous wild-type mucin 1 (MUC1-WT, red) in non-targeting control (gNTC) and TMED7 depleted (gTMED7) P cells. Scale = 50 μm. (**L**) Quantification of (K) and MUC1-WT intensity in all other TMED depleted P cells. Comparison to gNTC was non-significant. N = 4 technical replicates. 2,000-8,000 cells analyzed per gRNA.

### “Entrapment” versus “non-entrapment” TMEDs have antagonistic effects on cargo abundance and directionality of cargo trafficking

To more deeply probe TMED cargo receptor biology, we next sought to comprehensively identify TMED9-interacting partners by immunoprecipitation-mass spectrometry (IP-MS, Figure 1G). We identified > 500 significantly enriched TMED9 interactors, including additional members of the TMED family (Table S1), a finding that motivated an investigation into the role of the entire TMED family in the regulation of misfolded protein cargo. Since overexpression of TMEDs and their cargoes is a known confounder due to their exuberant ectopic accumulation in secretory compartments^44,45^, we employed a well-validated endogenous cell system: patient-derived human kidney epithelial cells (P cells; cells obtained from a patient with kidney disease harboring a frameshift *MUC1* mutation resulting in endogenous mutant MUC1-fs intracellular protein accumulation)^17^. In this cell model, endogenous TMED9 binds to endogenously expressed misfolded protein (MUC1-fs, Figure S2A) and other endogenous TMEDs (Figure S2B). We depleted the top 105 TMED9 interactors from the IP-MS experiment (Figure 1G and Table S1, based on log_2_ fold change) in Cas9-positive P cells and measured MUC1-fs abundance using high content multiplexed imaging (Figure 1H)^17^. TMED knockouts were the top hits for MUC1-fs clearance (Figure 1H and Table S2).

We engineered P cell lines with stable depletion of each TMED that interacts with TMED9 (Figure 1I-J and Figure S2C). Depletion of TMED7, TMED2, TMED10 and TMED9 itself led to a decrease in MUC1-fs; therefore we termed these four “entrapment” TMEDs, because their presence promoted misfolded cargo accumulation and their depletion allowed for cargo removal. Surprisingly, depletion of TMED1 and TMED5 (red bars) led to the opposite: a significant increase in MUC1-fs accumulation. We therefore termed these two “non-entrapment” TMEDs, because their presence promoted cargo removal and their depletion allowed for cargo accumulation (Figure 1I-J). TMED4 depletion had no effect on mutant cargo. In support of our immunofluorescence studies, the depletion of entrapment TMEDs (blue label) led to reduction of the partially glycosylated, primary MUC1-fs band^17^ in Western blots (Figure S2D, arrowhead), consistent with the release of MUC1-fs from the early secretory pathway. Also in accord with our imaging results, depletion of non-entrapment TMEDs (red label) led to an increase in the immature MUC1-fs band, suggesting increased accumulation of partially glycosylated MUC1-fs in the early secretory pathway (Figure S2D, arrowhead). Importantly, TMED depletion in no way affected the plasma-membrane localization and abundance of wild-type Mucin 1 (MUC1), suggesting that (a) canonical wild-type protein trafficking was not disrupted by the depletion of individual TMEDs in these studies and (b) TMED depletion did not downregulate MUC1 transcriptionally (because both mutant and wild-type MUC1 share the same promoter and are nearly indistinguishable at the mRNA level; Figure 1K-L). Turning our attention to misfolded rhodopsin cell models, we confirmed that TMED7 and TMED9 depletion resulted in a reduction of mutant RHO-P23H, whereas TMED5 depletion resulted in accumulation of RHO-P23H, further solidifying the phenotypic differences in cargo handling between entrapment and non-entrapment TMEDs (Figure S2E-H).

We predicted that disruption of the entrapment TMEDs would allow misfolded cargo to be released and proceed through the secretory pathway (Figure S3A-B). Indeed, we observed that MUC1-fs in entrapment TMED-depleted cells (Figure 1I-J) was not only reduced in abundance overall, but had also redistributed along the secretory pathway (Figure S3C-E). In fact, the remaining MUC1-fs was highly colocalized with late secretory pathway organelles, *e.g.* the endosome or lysosome (Figure S3C-E). On the contrary, depletion of non-entrapment TMEDs increased MUC1-fs accumulation overall, and augmented its colocalization with early secretory pathway structures (Figure S3D-E). Further, removal/degradation of misfolded cargo was dependent upon an intact secretory pathway. In entrapment TMED depleted cells, thapsigargin (100 nM, SERCA pump inhibitor/ER stressor), brefeldin A (200 ng/mL, Golgi disruptor), or bafilomycin A1 (100 nM, inhibitor of lysosomal acidification) led to re-accumulation of MUC1-fs at different molecular sizes representing different degrees of glycosylation^17^, consistent with disrupted cargo trafficking at different points along the secretory pathway (Figure S3F). Bortezomib (50 nM), a proteasome inhibitor, had no effect on MUC1-fs abundance, confirming no role for proteasomes in cargo degradation.

### Entrapment TMEDs form a hetero-oligomeric complex that is disrupted by the small molecule BRD7635

Since TMEDs are known to be interdependent in part due to their propensity to form oligomeric complexes^23,46–48^, we transfected equimolar amounts of TMED7, 2, 9 and 10 with different tags into Expi293F GnTI cells, followed by purification using a FLAG column (TMED9), His column (TMED2) and S6 size exclusion column (Figure 2A). Even after two subsequent rounds of affinity purification for a singular TMED, all 4 entrapment TMEDs remained stably complexed as seen in Coomassie staining of elution fractions (Figure 2B). Protein peaks from the size exclusion column indicated large oligomeric conformations for the TMEDs (Figure 2C) whose presence after this third round of purification was confirmed by Western blotting for each of the 4 protein tags (Figure 2D). We thus concluded that TMED7/2/9/10 likely formed a specific hetero-oligomeric entrapment complex.

**Figure 2:**
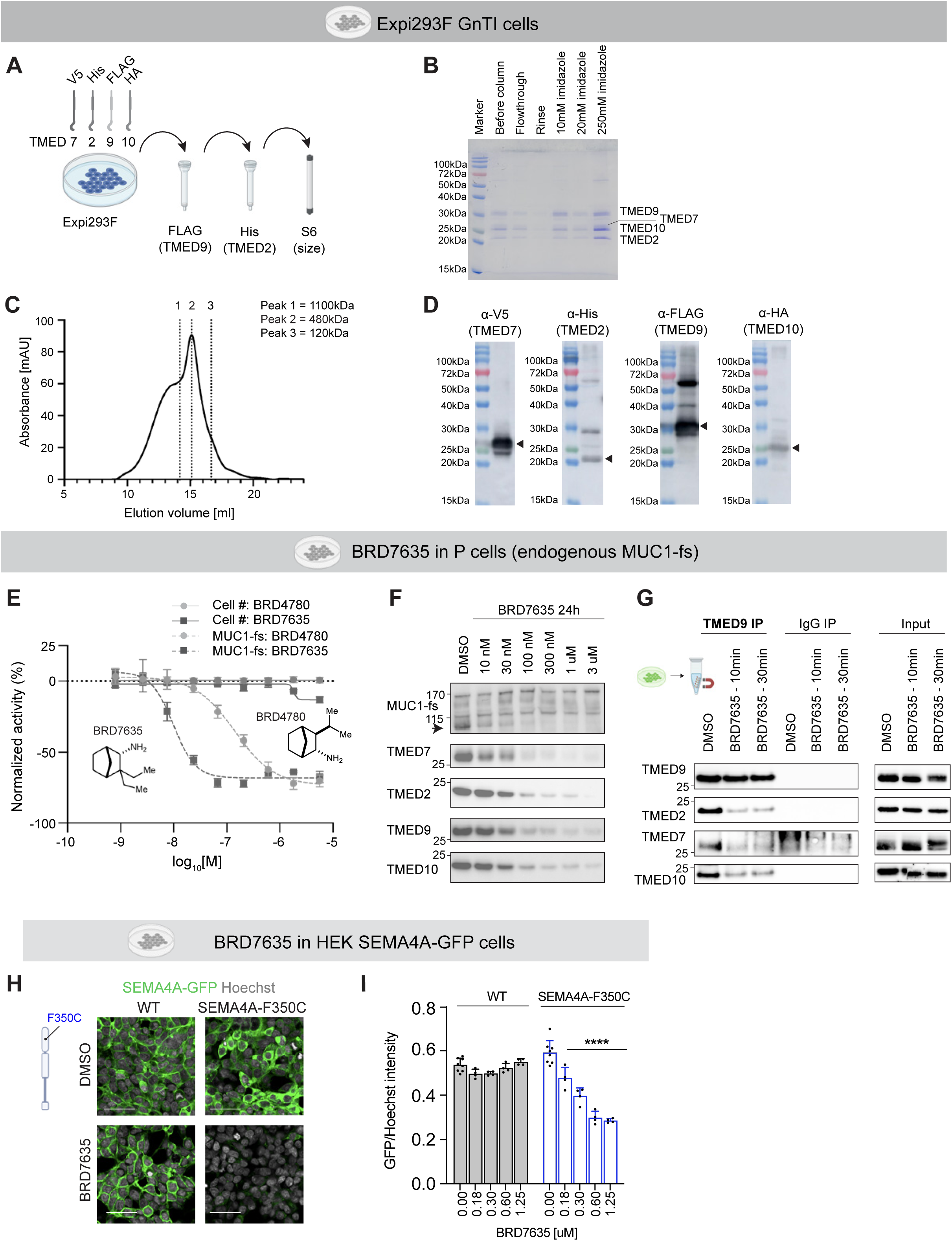
“Entrapment” TMEDs 2, 7, 9 and 10 form hetero-oligomeric complexes which are disrupted by the small molecule BRD7635. (**A**) Schematic of TMED7/2/9/10 purification procedure after overexpression in Expi293F GnTI cells. (**B**) Coomassie staining of protein fractions from the Ni-NTA column. TMEDs were labelled according to their expected size. (**C**) 250 mM imidazole eluate was run on an S6 size-exclusion column. Approximate molecular weights based on a set of known protein standards are indicated with perforated lines. (**D**) Size-exclusion column protein fractions were probed with antibodies against each specific TMED tag. (**E**) Dose-response curves of P cell number (solid lines, indicating no toxicity-related cell loss) and MUC1-fs IF intensity following 48 h treatment with legacy compound BRD4780 (light gray, dashed line) or new structure-activity optimized compound BRD7635 (dark gray, dashed line). Means ± SD. For MUC1-fs degradation BRD4780 IC_50_ = 141.0 nM vs. BRD7635 IC_50_ = 9.9 nM. N = 3-9 technical; 3 biological replicates. (**F**) Western blot of endogenous MUC1-fs (primary MUC1-fs band indicated by arrowhead) and endogenous TMED levels in P cells following 24 h treatment with BRD7635. (**G**) Co-IP of endogenous TMED9 with TMED2, 7 or 10 in P cells treated with 5 μM BRD7635 for 10 or 30 min. Lysate inputs, right. One representative of 3 biological replicates is shown. (**H**) Schematic of semaphorin 4A with F350C variant on the left. Representative images of GFP-tagged WT or F350C semaphorin 4A HEK293T cells treated with either DMSO or BRD7635 at 1.25 μM for 5 days. Scale = 50 µm. (**I**) Quantification of GFP/Hoechst signal in GFP-tagged wild-type semaphorin 4A (WT) or F350C semaphorin 4A cells after treatment with the indicated BRD7635 concentrations. Means ± SD. N = 4-8 technical replicates. 60,000 - 72,000 cells analyzed per genotype. **** = p < 0.0001.

A structure-activity relationship (SAR) campaign starting with BRD4780 resulted in our discovery of BRD7635, a compound that was 14-fold more potent in reducing mutant cargo than BRD4780^17^ (Figure 2E) without significant off-target effects across a standard panel of targets (Table S3). In addition to reducing TMED9 abundance, BRD7635 led to the reduction of all tested TMEDs (Figure 2F, Supplemental Figure 4A-B). This led us to hypothesize that BRD7635 might bind to and disrupt the TMED entrapment complex. We performed nanoscale differential scanning fluorimetry (nanoDSF, Figure S4C) with increasing concentrations of BRD7635 on the purified hetero-oligomeric TMED7/2/9/10 complex (Figure 2C). These experiments revealed large dose-responsive shifts in melting temperature (Figure S4C), consistent with binding of BRD7635 to the entrapment complex. BRD4780 induced smaller shifts when binding to the complex in line with its weaker efficacy for MUC1-fs removal (Figure 2E, Figure S4C). Furthermore, the mechanism of action of BRD7635 depended on the entrapment complex. We co-immunoprecipitated endogenous TMED9 from cells treated with BRD7635 for ten minutes (before onset of drug-induced TMED degradation) and found decreased TMED9 binding to TMED2, TMED7 and TMED10, indicating disruption of the entrapment complex (Figure 2G). Concomitant treatment of cells with BRD7635 and either brefeldin A (200 ng/mL), bafilomycin A1 (100 nM,) or MG132 (2 µM, a proteasome inhibitor, used as a negative control) revealed that BRD7635 promoted lysosomal degradation of the entrapment TMEDs (Figure S4D-E).

To further assess the general applicability of BRD7635 for proteinopathies, we looked at mutant semaphorin 4A, another cause of inherited blindness^49^ due to accumulation of misfolded cargo. BRD7635 treatment of cells expressing either wild-type or mutant F350C semaphorin 4A specifically removed intracellularly accumulated mutant F350C–but not wild-type semaphorin–in a dose-dependent fashion (Figure 2H-I). With regard to the specificity of this compound for the removal of mutant, intracellularly accumulated proteins–and not their wild-type versions–BRD7635 behaved like the previously published TMED9-targeted compound BRD4780 that did not clear or alter the membrane localization of wild-type glycoprotein mucin 1 or wild-type GPI-anchored uromodulin^16,17^. All together, the compound BRD7635 cleared disease-causing mutant cargoes in cell models for proteinopathies affecting the kidney and the eye.

### Disruption of TMED7 is sufficient to release entrapped cargo

The efficacy of BRD7635 in disrupting the TMED complex prompted us to more deeply probe the relationship between each of the components of the entrapment complex, namely how disrupting each of the four entrapment TMEDs might affect the others and their ability to handle cargo. In “double knockout” studies, we acutely knocked out individual TMEDs on a background of either stable TMED5 P cell depletion (i.e. increased accumulated cargo at baseline) or TMED9 P cell depletion (i.e. decreased cargo at baseline). Knockout of non-entrapment TMED1 and TMED5s did not increase cargo accumulation or otherwise change the phenotype in stable TMED9-depleted cells (i.e. MUC1-fs did not reaccumulate; stable gTMED9+gTMED5)(Figure 3A-B, red box). In contrast, knockout of entrapment TMEDs in stable TMED5-depleted cells reversed the phenotype (i.e. MUC1-fs was diminished; stable gTMED5+gTMED7)(Figure 3A-B, blue box). These experiments showed that the disruption of each entrapment TMED is equally dominant in terms of cargo handling–and yet a key unanswered question was how their disruption might affect the stability of all other TMEDs.

**Figure 3.**
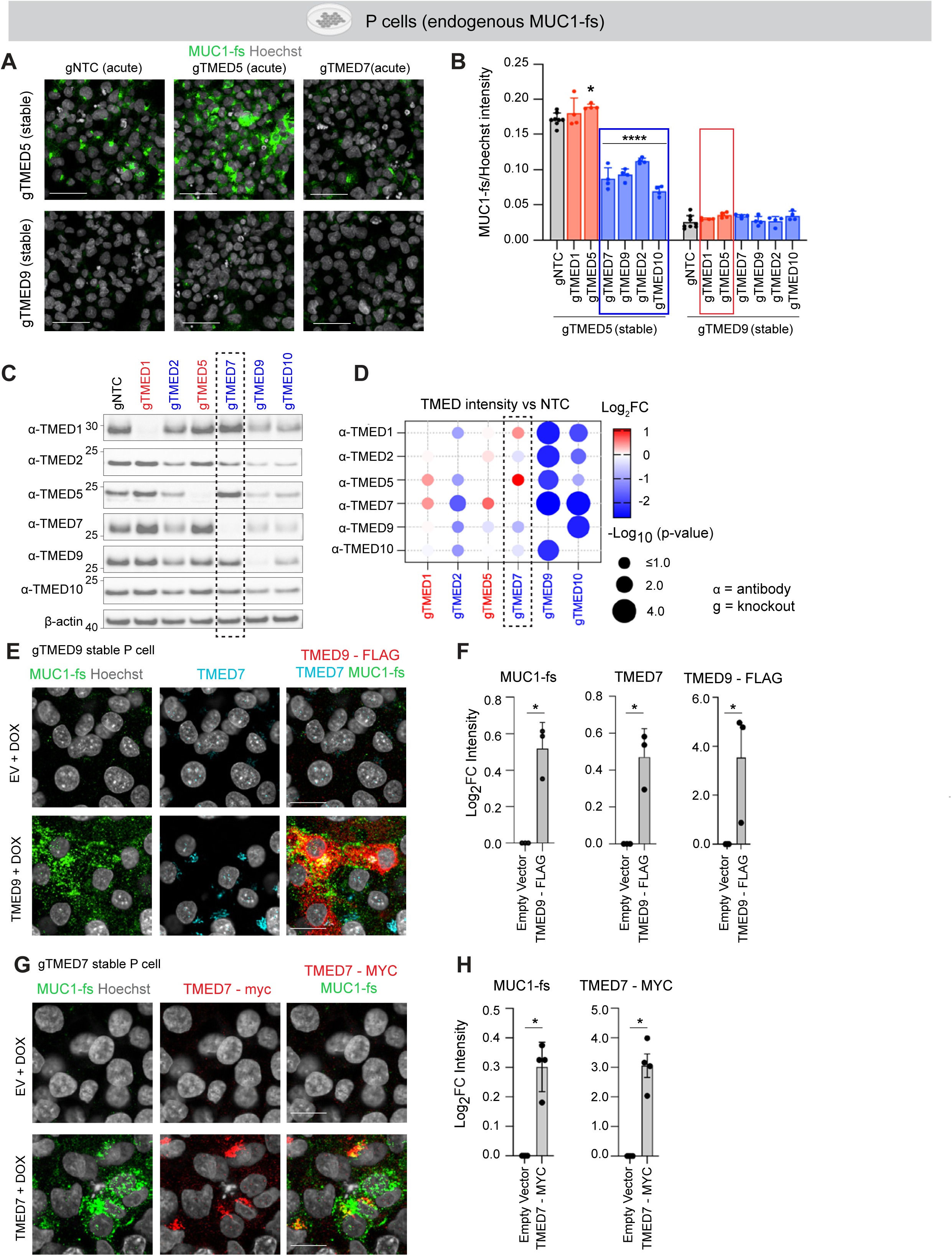
TMED7 is required to form a hetero-oligomeric TMED7/2/9/10 complex that mediates mutant cargo entrapment. (**A**) “Double knockout” experiments: IF images of endogenous MUC1-fs (green) in stable TMED5-depleted (top) or TMED9-depleted (bottom) P cells acutely transduced with the TMED gRNAs indicated in the column labels. Scale = 50 μm. (**B**) Quantification of (A). MUC1-fs intensity/Hoechst intensity per well is shown, compared to gNTC (control). Blue, entrapment TMEDs; Red, non-entrapment TMEDs. Means ± SD. N = 4 technical replicates. 20,230 - 40,880 cells per genotype analyzed. * = p < 0.05, **** = p < 0.0001, NTC vs gTMED. Blue box shows that depletion of each of four entrapment TMEDs is dominant over stable TMED5 depletion, reversing the MUC1-fs phenotype. Red box shows that depletion of each of two non-entrapment TMEDs is not dominant over stable TMED9 depletion and does not reverse the MUC1-fs phenotype. (**C**) Representative Western blots showing the abundance of endogenous TMEDs in TMED-depleted P cells. Knockout is labelled on the x-axis (gTMED) and the antibody recognizing each TMED on the y-axis (*α*-TMED). Black perforated box indicates minimal effect on the abundance of other TMEDs after depletion of TMED7 (gTMED7). (**D**) Quantification of (C) as a dot plot (endogenous TMED signal; Pseudocolor, Log_2_ fold change of TMED abundance vs. Control (NTC); Circle size, Log_10_ adjusted p-value). N = 3 biological replicates. Black perforated box indicates minimal effect on the abundance of other TMEDs after depletion of TMED7 (gTMED7). (**E**) IF images of endogenous MUC1-fs (green) and endogenous TMED7 (cyan) in stable TMED9-depleted P cells transduced with doxycycline-inducible empty vector or TMED9-FLAG (red). Scale = 20 μm. (**F**) Quantification of (E). Log_2_ fold change of MUC1-fs (left), TMED7 intensity (middle) or TMED9-FLAG intensity (right) compared to empty vector. Means ± SD. N = 3 biological replicates. 12,000 - 15,000 cells analyzed per condition per replicate. * = p < 0.05. (**G**) IF images of endogenous MUC1-fs (green) in stable TMED7-depleted P cells transduced with doxycycline-inducible empty vector or TMED7-MYC (red). Scale = 20 μm. (**H**) Quantification of (G). Log_2_ fold change of MUC1-fs (left) and TMED7-myc intensity (right) compared to empty vector. Means ± SEM. N = 4 biological replicates. 10,000 - 15,000 cells analyzed per condition per replicate. * = p < 0.05.

To probe how depletion of each TMED affects the stability and abundance of all the others, we quantified the abundance of each TMED after individual depletion of each other TMED by Western blot. Depletion of the entrapment TMEDs 2, 9 and 10 each led to decreased stability and abundance of all other TMEDs (Figure 3C-D). Intriguingly, loss of TMED7 was not broadly destabilizing to the other TMEDs (perforated box, Figure 3C-D) and yet was equally as effective at reducing cargo accumulation as any other entrapment TMED (Figure 1I-J). Of all the entrapment TMEDs, we noted that TMED7 was specifically and tightly correlated with MUC1-fs accumulation, e.g. gTMED5 cells with high MUC1-fs also had high TMED7 levels (Figure S5A-B). Further, re-introduction of tagged doxycycline-inducible TMED9 in TMED9-depleted cells (that had diminished TMED7, Figure S5A-B) resulted in re-accumulation of mutant cargo and, concomitantly, a significant increase in TMED7 abundance (Figure 3E-F). Most importantly, TMED7 was sufficient for cargo entrapment in gain-of-function studies: we transduced gRNA-resistant TMED7-myc into TMED7-depleted P cells and found that re-introduction of TMED7 was sufficient to induce significant re-accumulation of mutant cargo (Figure 3G-H). We concluded that TMED7 was the crucial component of the TMED complex conferring entrapment, therefore we focused our attention on TMED7.

### TMED7 binds to Golgi membrane protein GRASP55 to promote cargo entrapment

We reasoned that TMED7 might uniquely interact with another protein to facilitate cargo entrapment. To investigate this hypothesis, we performed IP-MS experiments with MYC-tagged TMED7 versus TMED5 as a non-entrapment control (Figure 4A and Table S4). A large subset of TMED7 interactors, unlike TMED5 interactors, were Golgi membrane proteins. This finding was consistent with high-throughput high-content imaging studies (∼10,000 single cells per marker or condition tested; Figure S5C-D, Table S5) showing that entrapment TMEDs were more heavily Golgi-localized than non-entrapment TMEDs (Figure S5C-D). Cargo binding was not sufficient to explain the differential localization, because all tested TMEDs interacted with MUC1-fs (Figure S5E), an interaction likely mediated by the GOLD domain (Figure S5F)^34^.

**Figure 4.**
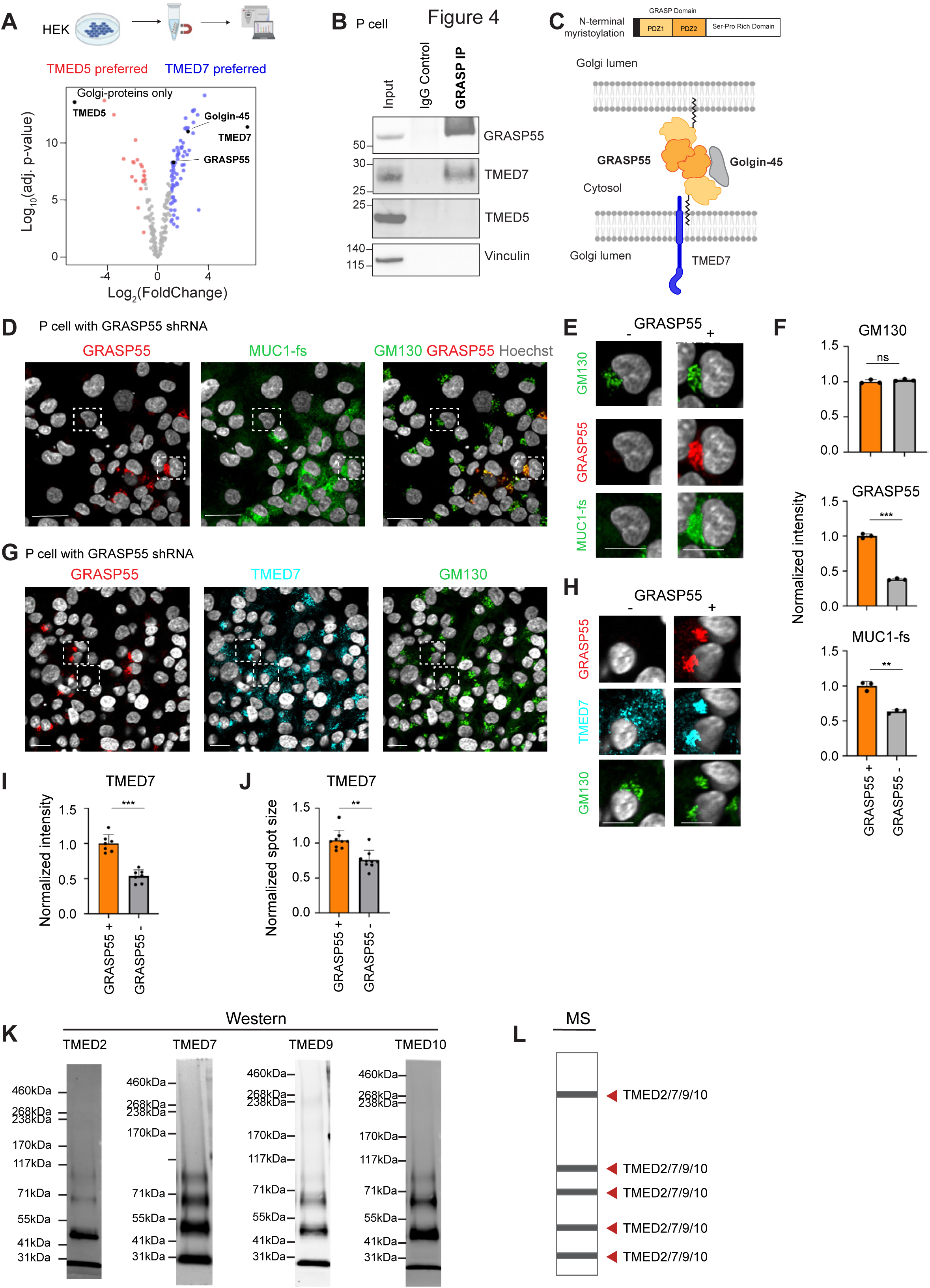
TMED7 interacts with the integral Golgi protein GRASP55 to mediate cargo entrapment. (**A**) Volcano plot of IP-MS experiment in HEK239T cells overexpressing TMED7 or TMED5-MYC. Analysis identified Golgi-localized proteins that preferentially (p<0.05 and Log_2_FC>1) bind TMED7 (blue) or TMED5 (red). Proteins not significantly enriched are in grey. N= 4 technical replicates. (**B**) Endogenous GRASP55 Co-IP probed for TMED7 and TMED5. N=3 biological replicates. (**C**) Schematic of TMED7 and the PDZ1 and PDZ2 domains of GRASP55, anchored to the Golgi membrane via myristoylation. Golgin-45 (grey) binds GRASP55. (**D**) GRASP55 depletion in a field of human P cells. White boxes highlight GRASP55 negative (-) and GRASP55 positive (+) cells. IF of GRASP55 (red, left), MUC1-fs (green, middle), and overlaid GM130 (pseudocolored green) and GRASP55 (right; colocalization of GM130 and GRASP55, orange). Scale = 50 μm. (**E**) Higher magnification images to visualize cells in white perforated boxes from (D). Scale = 10 μm. (**F**) Quantification at the single cell level for GM130 (top), GRASP55 (middle) and MUC1-fs (bottom) intensities in GRASP55 negative cells (-) (8,880 - 9,500 cells per replicate) vs. GRASP55 positive (+) cells (1,845 - 2,565 cells per replicate). Means ± SD. ns = non significant, ** = p < 0.01, *** = p < 0.001. N = 3 biological replicates. **(G)** GRASP55 depletion in a field of human P cells. White perforated boxes highlight GRASP55 negative (-) and GRASP55 positive (+) cells. IF images of GRASP55 (red, left), TMED7 (cyan, middle), and GM130 (green, right). Scale = 20 μm. (**H**) Higher magnification images to visualize cells in white perforated boxes from (G). Scale = 10 μm. **(I)** Quantification at the single cell level for TMED7 in GRASP negative (-) cells (400 - 2,250 cells per replicate) vs. GRASP positive (+) cells (115 - 700 cells per replicate). Means ± SD. * = p < 0.05, ** = p < 0.01, *** = p < 0.001. N = 6-9 technical replicates. (**J**) Quantification of TMED7 spot size as a measure of dispersion in GRASP negative (-) cells (400 - 2,250 cells per replicate) vs. GRASP positive (+) cells (115 - 700 cells per replicate). Means ± SD. ** = p < 0.01. N = 6-9 technical replicates. **(K)** Golgi-IP lysates were crosslinked with 5 mM DSSO and probed by Western blot for TMED2, 7, 9 and 10. N = 3 biological replicates. (**L**) MS of bands cut-out from Golgi-IP lysates ran on a PAGE gel confirmed that all four TMEDs were present in each of the bands, as indicated by the red arrows. N = 3 technical replicates.

Among the Golgi-localized TMED7 interactors, we identified significant interactions with the integral Golgi proteins Golgin-45 and GRASP55 (Golgi reassembly-stacking protein of 55 kDa)(Figure 4A). We confirmed binding of endogenous GRASP55 to endogenous TMED7 – but not TMED5 (Figure 4B; aligned sequence, Figure S6A-B; positive control Co-IP for TMED5 and its known interactor COPB2, Figure S6C). GRASP55 also co-immunoprecipitated with each of the other three entrapment TMEDs (Figure S6D), suggesting that GRASP55 is a specific entrapment complex interactor. GRASP55 (encoded by *GORASP2*) contains two PDZ domains and aids in Golgi stack formation through homodimerization and interaction with Golgin-45^50^ (Figure 4C). Importantly, at the single cell level, acute GRASP55 depletion mitigated MUC1-fs cargo accumulation without perturbing the integrity of the Golgi apparatus (Figure 4D-F; no change to Golgi marker GM130) or other canonical secretory pathway cargo proteins, including the wild-type glycoprotein MUC1 (Figure S6E-F, no change to MUC1-WT abundance and localization). We also found that TMED7 intensity was decreased in GRASP55 depleted cells (Figure 4G-I). In the absence of GRASP55, TMED7 appeared dispersed and localized away from the Golgi compartment (Figure 4H, 4J), suggesting that GRASP55 anchored TMED7 to the Golgi.

Since the interaction between TMED7 and the Golgi-localized GRASP55 was important for cargo entrapment, we reasoned that we should be able to isolate the entire entrapment complex biochemically in a cellular fraction enriched for Golgi localized protein complexes. We employed P cells stably expressing HA-tagged TMEM115, to allow isolation of intact Golgi organelles (Golgi IP). While the ER marker calreticulin was depleted, suggesting we had a clean Golgi-IP, all 4 entrapment TMEDs were indeed greatly enriched (Figure S6G-H). To investigate the oligomerization of these TMEDs, we chemically crosslinked Golgi-enriched protein complexes with DSSO. Probing by Western blot, we found the entrapment TMEDs 2, 7, 9 and 10 in multiple oligomeric states (Figure 4K). We confirmed by MS that all members of the entrapment complex TMED 2, 7, 9, and 10 were present in crosslinked samples from every molecular weight range (Figure 4L). Taken together, these experiments showed that GRASP55 mediated the Golgi localization of entrapment TMED hetero-oligomers.

### The TMED7 cytoplasmic tail and coiled-coiled domain mediate binding to GRASP55

Since GRASP55 is cytoplasmic (Figure 4C), we hypothesized that it interacts with TMED7 via the TMED7 cytoplasmic tail (C-tail). Indeed, TMED7 C-tail deletion abrogated TMED7 binding to GRASP55 (Figure 5A). We performed molecular dynamics (MD) simulations initialized from the AlphaFold3 predicted interfaces between either the C-tail of TMED7 (entrapment TMED) or TMED5 (non-entrapment TMED), and GRASP55 (Figure S6I), to evaluate residue-level interactions. The TMED7 C-tail was predominantly disordered and flexible in part due to a string of threonines (T217-T220), and engaged extensively with the GRASP55 PDZ2 domain via salt-bridges (Figure 5B, left). In contrast, the TMED5 C-tail showed sparse contacts with GRASP55, because the spatial confinement of charged residues within the TMED5 α-helical structure restricted their ability to engage in multiple salt-bridges (Figure 5B, right).

**Figure 5.**
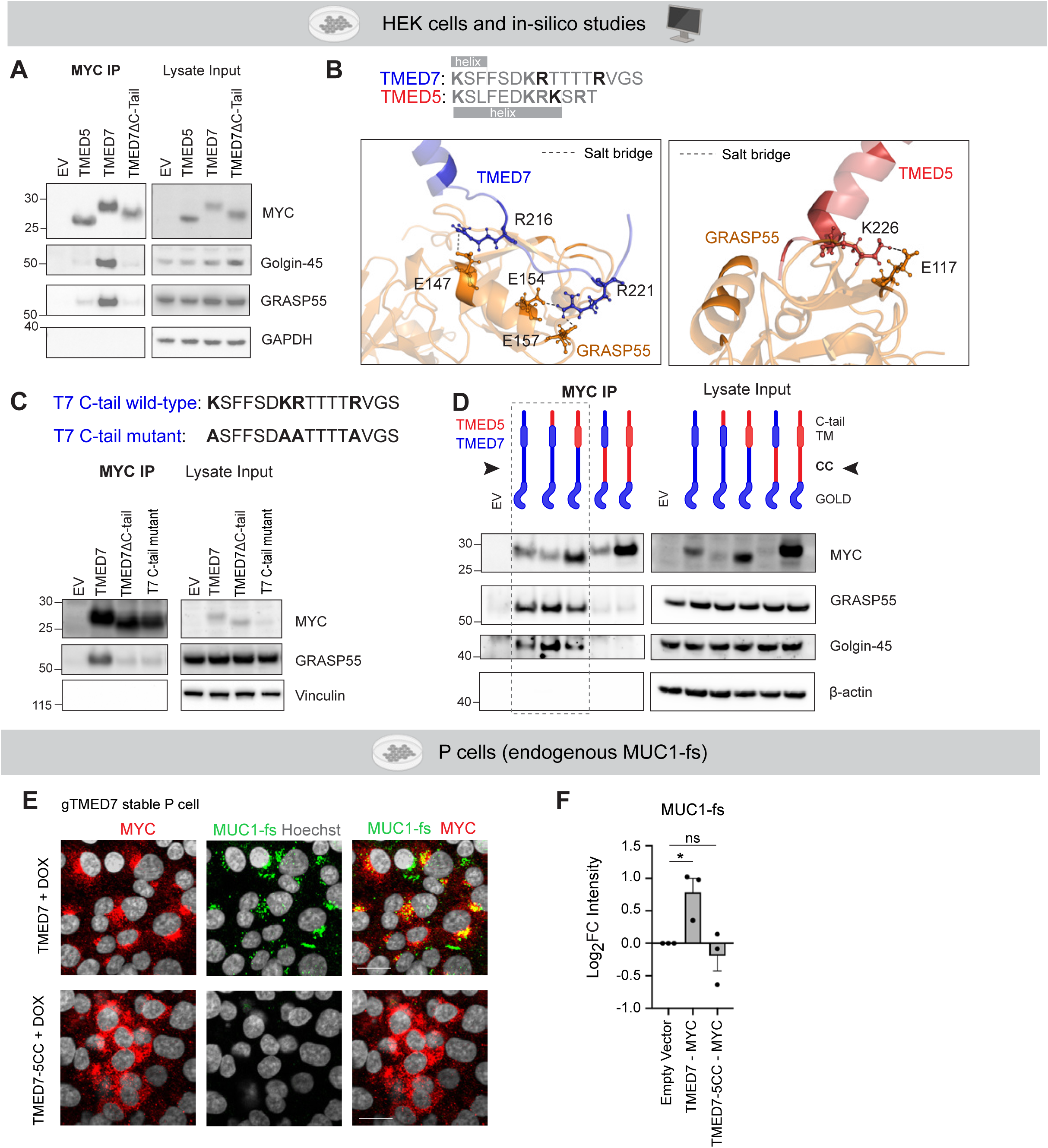
TMED7 binds to GRASP55 via critical residues on its cytoplasmic tail. (**A**) MYC Co-IP for TMED5-MYC, TMED7-MYC and TMED7ΔC-tail-MYC probing for pull down of Golgin-45 and GRASP55. Input on the right. (**B**) Sequence alignment of the C-tails of TMED7 and TMED5. Basic residues (Lys and Arg) are grey and bold; basic residues predicted to form salt bridges are highlighted in black and bold. Helical regions are indicated with a grey box. (Left) MD simulation snapshots of the most representative conformations for the TMED7-GRASP55 interface, where R216 and R221 in TMED7 (blue) were predicted to form salt bridges (dashed lines) with E147, E154, and E157 in GRASP55 (gold). (Right) At the TMED5-GRASP55 interface, only K226 in TMED5 (red) was predicted to form salt bridges with GRASP55 (gold), interacting with a negatively charged region coordinated by E117. Additional basic residues on TMED tails and glutamic acids on GRASP55 rendered with opaque coloring. (**C**) MYC Co-IP experiments with TMED7, TMED7ΔC-tail, or TMED7 C-tail mutant (charged residues mutated to alanines) probing for pull-down of GRASP55. N = 3 biological replicates. Right, lysate inputs. (**D**) MYC Co-IP experiment with chimeric domain-swapped constructs probing for pull-down of GRASP55. Top, TMED7 domains in blue; TMED5 domains in red. Black arrowheads denote CC-domain; Perforated box shows Co-IP results with TMED7 CC domain, which promotes the interaction with GRASP55. ß-actin as loading control. Right, lysate inputs. N = 3 biological replicates. (**E**) IF images of endogenous MUC1-fs (green) in stable TMED7-depleted P cells transduced with doxycycline-inducible TMED7-MYC or TMED7-TMED5CC chimera (red). Scale = 20 μm. (**F**) Quantification of (E). Log_2_ fold change of MUC1-fs intensity compared to empty vector. Mean ± SEM. N = 3 biological replicates. 5,000 - 10,000 cells analyzed per condition per replicate. * = p < 0.05.

To validate the exact binding site, we created a mutant (Figure 5C) that would abrogate salt-bridges, as determined by the MD simulation. We mutated all positively charged amino acids (Lysine (K), Arginine (R)) in the C-tail to alanine (A) and found that this AAAA mutant was unable to pull-down GRASP55 (Figure 5C), indicating that the salt-bridges indeed mediated the interaction between TMED7 C-tail and GRASP55 PDZ2 domains.

To probe the TMED7-GRASP55 interaction further, we generated TMED7-TMED5 chimeras across all major domains (C-tail, transmembrane (TM), coiled-coil (CC) and GOLD domains; Figure S6A, Figure 5D). Surprisingly, a TMED7 chimera containing the TMED5 C-tail domain was sufficient for binding GRASP55 (Figure 5D), suggesting that another TMED7 domain might be contributing to the interaction with GRASP55. Systematic exploration via domain swapping revealed that the TMED7 CC domain was critical for the GRASP55 interaction (Figure 5D, arrowheads). Chimeras missing the TMED7 CC domain showed little, if any, binding to GRASP55 (Figure 5D). Independently, IP-MS experiments with either wild-type TMED7 or TMED7-TMED5 CC chimeras, confirmed that removal of the TMED7 CC domain led to loss of interaction with GRASP55 (and other Golgi proteins such as Golgin-45)(Figure S6J-K and Table S6). As a final test, we transduced either wild-type TMED7 or TMED7-TMED5 CC chimeras into TMED7-depleted P cells. While wild-type TMED7 led to re-accumulation of mutant cargo, a TMED7 construct containing the TMED5 CC domain failed to do so (Figure 5E-F, Figure S6L). Because the CC domain has been implicated in TMED hetero-oligomer formation that facilitates binding to GRASP55^51^, we next tested whether the absence of the TMED7 CC in the TMED7-TMED5 CC chimera would abrogate its binding to the other three entrapment TMEDs. Surprisingly, the TMED7 chimera with a TMED5 CC was still able to interact with the other three entrapment TMEDs 2, 9 and 10 (Figure S6M), even though we confirmed that it had lost its ability to bind to GRASP55 (Figure S6M). This experiment indicated that GRASP55’s interaction with the other three entrapment TMEDs (Figure S6D) was critically dependent on an intact TMED7; TMED2, 9 or 10 could not compensate for the absence of the TMED7 CC domain and restore binding to GRASP55 (Figure S6M). The two most likely explanations for these findings - namely that both the TMED7 tail and CC domains contribute to GRASP55 binding - are either that (a) the TMED7 CC domain allosterically enhances the tail’s binding affinity for GRASP55, or (b) in addition to the other three TMEDs, one or more protein interactor(s) bind to the TMED7 CC domain thereby facilitating its interaction with GRASP55. Indeed, wild-type TMED7 has several interacting partners distinct from the TMED7-5CC chimera (Figure S6J). While additional future studies may parse through these possibilities, our *in silico*, biochemical and cell biological/functional studies have elucidated the molecular underpinnings of the TMED7-GRASP55 interaction for cargo entrapment by the hetero-oligomeric TMED complex.

### *In vivo*, inducible TMED7 deletion rescues proteinopathies in the kidney and the eye without on-target side effects

Our detailed mechanistic studies on TMED7 showed that TMED7 was specifically required for GRASP55-mediated Golgi entrapment of mutant cargo (Figure 4 and 5). To validate these studies, we tested the efficacy and safety of targeting TMED7 *in vivo*. To this end, we generated a new mouse model with tamoxifen-inducible, actin Cre-mediated deletion of TMED7. In these studies, inducible whole-body deletion of TMED7 for up to 23 weeks (from 12 weeks to 35 weeks of life) was well tolerated; TMED7 KOs were grossly indistinguishable from their littermate controls. We first assessed this TMED7 knockout model by crossing it into a well-established humanized knock-in mouse model of a kidney proteinopathy^17^ (*Tmed7^fl/fl^*, *CAGGCre–ER* mice, *Muc1^fs/+^,* Figure 6A). Mice with the Cre-driver (TMED7 KO) or without (CTRL) were injected with tamoxifen and then aged to assess MUC1-fs removal as well as tubular dilations and fibrosis (Figure 6B, Figure S7A-G). TMED7 KO mice displayed efficient removal of both TMED7 and the mutant cargo MUC1-fs (Figure 6C-E). While TMED2, 9 and 10 levels were slightly reduced, they were much less affected than TMED7 (Figure 6C), emphasizing the importance of TMED7 for MUC1-fs entrapment in-vivo. At 9 months, CTRL animals displayed severe tubular dilations (Figure 6F-G), tubulointerstitial fibrosis (Figure 6H) and infiltration of immune cells (Figure 6I), all of which were mitigated in TMED7 KO animals.

**Figure 6.**
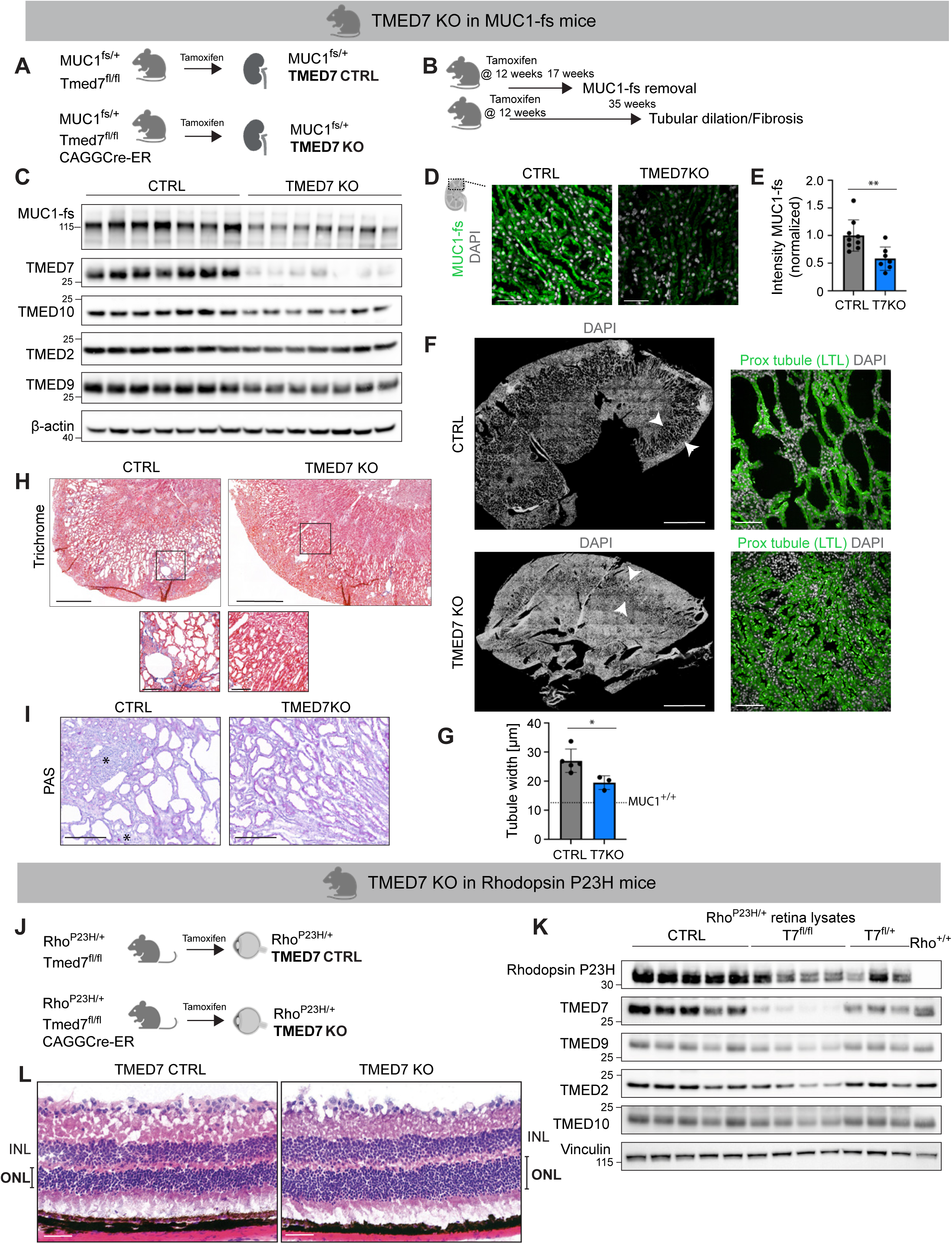
*In vivo*, inducible global deletion of TMED7 protects mice from kidney and eye disease. (**A**) Schematic of mouse model. *MUC1^fs/+^* mice were crossed with TMED7*^fl/fl^*mice either expressing CAGGCre-ER (TMED7 KO) or not (TMED7 CTRL). Inducible whole body Cre expression and subsequent genetic knockout was induced with Tamoxifen (3 doses of 125 mg/kg). (**B**) Tamoxifen was injected at 12 weeks of life, and kidneys were harvested at 17 weeks to assess MUC1-fs removal and at 35 weeks to assess tubular dilation and fibrosis. (**C**) Western blot of kidney lysates (17-week old) to assess MUC1-fs removal, TMED7 depletion and the abundance of other entrapment TMEDs. Loading controls, ß-actin. Each lane represents an animal. (**D**) Representative images of MUC1-fs (green) in mouse kidney cortical cross sections from TMED7 KO and CTRL 17 week-old female *Muc1^fs/+^*mice. N = 7-9 mice per group. Each dot represents an animal. Scale = 50 μm. (**E**) Analysis of (D) normalized to CTRL. Means ± SD. ** = p < 0.01. (**F**) Representative overview image of CTRL and TMED7 KO kidney cortical tissue sections (DAPI, left). Tubular dilation is noticeable especially in the outer medulla (area between white arrow heads) of CTRL animals. Scale = 2 mm. Insets of same kidneys showing DAPI (gray) and LTL (green, marking kidney proximal tubules) staining. Scale = 100 μm. (**G**) Quantification of (F) showing kidney tubule inner width. For comparison, the equivalent tubular width in control wild-type *Muc1^+/+^* mice is indicated by the perforated line. N = 3-5 mice per group. Each dot is an individual mouse. Means ± SD. * = p < 0.05. (**H**) Masson’s Trichrome stain of CTRL and TMED7 KO kidney tissue. Blue staining indicates fibrosis, notable in CTRL mice with prominent tubular dilations visible. N = 3 mice per group. Scale = 1 mm. Inset, higher magnification. Scale = 200 μm. (**I**) PAS staining for CTRL and TMED7 KO kidney tissue. Asterix indicates inflammatory/immune cell infiltration. N = 3 mice per group. Scale = 200 μm. (**J**) Schematic of mouse model. *Rho^P23H/+^* mice were crossed with TMED7*^fl/fl^*mice either expressing CAGGCre-ER (TMED7 KO) or not (TMED7 CTRL). Inducible whole body Cre expression and subsequent genetic knockout was induced with Tamoxifen (3 doses of 125 mg/kg). (**K**) Western blot of retina lysates to assess RHO-P23H removal, TMED7 depletion and the abundance of other entrapment TMEDs from TMED7 KO (homozygous and heterozygous) or CTRL mice. New mutant-specific P23H antibody is specific, as no signal was seen in Rho^+/+^ mice. Loading control, vinculin. Each lane represents one animal. (**L**) H&E staining of retina tissue showing preserved ONL thickness in P23H TMED7 KO vs. CTRL P23H animals. ONL = outer nuclear layer, INL = inner nuclear layer. Scale = 50 μm.

Next, we assessed the effect of deleting TMED7 *in vivo* in a mouse model of degenerative eye disease (RP due to rhodopsin P23H accumulation in the retina; Figure 6J). To assess allelic dose-dependence, we generated both heterozygous and homozygous TMED7 floxed animals. Tamoxifen induction led to a decrease in TMED7 proportional to the number of floxed alleles (Figure 6K) while TMED 2, 9 and 10 abundance was relatively unperturbed (Figure 6K). Excitingly, mutant RHO P23H cargo was similarly reduced in both genotypes (Figure 6K) implying that partial reduction in TMED7 might be sufficient to confer significant reduction in pathogenic misfolded rhodopsin burden. Hematoxylin & Eosin (H&E) staining of retina sections showed preserved ONL thickness in TMED7 KO retinas compared to control mice with RP (Figure 6L), a finding that not only illustrated the efficacy of targeting TMED7 for disease amelioration, but also reinforced the safety of this approach for preserving the overall structure of the retina. Taken together, these studies showed that global, inducible, long-term TMED7 deletion was tolerated and efficacious in genetically defined proteinopathies of the kidney and the eye.

### *In vivo,* oral administration of the small molecule BRD7635 induces clearance of mutant proteins and mitigates disease progression in proteinopathies of the kidney and the eye

As a final test, aiming to establish a therapeutic proof of concept, we probed the *in vivo* efficacy of BRD7635 which was administered to animals by oral gavage for up to 3 months of treatment. In the humanized MUC1-fs knock-in mouse model^17^, BRD7635 was effective at depleting the target TMEDs and their entrapped misfolded cargo (Figure 7A-B; drug pharmacokinetic (PK) data, Figure S7H). The drug dose-dependently decreased mutant cargo (MUC1-fs) accumulation (Figure 7B; Figure S7I-J). Since the hallmark of this disease is pathologic kidney tubular dilations (Figure S7A-G for natural history and technical details), we measured the diameter of kidney tubules^17^ in tissue from female *Muc1^fs/+^* mice after daily oral gavage with BRD7635. The luminal diameter of tubules in BRD7635-treated knock-in animals was similar to wild-type controls, suggesting that BRD7635 protected against pathologic kidney damage in this humanized proteinopathy model (Figure 7C-D, Figure S7A-G).

**Figure 7.**
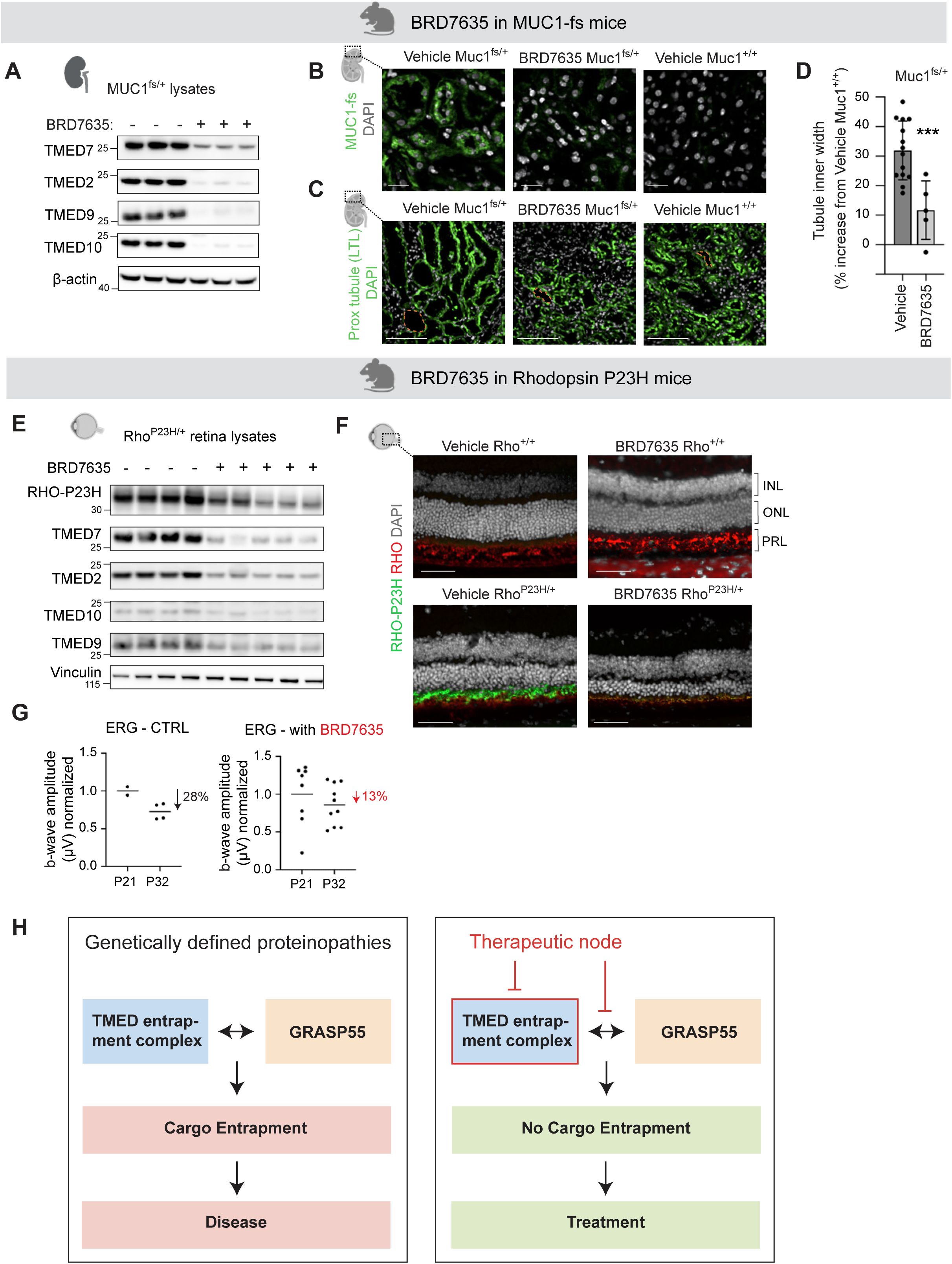
*In vivo*, BRD7635 efficiently removes toxic mutant protein cargoes in the kidney and the eye. (**A**) Target engagement confirmation by Western blot of TMEDs in whole kidney lysates from 4 month-old male *Muc1^fs/+^*mice following 5-day treatment with BRD7635 (3.0 mg/kg). β-actin as loading control. Each lane is an individual mouse. (**B**) Representative images of MUC1-fs (green) in mouse kidney cortical cross sections from 4 month-old male *Muc1^fs/+^*and *Muc1^+/+^* mice following 7-day BRD7635 or vehicle treatment. N = 3-5 mice per group. Scale = 20 μm. (**C**) Representative images of proximal tubules (LTL, green) in mouse kidney cross sections from 6 month-old female *Muc1^fs/+^* and *Muc1^+/+^* mice following 3 month treatment with BRD7635 (1.0 mg/kg) or vehicle. Examples of tubule inner width quantification regions (orange dashed lines) are indicated. Scale = 100 μm. (**D**) Quantification of (C) as % increase in kidney tubule inner width compared to vehicle-treated wild-type control *Muc1^+/+^* mice. N = 5-14 mice per group. Each dot is an individual mouse. Means ± SD. *** = p < 0.001 vehicle vs. BRD7635. (**E**) Western blot of TMEDs in retina lysates from P32 *Rho^P23H/+^* mice treated for 20 days with BRD7635 (1.0 mg/kg). Vinculin as loading control. Each lane represents 1 animal. (**F**) IF images of RHO-P23H (green) and RHO-WT (red) in mouse retina cross sections from 4 month-old *Rho^P23H/+^* and *Rho^+/+^* mice following 3 month treatment with vehicle, or BRD7635 (3.0 then 1.0 mg/kg for 1.5-months each). INL, inner nuclear layer; ONL, outer nuclear layer; PRL, photoreceptor layer. Scale = 25 μm. (**G**) ERG (0.5 cd/cm^2^) b-wave amplitude at P32 in CTRL and BRD7635 treated animals (1 mg/kg for 20 days) normalized to readings at P21. (**H**) Schematic overview of role of TMED entrapment complex in genetically defined proteinopathies (left) and potential treatment strategies (right) to prevent cargo entrapment and ameliorate disease. Future therapies may focus on clinically-applicable disruptors of the TMED entrapment complex and/or its interaction with GRASP55.

Turning our attention to retinitis pigmentosa, we asked whether BRD7635 could remove mutant rhodopsin P23H and whether this would result in improved retinal function. We probed mouse retina lysates from P32 *Rho^P23H/+^* mice following daily oral treatment with BRD7635. All entrapment TMEDs were reduced after drug treatment (Figure 7E). Next, using a newly developed P23H-specific antibody, both Western blot and immunofluorescence imaging indicated that BRD7635 treatment significantly decreased mutant RHO-P23H in retinas of *Rho^P23H/+^* mice as compared to control mice (Figure 7E-F). Similar to our results with wild-type semaphorin, wild-type rhodopsin (*Rho^+/+^* mice) was unaffected by drug treatment (Figure 7F). The b-waves–an ERG readout for photoreceptor activity in dark-adapted mice^43^–declined more slowly between P21 to P32 in *Rho^P23H/+^* mice treated with BRD7635 than those of controls (Figure 7G). These results showed that BRD7635 treatment reduced the relative risk for progression to blindness by approximately 50% (13% decline in BRD7635 treated versus 28% decline in untreated controls; [control-treatment] / [control] = [28-13]/28 = 53.5% relative risk reduction). However, to obtain these technically challenging ERG measurements in the face of rapidly advancing blindness in the *Rho^P23H/+^* mouse model, we had to begin drug dosing as early as postnatal day 12 (P12). Drug dosing in these young mice resulted in weight loss (not observed with treatment in older mice), a sign of toxicity, suggesting that the therapeutic window afforded by BRD7635 is narrow (i.e. narrow therapeutic index, the difference between efficacious versus toxic dose). Nevertheless, these proof-of-concept studies showed that targeting TMEDs with BRD7635 could mitigate progression to kidney disease and blindness in mice, overall validating the druggability of the entrapment node.

## Discussion

This study uncovered a new paradigm in cellular proteostasis whereby a specific hetero-oligomeric cargo receptor complex entrapped several degenerative disease-causing proteins that escaped the ER and entered a post-ER secretory pathway. To open up this new scientific direction, we harnessed a wide array of tools including CRISPR screens, proteomics, Alpha-Fold-enabled molecular dynamics modeling, biochemical assays and *in vivo* disease modeling with functional readouts including electroretinograms (ERGs) in living mice. We thus made several mechanistic and therapeutically important discoveries.

First, we found that TMED7 formed a hetero-oligomeric complex with TMED 2, 9 and 10 (entrapment TMEDs) to mediate the entrapment of many topologically distinct mutant cargoes. We further identified the precise molecular mechanism for entrapment and cargo retention through TMED7’s specific interaction with GRASP55 (Figure 7H), an integral Golgi protein. We propose that cells deploy the TMED entrapment complex to achieve proteostasis when faced with the challenge of handling misfolded cargoes that evade proteasomal degradation and manage to enter a post-ER secretory pathway. Under normal, homeostatic conditions, stochastic misfolding of proteins (i.e. not due to genetic mutations) could be corrected by a return to the ER for another opportunity to have them properly folded^52^. However, when misfolded cargoes are continuously produced as the result of a genetic mutation, the TMED entrapment mechanism may become maladaptive, leading to detrimental over-accumulation of mutant cargo that overwhelms the cell’s capacity to degrade it. This would explain why wild-type proteins in our experiments were unperturbed when we targeted TMEDs - they are not primary clients for the entrapment complex.

Second, our *in vivo* studies with inducible TMED7 deletion in the kidney and the eye (a) confirmed the long term on-target safety of a putative entrapment complex targeted therapy (even homozygous TMED7 deletion in the kidney over >23 weeks appeared to be safe), and (b) revealed a new therapeutic opportunity to target TMED7 with a nucleic acid modality such as siRNA or anti-sense oligonucleotides (ASO). There may be several advantages to this approach, especially in the eye. For example, hereditary blindness due to rhodopsin mutations alone can be caused by more than a hundred distinct missense mutations^38^, making gene therapy for each of these mutations profoundly impractical and prohibitively expensive. Targeting TMED7 (e.g. with direct intravitreal injection of an ASO^53^) offers a clear advantage: it would allow us to potentially address many of the 100 rhodopsin disease-causing mutations with a single therapy delivered specifically to the eye.

Third, our work on BRD7635 serves as the foundational proof-of-concept for a future drug development program that could benefit patients with many genetically and clinically distinct proteinopathies. *In vivo*, targeting the entrapment complex worked efficiently to reduce or abrogate the accumulation of many structurally and functionally distinct cargoes (i.e. glycoprotein, GPCR, single transmembrane domain protein) expressed in different cell types and organs (i.e. eye, kidney). In our studies, BRD7635 showed no off-target effects *in vitro* across a wide array of putative targets (Table S3). Prolonged drug treatment (for more than 3 months) in our kidney studies was well tolerated in mice. In the case of retinitis pigmentosa, dosing young animals (as young as P12) showed that BRD7635 has a narrower therapeutic window, thus pointing to the need for future medicinal chemistry work to produce improved, clinically applicable TMED-targeted small molecules. Nevertheless, the *in vivo* efficacy of BRD7635 in these proof-of-concept studies highlights the salutary potential of this approach to treat multiple currently incurable proteinopathies. Further confidence in the TMED complex as a therapeutic target was gained from our studies of inducible whole-body deletion of TMED9 and TMED7, especially the latter, which was globally ablated for up to 23 weeks in mice without any toxicities or side-effects. In total, we estimate that more than 50 different genes, each harboring dozens to hundreds of missense mutations, result in post-ER proteinopathies affecting different organs, many of which may be ultimately addressed by targeting the TMED entrapment node (Figure 7H). Consequently, if successful, a TMED-targeted treatment could help millions of patients around the world suffering from these devastating diseases.

In conclusion, our work revealed a convergent node for multiple clinically distinct diseases, thereby serving as a foundational example of a new concept we call nodal biology^54^. The key next question rooted in our work is how many more convergent nodes remain to be discovered. Searching for shared nodes downstream of distinct human disease-causing mutations could augment the scale and speed at which we approach our mechanistic understanding and treatment of genetic diseases. Targeting these convergent nodes could catalyze an unprecedented acceleration in the development of therapies for devastating diseases for the benefit of larger patient populations.

## Methods

### Buffers

(Co-IP) Lysis Buffer: 100 mM NaCl, 5 mM EDTA, 50 mM Tris pH 7.5, 1% NP-40, 1 tablet each of freshly added PhosSTOP (Roche, 0490683700) and complete EDTA-free Protease Inhibitor Cocktail (Roche, 04693159001) per 10 ml buffer, pH 7.5 at 4 °C

Elution buffer for Co-IP: 2X NuPAGE LDS Sample Buffer (Thermo Scientific, NP0008), 1X NuPage Sample Reducing Agent (Thermo Scientific, NP0009), and nuclease free water (Invitrogen, AM9937)

5x TBST buffer: 750 mM NaCl, 250 mM Tris pH 7.5, 0.25% Tween-20 (GFP IP) or 0.05% Tween-20 (MYC IP)

PBS-T: PBS, pH7.4 with 0.01% Tween-20

Antigen Retrieval Buffer: 3 M urea, 3 M guanidinium chloride, 0.5 M L-glycine, 70 mM tris(2-carboxyethyl)phosphine (TCEP) [Sigma, C4706], adjusted to pH 2.5 using HCl

Blocking solution 1 for IF: 100 mM Tris HCL pH 8; 150 mM NaCl; 5 g/L Blocking Reagent (Roche,11096176001)

Blocking solution 2 for IF: LICOR Intercept (PBS) Blocking Buffer (LICORbio, 927-70001) with 146 mM of malemide (Sigma-Aldrich,129585) per 5 ml of buffer

Imaging buffer: 700 mM NAC in MilliQ water, pH 7.4

### Mouse generation

MUC1-fs knock-in *MUC1^fs/+^* 129S2 mice (*MUC1-fs*) were previously generated by GenOway (Lyon, France) using embryonic stem cells genetically modified by homologous recombination to knock-in an ADTKD mutant human *MUC1* gene into the murine *MUC1* locus^17^. Mutant rhodopsin *Rho^P23H/+^* mice were ordered from The Jackson Laboratory (RRID:IMSR_JAX:017628). Conditional Tmed9 knockout *Tmed9^fl/fl^* C57BL/6J mice were generated by Taconic Biosciences GmbH (Köln, Germany) by CRISPR/Cas-mediated genome engineering, with conditional knockout of exons 3-5 of the murine *Tmed9* gene. Conditional Tmed7 knockout TMED7fl/fl C57BL/6J mice were generated by Brown Transgenic Facility (Providence, Rhode Island) by CRISPR/Cas-mediated genome engineering, with conditional knockout of exons 2-3 of the murine *Tmed7* gene. CAGGCre-ER mice were obtained from the Jackson Laboratory (RRID:IMSR_JAX:004682) and crossed with *Tmed9^fl/fl^ Rho^P23H/+^*, *Tmed7^fl/fl^ Rho^P23H/+^*and *Tmed7^fl/fl^ MUC1^fs/+^* mice.

Mice were maintained on a 12 h light/dark cycle at 18–26 °C in an AAALAC accredited facility and fed ad libitum with water and PicoLab Rodent Diet 20 pellets (LabDiet). Male and female mice ranging in age from 4-9 months were used for experimentation, except for Rho*^P23H/+^* mice which were 3-7 weeks old. All animal experiments were approved by the Institutional Animal Care and Use Committee (IACUC) at The Broad Institute of MIT and Harvard and were conducted in accordance with National Institutes of Health (NIH) animal research guidelines.

### Cell culture

HEK293T cells (ATCC) and Rho GFP HEK293T cells were cultured according to standard protocol at 37 °C with 5% CO_2_ in Dulbecco’s Modified Eagle’s Medium (DMEM), high glucose, GlutaMAX^TM^ (Gibco, 10569-010) supplemented with 10% Fetal Bovine Serum (Invitrogen, A5670701) and 1% Penicillin/Streptomycin (Life Technologies, 15140-122).

ADTKD/MKD-MUC1 patient immortalized kidney epithelial cells (P cells) and healthy control immortalized kidney epithelial cells (N cells)^17^ were maintained at 37 °C with 5% CO_2_ in RenaLife Renal Basal Medium (LM-0010) supplemented with RenaLife LifeFactors (Lifeline Cell Technology, LS-1048), with the exclusion of gentamicin and amphotericin B. Cas9-positive P cells were selected with media supplemented with 8 µg/mL blasticidin S (Gibco, A1113903) and CRISPR/Cas9 stable depletion P cells were selected in media supplemented with 4 µg/mL puromycin (Gibco, A1113803).

### CRISPR/Cas9 generation of TMED stable depletion P cells

To generate Cas9-expressing P cells, the Cas9 expression vector pXPR_BRD111 was used (Broad Institute Genetic Perturbation Platform). Stable Cas9-positive P cells were plated at 670,000 cells/well in 12-well plates in RenaLife media containing 4 µg/mL protamine sulfate. 120 µL/well gRNA virus (Broad Institute Genetic Perturbation Platform) was added per target separately to freshly plated cells. Plates were then centrifuged at 930 x g for 1 h at 30 °C and incubated overnight at 37 °C. The next day, media was removed, cells were washed once with PBS, and selection media (containing 4 µg/mL puromycin) was added. The following gRNAs were used to generate individual stable depletion cell lines: BRDN0003462535 (gTMED1-2), BRDN0003230309 (gTMED1-1), BRDN0003581071 (gTMED10-1), BRDN0003230535 (gTMED10-2), BRDN0003230208 (gTMED2-1), BRDN0003230396 (gTMED2-2), BRDN0003492798 (gTMED5-1), BRDN0003482993 (gTMED5-2), BRDN0003481231 (gTMED7-1), BRDN0003482908 (gTMED7-2), BRDN0003481199 (gTMED9-2), BRDN0003481863 (gTMED9-1), BRDN0001146753 (gNTC-1), BRDN0001146709 (gNTC-2).

### GFP-Rhodopsin and GFP-Semaphorin 4A Cell Line generation

We created HEK293T cell lines expressing Rhodopsin WT, Rhodopsin P23H, Rhodopsin Q344X, Rhodopsin T17M, Rhodopsin R135W, Semaphorin 4A WT and Semaphorin 4A F350C. Plasmids were designed by Vectorbuilder with a 2 vector system. One vector contains a reverse tetracycline-controlled activator (rtTA, Hygromycin B resistant), the other one contains the gene of interest tagged with EGFP (Puromycin resistant). First, 3×10^5^ HEK293T cells were plated into a 6 well plate and transduced with 1 MOI of pLV[Exp]-CMV>tTS/rtTA/Hygro (rtTA) virus in 1 mL of complete DMEM+Glutamax media the next day. Virus was removed after 24 h and cells were subjected to selection in 200 µg/mL Hygromycin B (Thermo Fisher Scientific, 10687010).

Surviving rtTA HEK293T cells were plated at 3×10^5^ cells per well of a 6 well plate and transduced the next day with 1 MOI of virus in 1 mL of media for 24 h. Cells were selected in 200 µg/mL Hygromycin and 1.2 µg/mL Puromycin Dihydrochloride (Life Technologies, A1113803). All GFP-tagged RHO and SEMA4A HEK293T lines were expanded. For all these lines, gene expression was induced by 1 µg/mL doxycycline hyclate (Sigma-Aldrich, D9891) for at least 2 days. Separate rtTA induction was found to be unnecessary.

### Design and generation of plasmid constructs

For tagged TMED constructs, epitope tags were added after the signal peptide in the N-terminal region, and gene fragments were synthesized by IDT. TMED and MUC1-fs fragments were then cloned into the pcDNA™3.1(+) Mammalian Expression Vector (Thermo Fisher Scientific #V79020) and transformed into *E. coli*. MUC1-wt constructs were designed and purchased separately from VectorBuilder.

Chimera constructs for TMED5 and 7 were designed by identifying the C-tail, Transmembrane, CC and GOLD domains using existing Uniprot annotations and making in-frame swaps of these domains, with a total of 10 mutant versions of TMED5 and 7 being generated. Double-stranded gene fragments of the chimeric sequences were ordered from IDT or Twist Biosciences and placed into two different vectors. For co-immuniprecipitation experiments, the gene fragments were cloned into the pcDNA™3.1(+) Mammalian Expression Vector (Thermo Fisher Scientific #V79020) using the NEBuilder® HiFi DNA Assembly Cloning Kit (New England Biolabs #E5520) in NEB 5-alpha High Efficiency Competent *E. coli*. For rescue experiments, the gene fragments were synthesized in an Ampicillin-resistant Entry clone vector, which were then cloned into the pInducer20 lentiviral vector (Addgene #44012) via a two-step Gateway recombination cloning protocol with BP and LR Clonase enzymes (Thermo Fisher Scientific #11791020, #11789020) to allow for doxycycline-inducible conditional gene expression.

In all cases, transformants were screened via full plasmid sequencing by Primordium Labs. Maxiprepped constructs were additionally sequence verified prior to use in downstream experiments.

### Transfection

For overexpression Co-IP, HEK293T cells were plated in 10-cm or 15-cm dishes and the next day transfected with 4.86 µg/plate or 10-40 µg/plate of total plasmid DNA in a 2:1 ratio with Lipofectamine 2000 (Invitrogen, 11668019) in Opti-MEM I Reduced Serum Medium (Gibco, 31985088) and incubated for 24 h before further processing. For localization of TMED7-MYC, TMED1-MYC, and TMED5-MYC, P cells were plated in 96-well PhenoPlates (Revvity, 6055308) at 12,500 cells/well and transfected the next day with 0.1 µg/well of total plasmid DNA, as described above.

### Lentivirus packaging

HEK293T cells cultured in complete HEK media without antibiotics at 37 °C and 5% CO_2_ were used to produce lentivirus particles. Three plasmids (pCMV Δ 8.91, pMDG2 VSV-G, and a cargo containing plasmid) were co-transfected into HEK293T cells with Lipofectamine 3000 (Invitrogen, L3000001) or TransIT®-LT1 (Mirus Biom, 2304). After 18 h, culture medium was replaced with harvest media, DMEM GlutaMAX^TM^ supplemented with 30% FBS. Media was harvested after 24 and 48 h, and passed through a 0.45 μm filter. The lentiviral particles were concentrated using Lenti-X concentrator (Takara Bio, 631232). The collected lentiviral particles were stored at −80 °C.

### Acute CRISPR/Cas9 depletion of TMEDs in stably depleted P cells

Stable depletion P cell lines were plated at 8,000 cells/well in 384-well PhenoPlate microplates (Revvity) in 50 µL media and immediately treated with 8 µL/well gRNA virus (Broad Institute Genetic Perturbation Platform) and incubated at 37 °C overnight. The next day, media were aspirated and cells were washed twice with PBS, then fed with 50 μL fresh media and incubated at 37 °C for seven days before processing for immunofluorescence staining. The following gRNA stable TMED depletion P cell lines were used: BRDN0003482993 (gTMED5-2), BRDN0003481199 (gTMED9-2), BRDN0001146709 (gNTC-2). The following gRNAs were used for acute TMED depletion: BRDN0003462535 (gTMED1-2), BRDN0003230535 (gTMED10-2), BRDN0003230208 (gTMED2-1), BRDN0003492798 (gTMED5-1), BRDN0003481231 (gTMED7-1), BRDN0003481199 (gTMED9-2), BRDN0001146753 (gNTC-1), BRDN0001146709 (gNTC-2).

### Acute knockdown of TMEDs in RHO P23H HEK cells

RHO P23H HEK cells (500,000/well) were plated in a 6 well plate for 24 h prior to Edit-R all in one TMED5, 7, 9 virus being added for 18-24 h in media containing Polybrene (8 μg/mL). EditR-all-in-one plasmids contain the gRNA and the Cas9 nuclease in the same vector. Our Edit-R-all-in-one plasmids contained the pXPR_023 backbone (GPP Broad Institute) and the same guide as used for making stable P cell lines: gNTC (BRDN0001477947), gTMED5 (BRDN0008030436), gTMED7 (BRDN0008030427), gTMED9 (BRDN0008030391). After infection, cells were incubated in selection media with Blasticidin (3 μg/mL) for 4 days. Then cells were counted and plated (20,000/well) into 384-well imaging plates for 4 days in media containing 500 ng/mL doxycycline for induction of P23H-GFP followed by fixation with 4% PFA/PBS. Hoechst was added for 15 min prior to imaging of GFP in the 488 channel.

### Acute knockdown of GRASP55 in P cells

P cells (300,000 cells) were grown in a 6 well plate. After 24h, media was exchanged to media containing 8 μg/mL (Millipore Sigma) Polybrene. Lentivirus particles of NTC shRNA (Millipore Sigma, SHC216) and GRASP55 shRNA (Millipore Sigma, TRC Clone ID: TRCN0000278363) were added for 18-24 h followed by replacement with normal P cell media. After 48 h, cells were plated into 384-well plates (30,000 cells/well) for 72 h prior to fixation and immunofluorescence imaging.

### Lentiviral transduction of TMED9-FLAG, TMED7-MYC or TMED7-5CC-MYC in TMED9 and TMED7 stable depletion P cells

P cells were cultured overnight to 70% confluency in a 6-well dish. The collected lentiviral particles supplemented with 4 μg/mL protamine sulfate were added to the prepared P cells. After 24 h of infection, the supernatant was replaced with fresh medium containing 100 μg/mL G418 and incubated for 3 days. After successful selection, P cells were seeded at a density of 14,000 cells/well in 384-well PhenoPlate microplates (Revvity) pre-coated with 0.25 mg/mL Synthemax II SC Substrate (Corning). G418-selected cells were cultivated in the absence or presence of 500 ng/mL (TMED9-FLAG) or 1000 ng/mL (TMED7-MYC, TMED7-5CC-MYC) doxycycline for 96 h (TMED9-FLAG) or 6 days (TMED7-MYC, TMED7-5CC-MYC) to induce the expression of WT TMED9-FLAG and TMED7-MYC, followed by immunofluorescence.

### Lysate preparation

Cells were washed in ice-cold PBS and then lysed in Co-IP lysis buffer containing protease inhibitors (Roche) and phosphatase inhibitors. Lysates were incubated for 30 minutes at 4 °C while rocking then centrifuged for 10 minutes at 16000 g at 4 °C. Supernatants were isolated and protein quantified using the Pierce BCA Assay Kit (Thermo Fisher Scientific, 23225). 1x NuPAGE LDS loading buffer and 1x NuPAGE reducing agent with nuclease-free water were added and samples were heated to 75 °C for 10 min (note that retina lysates probed for Rhodopsin were not heated).

### Co-immunoprecipitation

FLAG Co-IPs were performed using 25 µl Anti-FLAG M2 Magnetic Beads (Sigma-Aldrich, M8823) per IP. Beads were washed twice with 1 mL TBS (150 mM NaCl, 50 mM Tris, pH 7.5) and resuspended in the original bead volume of Co-IP lysis buffer. One mg of sample lysate protein was added and rotated at 4 °C overnight. The next day samples were washed twice with 1 mL Co-IP lysis buffer and twice with 1 mL TBS before resuspension in elution buffer. Samples were then heated at 75 °C for 10 min and separated on a magnetic rack. Eluate was collected for further analysis.

MYC Co-IPs were performed using 25-50 µL of Pierce anti-c-MYC Magnetic Beads (Thermo Scientific, 88843) per IP. Beads were washed twice with 1 mL Co-IP lysis buffer and resuspended in the original bead volume of Co-IP lysis buffer. One to 2 mg of sample lysate protein was added and rotated at 24 °C for 30 minutes. Samples were washed three times with 1 mL 5X TBST and once with 1 mL PBS or water before resuspension in elution buffer. Samples were then heated at 75 °C for 10 min and separated on a magnetic rack. Eluate was collected for further analysis.

GFP Co-IP was performed using 50 mL of GFP-Trap Magnetic Agarose Beads (ChromoTek, gtma-100) per sample. Beads were washed twice with 1 mL of 1x TBST before resuspending in the original bead volume of Lysis Buffer. Co-IP samples were prepared by adding 2 mg of protein and diluting to 300 µL with Co-IP Lysis Buffer, then adding 50 µL of GFP-Trap Beads. Samples were rotated for 1 h at 4 °C, then samples were washed thrice with 1 mL of 5x TBST and then once with 1 mL of MilliQ Water. All liquid was removed then samples were resuspended very lightly by vortexing in 30 µL of elution buffer. Samples were then heated at 55 °C for 10 min and beads were removed. 10 µL (Rhodopsin Co-IP) or 25 µL (Semaphorin Co-IP) of Co-IP samples were loaded.

Endogenous P cell Co-IPs were performed using 25 µL of Pierce Protein A/G Magnetic Beads (Thermo Scientific, 88803) per IP. The day prior to bead addition, 2-4 mg of sample protein lysate was added to 6-10 µg TMED9 antibody (Thermo Fisher #PA5-21422) and 6µg GORASP2 antibody (Grasp55, Proteintech #10598-1-AP) and then rotated at 4 °C overnight. The next day, beads were washed 3 times with 1 mL Co-IP lysis buffer and resuspended in the original bead volume of Co-IP lysis buffer, then added to samples and rotated 1 h at 4 °C. Samples were incubated in elution buffer with occasional vortexing for 10 min, then beads were removed and samples heated at 75 °C for 10 minutes. Further analysis was performed by Western blot. Input samples were prepared at 1 µg/µL.

### Western blot

For Western blot we used a SDS-PAGE gel electrophoresis system (Invitrogen™ XCell SureLock™ Mini-Cell Electrophoresis System) with either NuPAGE 3-8% Tris-Acetate gels with Tris-Acetate Running Buffer for MUC1-fs (Thermo Scientific) or NuPAGE 4-12% Bis-Tris gels with MES running buffer (Thermo Fisher Scientific) for all other proteins. Samples were loaded with Pageruler Preset or Plus (Thermo Fisher Scientific, Cat # 26616, 26619) for Bis-Tris gels and HiMark pre-stained protein standard (Thermo Fisher Scientific, Cat #LC5699) for Tris-acetate gels and run for 100 - 120 min at 120 V. Gels were transferred onto a Nitrocellulose membrane (Trans-BlotR TURBO filter pack, Bio-Rad) and transferred for 5-10 min using a Bio-Rad Trans-Blot TURBO transfer system followed by blocking for 1 h in 5% nonfat dry milk in PBS-T. Primary antibodies were added in 5% nonfat dry milk in PBS-T and incubated overnight. The next day, after three washes with PBS-T, the membranes were incubated with secondary antibody (1:2500, rabbit (Cell Signaling Technology, 7074S), mouse (Cell Signaling Technology, 7076P2)) for 1 h at room temperature, washed three more times in PBS-T, and incubated in SuperSignal West Femto (Thermo Fisher Scientific, 34096) or SuperSignal West Pico (Thermo Fisher Scientific, 34580). Immunoreactive bands were imaged in chemiluminescent and colorimetric channels by BioRad ChemiDoc. IP membranes were incubated (unless antibodies were conjugated with HRP) with Mouse or Rabbit Trueblot (1:1000, Rockland, Cat# 18-8817-33 and 18-8816-33) or VeriBlot HRP (1:200, Abcam, ab131366) for 1 h at RT.

Unless otherwise stated, the following primary antibodies were used at 1:1000 dilution: mouse monoclonal anti-GFP (Cell Signaling Technology, 2955S), mouse monoclonal anti-FLAG-HRP (Sigma-Aldrich, A8592), mouse monoclonal anti-myc-HRP (1:500, Santa Cruz Technologies, sc-40), rabbit anti-β-actin-HRP (Cell Signaling Technology, 5125S), rabbit monoclonal anti-GAPDH-HRP (Cell Signaling Technology, 3683S), monovalent Fab-V5H anti-MUC1-fs antibody (BioRad AbD22655.4, AbD22655.9, created using HuCAL® antibody generation against a neo-epitope in MUC1-fs), rabbit polyclonal anti-GORASP2 antibody (Proteintech, 10598-1-AP), rabbit polyclonal anti-BLZF1 (Golgin-45, GeneTex, GTX116434), rabbit anti-P23H rhodopsin antibody (Innovagen, made from a short synthetic peptide containing the mutation, for specificity see Figure 8B). For detection of MUC1-fs an anti V5 Tag-HRP Monoclonal Antibody (Invitrogen R96125) was used. For TMED antibodies please refer to Table S7.

### Co-immunoprecipitation for proteomics of TMED9-MYC

For each experimental condition, 8.1×10^6^ HEK293T cells were plated for transfection onto a 10 cm plate in complete media with three replicates per condition. The next day, cells were transfected with 9.72 µg of total plasmid DNA per condition (TMED9-MYC and FLAG-MUC1-fs or empty vector and FLAG-MUC1-fs) with Lipofectamine 2000 (Invitrogen, Cat: 11668-019, Lot: 2423710) in a 1:2 ratio.

To immunoprecipitate TMED9-MYC, cell lysates were prepared as above and 3 mg of lysate was combined with 37.5 μL of well-resuspended Myc-Trap® Magnetic Agarose Beads (Chromotek, ytma-20) adjusted with lysis buffer to 2 mL. Beads were first washed 3 times with 1 mL of lysis buffer and resuspended. Lysate and bead mixtures were rotated for 1 h at 4 °C, followed by 3 washes with lysis buffer and three times with wash buffer (10 mM Tris-HCl pH 7.5, 150 mM NaCl, 0.5 mM EDTA, pH 7.5 at 4 °C, with protease and phosphatase inhibitors). Volume equivalent to 2 mg of input protein was taken for mass spectrometry analysis (Broad Institute Proteomics Platform). The remaining protein was eluted in 25 µL of elution buffer followed by heating at 95 °C for 10 min. Eluted protein was saved for quality control western blot analysis.

### Proteomics sample processing for TMED9-MYC MS-IP

Proteins bound to antibody beads were washed 4x with 200 μL of 50 mM Tris-HCl (pH 7.5). The final wash was removed and beads were incubated with 80 μL of 2 M urea in 50 mM Tris-HCL containing 1 mM DTT and 0.4 μg trypsin at 25 °C for 1 h while shaking at 1,000 r.p.m. After 1 h, the supernatant was removed and transferred to fresh tubes. Beads were washed twice with 60 μL of 2 M urea in 50 mM Tris (pH 7.5) buffer and combined with the on-bead digest supernatant. Disulfide bonds were reduced with 4 mM DTT at 25 °C for 30 min with shaking at 1,000 r.p.m.

Sample eluent was alkylated with 10 mM iodoacetamide at 25 °C for 45 min in the dark while shaking at 1,000 r.p.m. Samples were digested with 0.5 μg of trypsin overnight at 25 °C with shaking at 700 r.p.m. Following overnight digestion, formic acid (FA) was added to eluents to ∼1% (vol/vol) and pH 3.

Peptides were desalted using C18 StageTips. Briefly, C18 StageTips were conditioned with 100 μL of 100% MeOH, 100 μL of 0.1% (vol/vol) FA and 50% (vol/vol) acetonitrile, and twice with 100 μL of 0.1% (vol/vol) FA. Acidified peptides were loaded onto the conditioned StageTips and washed twice with 100 μL of 0.1% (vol/vol) formic acid. Peptides were eluted from the StageTips with 50 μL 0.1% (vol/vol) FA and 50% (vol/vol) acetonitrile and vacuum centrifuged until completely dry.

For TMT labeling, peptides were reconstituted in 80 μL of 50 mM HEPES. Each sample was labeled with 20 μl of a specific 20 mg/mL TMT10 label for 1 h while shaking at 1000 r.p.m. TMT-labeling reactions were quenched with 4 μL of 5% (vol/vol) hydroxylamine at room temperature for 15 min with shaking and combined. Samples were dried and peptides were desalted on C18 StageTips as described above.

For each sample, 50% was fractionated by basic pH reversed-phase (bRP) fractionation. For bRP fractionation, a StageTip was prepared using two disks of SDB-XC material (Empore 2240), washed and equilibrated with 100% MeOH, followed by 50% ACN/1% FA and 0.1% FA. Dried peptides were resuspended in 3% FA/5% ACN and loaded onto the Stage-Tip. Peptides were elution in six fractions with increasing concentrations of ACN (5%, 10%, 15%, 20%, 25% and 45%) in 0.1% (w/v) NH4OH (28% NH3 [w/v]), pH 10. Fractions were dried down in a vacuum concentrator.

### Mass spectrometry data processing and analysis for TMED9 MS-IP

Desalted, TMT-labeled peptides were resuspended in 9 μL of 3% MeCN, 0.1% FA and analyzed by online nanoflow liquid chromatography tandem mass spectrometry (LC-MS/MS) using an Exploris 480 (Thermo Fisher Scientific) coupled on-line to a Proxeon Easy-nLC 1200 (Thermo Fisher Scientific). 4 μL of each sample was loaded at 500 nL/min onto a microcapillary column (360 μm outer diameter x 75 μm inner diameter) containing an integrated electrospray emitter tip (10 μm) packed to approximately 24 cm with ReproSil-Pur C18-AQ 1.9 μm beads (Dr. Maisch GmbH) and heated to 50 °C. The HPLC solvent A was 3% MeCN, 0.1% FA, and the solvent B was 90% MeCN, 0.1% FA. Peptides were eluted into the mass spectrometer at a flow rate of 200 nL/min. Non-fractionated samples were analyzed using a 154 min LC-MS/MS method with the following gradient profile: (min:%B) 0:2; 2:6; 120:35; 122:60; 130:90; 143:90; 144:50; 154:50 (the last two steps at 500 nL/min flow rate). The bRP fractions were run with 110-minute method, which used the following gradient profile: (min:%B) 0:2; 1:6; 85:30; 94:60; 95:90;100:90; 101:50; 110:50 (the last two steps at 500 nL/min flow rate). The Exploris 480 was operated in the data-dependent mode acquiring HCD MS/MS scans (r = 45,000) after each MS1 scan (r = 60,000) on the top 20 most abundant ions using an MS1 target of 3E6 and an MS2 target of 5E4. The maximum ion time utilized for MS/MS scans was 120 ms (single-shot) and 105 ms (bRP fractions); the HCD normalized collision energy was set to 32; the isolation window (m/z) = 0.7; the dynamic exclusion time was set to 20 s, and the peptide match and isotope exclusion functions were enabled. Fit filter was enabled at 50% using a purity window of 1.2.

Collected data were analyzed using the Spectrum Mill software package v6.1 pre-release (Agilent Technologies). Nearby MS scans with similar precursor m/z were merged if they were within ± 45 s retention time and ± 1.4 m/z tolerance. MS/MS spectra were excluded from searching if they failed the quality filter by lacking a sequence tag length >0 or did not have a precursor MH+ in the range of 600 - 6000. All extracted spectra were searched against a UniProt human database. Search parameters included: ESI QEXACTIVE-HCD-v2 scoring parent and fragment mass tolerance of 20 ppm, 30% minimum matched peak intensity, trypsin allow P enzyme specificity with up to four missed cleavages and calculate reversed database scores enabled. Fixed modifications were carbamidomethylation at cysteine. TMT labeling was required at lysine, but peptide N termini were allowed to be either labeled or unlabeled. Allowed variable modifications were protein N-terminal acetylation and oxidized methionine. Individual spectra were automatically assigned a confidence score using the Spectrum Mill auto-validation module. Score at the peptide mode was based on target-decoy false discovery rate (FDR) of 1%. A second round of validation was performed at the protein level, requiring a minimum protein score of 0. Relative abundances of proteins were determined using TMT reporter ion intensity ratios from each MS/MS spectrum, and the median ratio was calculated from all MS/MS spectra contributing to a protein subgroup. Proteins identified by 2 or more distinct peptides and ratio counts were considered for the dataset and analyzed as previously described^55^ and deposited in MassIVE(MSV000089757).

### Co-immunoprecipitation for proteomics of TMED5 and TMED7-MYC MS-IP (wild-type and coiled-coil swap)

HEK293T cells were plated in 15 cm dishes with 4 biological replicates per condition. Cells were transfected with plasmids (see Transfection) the following day and harvested after 36 h. Cells were washed 1x with ice-cold PBS, lysed in Co-IP buffer and lysates were prepared as mentioned above. Pierce anti c-Myc magnetic beads were vortexed and washed 3x with 2 mL of 1xTBS-T and resuspended to the original volume with Co-IP buffer. Per replicate, 5 mg of lysate was resuspended with 75 µL of beads and placed on the magnetic stand. Beads were washed twice with 5xTBST buffer followed by 3 washes with PBS. Volume equivalent of 0.75 mg input was taken out and eluted for a Western blot to ensure sample quality. The rest of the beads were flash frozen in liquid nitrogen and stored at −80 °C prior to transfer to the mass spectrometry facility at the Whitehead Institute. Protein-bound magnetic beads were reduced and alkylated at 70 °C for 10 min in 100 mM TEAB with 40 mM CAA and 10 mM TCEP. Trypsin/Lys-C mix was added 1:100 and bead suspensions were digested overnight at 37 °C in a shaking incubator at 115 RPM. The following day, an additional dose of trypsin/LysC mix was added 1:100 in 100 mM TEAB and the digestion proceeded at 37 °C for 4 h. Peptide digests were purified using SDB-RPS stage tips as described^56^. Peptides were then dried in a SpeedVac concentrator at 55 °C and resuspended in 0.2% (v/v) formic acid in MS-grade water to a final concentration of 0.2 µg/µL. The Pierce Quantitative Fluorometric Peptide Assay (Thermo Scientific, 23290) was used to determine peptide concentrations following the manufacturer’s instructions.

LC-MS/MS data were collected on an Orbitrap Eclipse mass spectrometer coupled with a Vanquish Neo nanoLC, a FAIMS Pro Interface, and an Easy Spray ESI source, all by Thermo Fisher Scientific (Waltham, MA, USA). For nanoLC separation, an Aurora Ultimate TS25 column (75 µm x 25 cm, 120 Å) from IonOpticks (Fitzroy, VIC, AUS) was utilized. Peptide extracts were injected with a volume of 2 µL. Peptides were separated using a mobile phase of 0.1% (v/v) formic acid in water (solution A) and 0.1% (v/v) formic acid in 80% (v/v) acetonitrile (solution B), flowing at a rate of 400 nL/min, while maintaining the column temperature at 50 °C. Initial column conditioning was performed with 3% solution B, followed by a linear gradient increasing to 25% B over 45 minutes, then to 40% B over 15 minutes, and finally to 95% B over 5 minutes. Remaining peptides on the C18 resin were eluted at 95% B for 6 minutes. In positive ion mode, the ion source temperature was set to 305 °C with a voltage of 2000 V, and ionized peptides were passed through the FAIMS Pro unit at −50 V. Mass spectra were acquired in MS1 mode with a resolution of 120,000, spanning a mass range of 350-2000 m/z, employing standard automatic gain control settings and automatic maximum injection times. For MS2 data collection, the mass spectrometer was operated in DIA mode with a resolution of 30,000. MS2 spectra were gathered across a precursor mass range of m/z 375-1200. Isolation windows of m/z 25 were used, with 0.5 m/z overlaps, a custom AGC target of 1000 units, and 30% normalized CID collision energy.

The DIA-NN 1.9 software platform^57^ was utilized to analyze proteome data using the default settings for the software. Library generation and library-free searches were conducted using the Homo sapiens FASTA database (UP000005640). Raw data was then analyzed using the Limma package and random forest imputation was applied in R. Volcano plots were generated using the ggplot package in R.

### Arrayed CRISPR/Cas9 knockout screen of TMED9-MYC interactors

To generate Cas9-expressing P cells, the Cas9 expression vector pXPR_BRD111 was used (Broad Institute Genetic Perturbation Platform). We conducted a targeted arrayed CRISPR screen covering 105 genes (3 gRNAs per gene whenever possible) with quadruplicate for each gRNA (See Table S2 for top gRNA clones, gRNAs sourced from Broad Institute Genetic Perturbation Platform). To identify the viral titer, we used lentiviral gRNA plasmids expressing a constitutive GFP and puromycin resistance to mark and select for infected cells. The viral titer of the gRNA libraries was determined by a 6-point dose response in 384-well plates by immunofluorescence of GFP+ P cells 3 days after infection. Cas9-expressing P cells were seeded 24 h prior to transduction at a density of 8000 cells/well in 384 well CellCarrier Ultra plates pre-coated with 0.25 mg/mL Synthemax II SC Substrate (Corning, 3535). 8 μL of viral supernatants were applied to cells for 24 h in the presence of 10 μg/mL polybrene (Millipore Sigma, TR-1003-G). 24 h later infected cells were washed three times to remove viral particles and transduced cells were selected in 2 μg/mL puromycin.

After 5 days incubation, cells were fixed for 20 min in 4% PFA in PBS, washed twice, then permeabilized for 10 min with 0.5% Triton X-100 in PBS and washed once more. Cells were blocked for 10 minutes at RT with Blocking solution 1, then incubated 90 min at RT with one of the following primary antibodies in Blocking solution 1: 1:500, monoclonal Fab-A-V5H anti-MUC1-fs (AbD22655.4) or 1:2000, monoclonal mouse anti-MUC1 (StemCell Technologies, Inc., 60137,214D4). The primary antibody cocktail was incubated at RT for 1.5 h, followed by four PBS wash cycles. The secondary antibody cocktail (Alexa Fluor® 488-conjugated AffiniPure F(ab’)2 Fragment Goat anti-Human IgG, Alexa Fluor® 647-conjugated Goat anti-Rabbit IgG - all Invitrogen, and Hoechst 33342 stain, each 1:1000 dilution) in Blocking solution 1 was incubated at RT for 45 min, followed by four PBS wash cycles. Finally, plates were sealed with a Plate Loc plate and stored in a Liconic incubator at 10 °C until imaging.

For data processing, the levels of MUC1-fs (see “Fluorescence image analysis”) and the cell number parameter in each plate were normalized to the negative control (Non-targeting gRNA) and to the positive control (MUC1 gRNA) defined as 0 and −100% activity, respectively. All gRNA showing > −20% reduction in cell number were masked out. As we noted that the level of MUC1-fs in the negative control was linearly correlated to cell number, we used this correlation coefficient to correct our measured MUC1-fs values. Guides targeting *MUC1* reliably decreased MUC1-fs immunofluorescence, while NTC guides had little effect.

### BRD7635 and additional drug treatments in cells

BRD7635 was added at a concentration of 0.5-1 µM for 24 h (and 5 d for Semaphorin 4A HEK cells). Control cells were exposed to the same amount of DMSO. For shorter time periods (10 min - 30 min) a concentration of 5 µM was used. For western blot analysis, stable TMED depletion cell lines were plated at 10^6^ cells/well in 6-well plates, media was changed after 4 days, and on day 8, cells were treated 24 h with the following trafficking inhibitors or DMSO vehicle in RenaLife media: 50 nM bortezomib or 2 uM MG132, 100 nM thapsigargin, 200 ng/mL brefeldin A, or 100 nM bafilomycin A1. Treated cells were lysed and processed for Western blot. For IF experiments following inhibitor/BRD treatment, P cells were plated at 14,000 cells/well in 96-well PhenoPlate microplates in 100 µL media and treated after 48 h for a total of 24 h prior to fixation.

### Immunofluorescence staining in cells

Cells grown on PhenoPlate 384-well (Revvity, 6057308) or 96-well microplates (Revvity, 6055308) were fixed 20 min in PBS containing 4% PFA (Electron Microscopy Sciences) and permeabilized for 15 min in 0.5% Triton X-100 (Sigma-Aldrich). Cells were then washed 2x with PBS and 2x with MilliQ water and treated for 30 min at RT with antigen retrieval buffer. Following treatment with antigen retrieval buffer, cells were washed four times in PBS, blocked in Blocking buffer 1 or 2 for 1 h, and washed four times in PBS again before either 90 min treatment at 37°C or 60 min at RT followed by 60 min at 37°C with primary antibodies diluted in Blocking buffer 1 containing 0.05% Tween-20 or LICOR Intercept (PBS) Blocking Buffer (see Table S5 and S7 for List of antibodies). Cells were then treated with secondary stains diluted in Blocking buffer 1 or 2 for 60 min at RT, washed four times in PBS, and stored under PBS for imaging. To reduce radical formation during imaging, cells were maintained in an Imaging Buffer during image acquisition. Finally, plates were sealed with a Plate Loc plate and stored in a Liconic incubator at 10 °C until imaging.

List of other antibodies used for IF: mouse monoclonal anti-MYC (1:1000, CST 2276S), human anti-MUC1-fs (1:1000, BioRad AbD22655.4), mouse monoclonal Fab-A-V5H anti-MUC1-fs (1:500), mouse monoclonal anti-FLAG (1:1000, Sigma Aldrich, A8592). For TMED antibodies please refer to Figure S1 and Table S7, for organelle antibodies please refer to Table S5.

List of secondary antibodies (all used at 1:1000 dilution): Donkey anti Mouse IgG (H+L) Highly Cross Adsorbed Secondary Antibody, Alexa Fluor 488 (Thermo Scientific, A-21202), Donkey anti-Mouse IgG (H+L) Highly Cross-Adsorbed Secondary Antibody, Alexa Fluor Plus 568 (Thermo Scientific, A10042), Donkey anti-Goat IgG (H+L) Highly Cross-Adsorbed Secondary Antibody, Alexa Fluor Plus 647 (Thermo Scientific, A32849), Alexa Fluor® 647-conjugated AffiniPure F(ab’)2 Fragment Goat Anti-Human IgG (Jackson ImmunoResearch, 109-606-097), Alexa Fluor® 647-conjugated Goat anti-Rabbit IgG (Invitrogen, A21246), Alexa Fluor® 568-conjugated Goat anti-Rabbit IgG (Invitrogen, A011004), Alexa Fluor® 488-conjugated anti-human IgG (Jackson ImmunoResearch, 109-546-097), Alexa Fluor® 488-conjugated AffiniPure F(ab’)2 Fragment Goat anti-Human IgG, and Hoechst 33342 stain (1:5000, Invitrogen H3570).

### BRD7635 treatment *in vivo*

MUC1-fs mice: A male cohort of 4 month-old *MUC1^fs/+^* and *Muc1^+/+^* mice were treated by oral gavage for 5-7 days with varying concentrations of BRD7635 or vehicle.

RHO-P23H mice: *Rho^P23H/+^* pups were randomly assigned to either the drug-treated experimental group or the untreated control group. To accommodate the early developmental stage of the animals, a two-phase oral administration protocol was utilized. From postnatal day 11 (P11) to P21, pups were orally administered the drug *via* a pipette. From P21 until the completion of the 21-day treatment regimen at P31, administration was transitioned to standard oral gavage (gauge 24 straight, Fine Science Tools #18061-24). Drug dosing was adjusted dynamically based on individual body weight to maintain a dose of 1 mg/kg (equivalent to 1.22 mg/kg when accounting for the salt-to-free base conversion). A male cohort of four-month-old wild-type *Rho^+/+^* and heterozygous *Rho^P23H/+^* mice was treated by oral gavage with vehicle or 1.5-month treatment with 3.0 mg/kg BRD7635, then 1.5-month treatment with 1.0 mg/kg BRD7635, for imaging. After the final dose, the mice were sacrificed, the eyes extracted, and the lenses removed from the eyes before the tissues were flash frozen in liquid nitrogen and stored at −80 °C.

### Mouse organ processing

Mice were anesthetized with 3 L/min of 3% isoflurane in O_2_ for 5 min (Combi-vet system; Rothacher Medical, Bern, Switzerland). Anesthetized mice were transcardially perfused with 0.1 M PBS (pH 7.4). For immunofluorescence studies, kidney capsules were removed. For whole kidney lysate studies, tissue was either snap-frozen whole in liquid nitrogen or lysed immediately using a tissue homogenizer (Tissue Tearor, Polytron) in Co-IP buffer. Mouse eyes were removed, enucleated and immediately frozen in Tissue-Tek OCT compound (Sakura Finetek USA, Torrance, CA) for imaging. Retina sections from the other eye were homogenized using a Kinematica POLYTRON PT1200 Hand Dispenser in 100 - 300 µL of Co-IP Lysis Buffer. A total of 50 µg of protein (kidneys) and 5 - 18 µg of protein (retina) were loaded per lane for Western blotting.

### Immunofluorescence staining in mouse kidney

5 µm kidney cryosections were placed in a 24-well Sensoplate (Greiner Bio-One, 662892) coated with (3-Aminopropyl) triethoxysilane (APTES, VWR, IC15476680). Sections were air-dried for 10 min followed by a 3 min PBS wash. Sections were then blocked in LICOR Intercept® blocking solution (Neta Scientific, 927-70001) supplemented with 0.05% TritonX-100 (Sigma, CAS 9002-93-1), followed by overnight incubation at 4 °C with the following primary antibodies diluted in blocking buffer: human anti-MUC1-fs (Bio-Rad, AbD22655.4, 1:200); rabbit anti-NCC (StressMarq Biosciences, SPC-402D, 1:500); FITC-conjugated Lotus Tetragonolobus (Asparagus Pea) Lectin (LTL) (Invitrogen, L32480, 1:1000). Following primary incubation, tissues were washed with PBS then incubated in secondary antibodies (Invitrogen or Jackson) diluted 1:500 in blocking solution for 2 h at room temperature. Lastly, sections were incubated with DAPI diluted 1:5000 in PBS for 15 min, washed three times and left in PBS for imaging.

### Immunofluorescence staining in mouse retina

For retinal IF images, 10 µm cryosections were taken through the vertical meridian, including the optic nerve head. Primary mouse anti-mutant specific rabbit anti-P23H rhodopsin antibody (Innovagen, made from a short synthetic peptide containing the mutation) was diluted at 1:3000, respectively, in blocking buffer (5% donkey serum + 2% BSA in 0.2% Triton-X in PBS) and incubated at 4 °C overnight. Secondary incubation steps were performed as stated above. After the last wash, anti-fade mounting medium was added to slides and a number 1.5 coverslip was applied. IF of mouse retinal sections from BRD4780-treated mice were taken at 5 µm at the optic nerve and placed in 24-well Sensoplates.

### Fluorescence image acquisition

All fluorescence imaging was performed using the Opera Phenix High-Content Screening System (PerkinElmer). For fluorescence imaging of cells, Phenoplates (96- or 384-well, Revvity) were used, and a minimum of nine fields was acquired per well using 20x/40x or a minimum of 16 fields and two planes (1.5 um distance between) using 63x water immersion objectives in a confocal mode. For mouse kidney section imaging, 24-well Sensoplates (Greiner Bio-One) were used. The entire specimen was first imaged for DAPI at 5X using the PreciScanTM feature (Perkin Elmer) to identify tissue. Pre-identified tissue regions were then imaged at higher resolution (20X, water immersion objectives, confocal mode). For retinal sections, images were acquired using a Zeiss Axio Imager M2 equipped with a Plan-Apochromat 20X/0.8 M27 objective. Identical settings were applied for all slides.

### Fluorescence image analysis

Image analysis for all imaging experiments was performed using Harmony software (PerkinElmer).

#### Cells

Cell nuclei were first identified using Hoechst staining, and cell number was calculated in each well. Cytoplasmic regions were then detected around each nucleus based on combined channels. Cells at the edges of the field were eliminated from the analysis. To study subcellular distribution and trafficking of MUC1-fs and TMED proteins, the “spot” identification feature was used to detect the protein of interest and different organelles.

For the quantification of protein abundance, total signal intensity in the spots for each antibody was calculated separately in the cell cytoplasm and the average signal per cell was calculated for each well unless mentioned otherwise. GFP signal in acute TMED knock-down in HEK-RHO-P23H cells was analyzed per field and then averaged per well. GRASP55 signal in P cells with GRASP55 knockdown was sorted depending on signal intensity into “low” and “high”. MUC1-fs signal was measured in cells with low and high GRASP55 intensity. For the quantification of protein “A” localization in a particular organelle “B”, three parameters were measured in each well: the total overlap area between spot A and B; the total area of the A spots and the total area of the B spots. The proportion of co-localization was counted as the area of overlap / area of a protein A / area of protein B (Figure S3B).

#### Kidney and retina

For image analysis of MUC1-fs expression in mouse kidney sections, MUC1-fs intensities were calculated in kidney sections of mice treated with either vehicle or BRD7635. As MUC1-fs levels varied in different kidney regions, and as sections could contain different portions of kidney regions, the levels of these proteins were systematically analyzed only in NCC-positive distal convoluted tubules. To this end, single cell nuclei were first identified using the DAPI channel, followed by cytoplasm detection using all combined channels, excluding the nuclei. Each fluorescent channel intensity was measured, and a threshold was set for the identification of NCC-positive cells. MUC1-fs levels were then calculated only in the NCC-positive cells. MUC1-fs protein was detected using the “spot” identification feature. For the quantification of MUC1-fs protein abundance, the total signal intensity in the spots was calculated and the average signal per cell was calculated for each well.

Tubule dilation was analyzed in LTL-positive proximal tubules in kidney sections from mice treated with vehicle or with BRD7635 (Figure S7A-G). The tissue section was identified based on DAPI, LTL and NCC staining and split into “Manhattan blocks”. Within each block, intensity and texture parameters for all channels were calculated and the “linear classification” in Harmony was used to identify Cortex, Outer and Inner medulla based on those parameters. Within the outer medulla area the “image region” below threshold intensity of the LTL channel and above the threshold for area, was selected. The shape of these regions; the intensity and texture of their borders in the LTL channel were calculated for each region and those parameters were used for “linear classification” based separation between “real tubules” (green) and gaps (red). The average width parameter for those regions was then used to calculate the tubular dilation. Before the treatment experiment, we performed a “Natural History” study, measuring the tubular dilation in *Muc1^+/+^* and *Muc1^fs/+^* mice at ages 3, 4, 5, 6, 7 and 8 months. We found that while the 3 month-old mice did not differ, starting at 5 months the *Muc1^fs/+^*mice tubules were larger than *Muc1^+/+^*mice tubules (Figure S7G-H). To prevent tubular dilation in *Muc1^fs/+^* mice, we decided to treat three-month old mice for three months.

For retinal sections, image analysis was conducted utilizing ImageJ software (NIH, Bethesda) paired with a custom macro designed for semi-automated quantification of signal intensity, minimizing potential biases. In brief, merged channel images were opened and the region between the outer nuclear layer and the retinal pigment epithelium was delineated. Subsequently, images of specific channels were examined. Within these, the area of interest was selected, and the integrated density for each channel was quantified.

### Masson’s Trichrome Stain, PAS stain for kidney sections and Hematoxylin and Eosin (H&E) stain for retina sections

For kidneys, slides with 5 µm sections (sagittal cut, from 4% PFA tissue in OCT) were sent to iHisto (Salem, MA) for further processing. Eyes were flash frozen, embedded in OCT and blocks were sent to iHisto. Sections were obtained by cutting at the level of the insertion of the optic nerve. iHisto uses a semi-automated staining workflow for consistency, including a Leika ST5020 Autostainer, TS5025 Transfer Station and CV5030 Coverslipper.

### Nano differential scanning fluorimetry (nanoDSF)

TMED7, 2, 9 and 10 was added to HEK293 cells at 1:1:1:1 stoichiometry, followed by purification of the TMED7/2/9/10 complex using gentle detergents (LMNG, 5% and HCS, 0.5%). nanoDSF experiments were performed on a Prometheus NTPlex instrument (NanoTemper Technologies). Briefly, a temperature ramp of 20 °C to 90 °C, 1 °C per minute, was applied to the purified TMED7/2/9/10 complex, at 0.5 mg/mL apo protein, in solution with 10 μM, 50 μM, or 100 μM BRD7635, BRD4780, or DMSO. The melting temperature (Tm) of the TMED7/2/9/10 complex in each condition, or the temperature at which 50% of the TMED7/2/9/10 complex was unfolded, was determined from the first derivative curve midpoints of the 350/330 nm fluorescence emission ratios plotted against the temperature range.

### Golgi-Immunoprecipitation (Golgi-IP)

P cells were infected with TMEM115-3xHA virus (pLJC5 KOZAK TMEM115 3HA, University of Dundee, DU68534)^58^ or the backbone (pLJC5 short Kozak 3HA, University of Dundee, DU70022) alone, followed by selection with puromycin to create a stable cell line. Expression of TMEM115-3xHA was validated using immunofluorescence and western blot. Golgi-IP P cells and control cells were grown to confluence in a 15cm dish followed by Golgi-IP as described previously^58^. In short, cells were quickly washed with ice-cold PBS, then scraped into 1mL of ice-cold KPBS (136 mM KCl, 10 mM KH_2_PO_4_. Adjust to pH 7.25 with KOH). After a quick centrifugation, cells were resuspended in 1 mL of KPBS and a sample was taken for “Cell lysate”. Cells were gently homogenized on ice using a 2 mL Dounce Homogenizer, a sample was taken for “IP-input”. Cell homogenate was quickly centrifuged to remove cell debris and nuclei, then added to 150 μl washed HA-beads (Thermo Scientific 88837) for 15 minutes rotating in the cold. Samples were put on a DynaMag-2 magnetic rack and a sample from the supernatant was taken for the “Unbound” fraction. Beads were gently washed 3 times with 1mL of ice-cold KPBS. After the last wash, beads were eluted using 60 μl of ice-cold lysis buffer (50 mM HEPES pH 7.4, NaCl 40mM, EDTA 2mM, 1% Triton X-100, 10 mM β-glycerol phosphate, 10 mM Na-pyrophosphate, 1.5 mM NaV0_4_, 50 mM NaF, supplemented with Complete EDTA-free Protease Inhibitor Cocktail (Roche 04693159001)) for 10 minutes. For crosslinking, DSSO at final concentration of 5 mM in DMSO was added and incubated with Golgi lysate for 30 min at 25°C prior to quenching with 50 mM Tris-HCl, pH 8.0. Beads were then separated from the eluate using a DynaMag-2 magnetic rack. The eluate contains the Golgi fraction and was prepared for western blot as indicated below. After centrifugation of “Cell lysate” samples 100 μl of lysis buffer was added for 20 minutes on ice. “IP-input” and “Unbound” samples were also incubated with 100 μl of lysis buffer, then all samples were centrifuged and prepared for western blot analysis or Coomassie staining (Laemmli SDS-Sample Buffer, BP-110NR; NuPage reducing agent, NP0009; boiled at 75°C for 10 minutes). Western blot was run as detailed above, using a 3-8% Tris-Acetate gel (Thermo Fisher EA0378) and Tris-Acetate running buffer (Thermo Fisher LA0041) and HiMark pre-stained protein standard (Thermo Fisher LC5699). For MS, samples were run on a 3-8% Tris-Acetate gel and stained with PageBlue Protein Staining Solution (Thermo Scientific 24620), destained with MilliQ water and respective bands were cut out and stored in −80°C for further processing.

### Mass spectrometry analysis of Golgi-IP gel bands

Excised gel bands were cut into approximately 1 mm^3^ pieces. Gel pieces were then subjected to a modified in-gel trypsin digestion procedure^59^. Gel pieces were washed and dehydrated with acetonitrile for 10 min. followed by removal of acetonitrile. Pieces were then completely dried in a speed-vac. Rehydration of the gel pieces was with 50 mM ammonium bicarbonate solution containing 12.5 ng/µl modified sequencing-grade trypsin (Promega, Madison, WI) at 4°C. After 45 min., the excess trypsin solution was removed and replaced with 50 mM ammonium bicarbonate solution to just cover the gel pieces. Samples were then placed in a 37°C room overnight. Peptides were later extracted by removing the ammonium bicarbonate solution, followed by one wash with a solution containing 50% acetonitrile and 1% formic acid. The extracts were then dried in a speed-vac (∼1 hr). The samples were then stored at 4°C until analysis. On the day of analysis the samples were reconstituted in 10 µl of HPLC solvent A (2.5% acetonitrile, 0.1% formic acid). A nano-scale reverse-phase HPLC capillary column was created by packing 2.6 µm C18 spherical silica beads into a fused silica capillary (100 µm inner diameter x ∼30 cm length) with a flame-drawn tip^60^. After equilibrating the column each sample was loaded via a Thermo EASY-LC (Thermo Fisher Scientific, Waltham, MA). A gradient was formed and peptides were eluted with increasing concentrations of solvent B (90% acetonitrile, 0.1% formic acid).

As peptides eluted they were subjected to electrospray ionization and then entered into a Orbitrap Exploris480 mass spectrometer (Thermo Fisher Scientific, Waltham, MA). Peptides were detected, isolated, and fragmented to produce a tandem mass spectrum of specific fragment ions for each peptide. Peptide sequences (and hence protein identity) were determined by matching protein databases with the acquired fragmentation pattern by the software program, Sequest^61^ (Thermo Fisher Scientific, Waltham, MA). All databases include a reversed version of all the sequences and the data was filtered to between a one and two percent peptide false discovery rate.

### Purification of TMED entrapment complex

pTT5-hTMED9(1-37)-3*Flag-hTMED9(38-235)-P2A-hTMED2(1-20)-10*His-hTMED2(21-201) and pTT5-hTMED10(1-31)-HAtag-hTMED10(32-219)-P2A-hTMED7(1-34)-V5tag-hTMED7(35-224) were overexpressed in Expi293F GnTI cells. In short, cells were split to a density of 2.7-3.0 ×10^6^ cells/mL and 0.5mg DNA per 0.5L of culture together with 3mg PEI transfection reagent were added. Cells were incubated on an orbital shaking platform at 37 °C with 8% CO_2_ at a speed of 125 rpm for 20 h prior to treatment with Valproic acid sodium salt (Sigma, P4543)(200mM stock in water, final concentration in the culture: 1mM). Cell pellet was harvested 48 h post transfection. The final total cell density of the culture was 2.4×10^6^ cells/mL, and cell viability was 60%. The cells were centrifuged at 2000 rpm, 4 °C for 5 min. The cell pellet was collected, resuspened with PBS, and then centrifuged at 2000 rpm, 4 °C for 5 min again.

The cells were resuspended in lysis buffer (5 mL/1 g, 20 mM Tris at pH 8.0, 150 mM NaCl) with Protease inhibitor cocktail (Roche) at 50 mL for one piece. Then it was added with 1% (w/v) DDM-0.2% (w/v) CHS (final concentration), and then incubated at 4 °C for 2 hrs. Then it was centrifuged at 13000rpm (20300*g*), 4 °C for 30 min, and the supernatant was collected and centrifuged again. We first ran a Flag column, with a column volume (CV) of 35mL(ANTI-FLAG® M2 Affinity Gel, Sigma, A2220-25ML). The resin was pre-equilibrated with buffer A (20 mM Tris at pH 8.0, 150 mM NaCl, and 0.05% (w/v) DDM-0.01% (w/v) CHS), and then incubated with the supernatant for 2 h at 4 °C on a rotator. The resin was washed in the column with buffer A, B (5 CV, 20 mM Tris at pH 8.0, 150 mM NaCl, and 0.05% (w/v) DDM-0.01% (w/v) CHS, 5 mM ATP, 10 mM MgCl_2_;), A until no signal was observed by G-250. The target protein was eluted from the column with buffer C (20 mM Tris at pH 8.0, 150 mM NaCl, and 0.05% (w/v) DDM-0.01% (w/v) CHS, 250 ug/mL Flag peptide) and D (20 mM Tris at pH 8.0, 150 mM NaCl, and 0.05% (w/v) DDM-0.01% (w/v) CHS, 500 ug/mL Flag peptide) and a 12% SDS-PAGE was run. We then ran a Ni-NTA column (QIAGEN, No. 30210) with a CV of 3 mL. The resin was pre-equilibrated with buffer A, and then incubated with the target protein for 2 hs at 4 °C on a rotator. The resin was washed in the column with buffer A, E (20 mM Tris at pH 8.0, 150 mM NaCl, and 0.05% (w/v) DDM-0.01% (w/v) CHS, 10 mM imidazole), F (20 mM Tris at pH 8.0, 150 mM NaCl, and 0.05% (w/v) DDM-0.01% (w/v) CHS, 20 mM imidazole) and G (20 mM Tris at pH 8.0, 150 mM NaCl, and 0.05% (w/v) DDM-0.01% (w/v) CHS, 250 mM imidazole) until no signal was observed by G-250 and a 12% SDS-PAGE was run. The target protein was carefully concentrated by ultra-filtration tube (Amicon Ultra, Merck Millipore) at the molecular cutoff 10kD, and then loaded onto a Superose^TM^ 6 Increase 10/300 GL column (CV = 24 mL, GE, pre-packed, Cytiva, 29091596). The column was pre-equilibrated with final buffer (50 mM Hepes at 7.5, 150 mM NaCl, 0.02% (w/v) DDM-0.004% (w/v) CHS), and then the target protein was loaded onto the column. The column was washed with final buffer and a 12% SDS-PAGE was run. The target protein was carefully concentrated by ultra-filtration tube (Amicon Ultra, Merck Millipore) with the molecular cut off of 10 kDa. It was aliquot, flash frozen with liquid nitrogen and then stored at −80 °C. A standard curve was developed by running “Gel Filtration Standard” (BIO-RAD, 1511901, dissolved in 1 mL water, 50ul for injection). Fraction 16 from 250 mM imidazole wash was visualized on an SDS page gel.

### Phylogenetic tree

Human TMED family protein sequences were retrieved from UniProt (reviewed, canonical isoforms; organism ID 9606) and aligned using MUSCLE in the msa R package, excluding TMED8. The resulting alignment was converted to a phyDat object in phangorn to compute a maximum-likelihood distance matrix, from which a neighbor-joining tree was constructed using ape as the starting topology for phylogenetic inference^62^. Maximum-likelihood optimization was performed under the WAG amino acid substitution model with stochastic tree rearrangements, and branch support was assessed with 1,000 bootstrap replicates using nearest-neighbor interchange. The final tree was ladderized, annotated with bootstrap values, and visualized using ggtree.

### AlphaFold3 Prediction of TMED and GRASP55 Binding Interface

Initial structural models of the GRASP55-TMED5, GRASP55-TMED7 were generated using the AlphaFold3 multimer prediction server^63^. Signal peptides of TMED proteins (as per UniProt annotation) were excluded during complex prediction to focus on mature protein interactions.

### All-Atom Molecular Dynamics Simulations

All-atom Molecular Dynamics (MD) simulations were performed to refine the conformational ensembles and evaluate the stability of the predicted complex using GROMACS 2022.5^64^. The initial structure models were solvated in explicit TIP3P water^65^ with 0.15 M NaCl, yielding systems of 108,768 and 112,134 atoms. The CHARMM36m force field^66^ was applied to all molecular components. To maintain membrane anchoring while allowing conformational flexibility elsewhere, positional restraints were applied to transmembrane helix backbones; the cytosolic tail and GRASP domains remained unrestrained. Initial minimization and equilibration followed standard CHARMM-GUI protocols^67^.

All production MD simulations were carried out in the NPT ensemble at 310 K and 1 bar using with CHARMM36m force-field^66^. The leap-frog integrator was used with a 2 fs time step. Temperature was controlled via the velocity-rescale thermostat^68^ (1.0 ps coupling constant), and pressure was maintained isotropically using the Parrinello–Rahman barostat^69^ (2.0 ps coupling, compressibility 4.5 × 10⁻⁵ bar⁻¹). Bonds involving hydrogen atoms were constrained using the LINCS algorithm^70^. Long-range electrostatics were computed via the Particle Mesh Ewald method (1.2 nm real-space cutoff), and van der Waals interactions employed a force-switch function (1.0–1.2 nm). Center-of-mass motion was removed every 100 steps. Positional restraints were retained on transmembrane helix backbones to preserve membrane orientation. Each system was simulated for 200 ns in 5 independent replicates, reaching an aggregate sampling time of 1 μs. Analyses were performed using the final 100 ns of each trajectory. Hydrogen bonding and salt bridge interactions were quantified using VMD (Visual Molecular Dynamics) 1.9.4 plugins^71^. Hydrogen bonds are defined when a distance between a hydrogen donor (D) and acceptor (A) atom is less than 3.2Å and the D-H-A angle is less than 60 degree. Salt bridge formation was defined as distance between the oxygen atoms of acidic residues and the nitrogen atoms of basic residues within 3.2Å. To find the representative conformational ensemble, TTClust^72^ software was used to perform hierarchical clustering and the most representative frames of the largest cluster were selected.

### Electroretinogram (ERG) and Optical Coherence Tomography (OCT) in mice

C57BL/6J mice heterozygous for the P23H rhodopsin mutation and homozygous for the TMED9 floxed allele, with or without a tamoxifen-inducible CAGGCre-ER transgene were used to assess the effect of TMED9 knockout on retinal structure and function. The knockout group consisted of 14 Cre-positive (Cre+) mice (10 male, 4 female), and the control group comprised 11 Cre-negative (Cre−) littermates (5 male, 6 female). All mice received intraperitoneal tamoxifen injections on postnatal days 21, 23, and 25 to induce recombination selectively in Cre+ mice.

Electroretinography (ERG) was used to evaluate retinal function using a Celeris system (Diagnosys, Lowell, MA), and spectral-domain optical coherence tomography (SD-OCT) using the Envisu system (Bioptigen, Durham, NC) was used to evaluate retinal morphology. Both modalities were performed in succession during a single imaging session at P42. Eyes were harvested after the procedures for Western blot analysis. Before recording, a mouse was dark adapted for at least 4 hr. Under dark conditions and dim long-wavelength illumination using headlamps, mice were anesthetized via intraperitoneal injection of a ketamine–xylazine mixture, prepared from stock solutions of ketamine 100 mg/mL (Dechra, KET-10) and xylazine 100 mg/mL (Pivetal, 69043-0043-05) to yield final concentrations of 10% ketamine and 1% xylazine. The anesthetic solution was administered at a dose of 10 µL per gram of body weight. Following induction, mydriasis was achieved using topical administration of tropicamide 1% and phenylephrine HCl 2.5% solution (Pine, 762). Corneal anesthesia was achieved by administration of proparacaine HCl ophthalmic solution, USP 0.5% (Sandoz, AK2D15DS). The corneas were hydrated with the application of hypromellose 0.3% eye gel (Alcon, AX11933). A mouse was placed on the Celeris platform maintained at 37 °C and electrodes were positioned on its corneas ^41^. A series of flashes (0.01, 0.1, 1 cd·s/m^2^) were given in sequence; each eye’s response was the average of three sweeps. Software interstimulus intervals were 5, 10, and 10 seconds, which became 10, 20, and 20 seconds per eye as flashes were presented monocularly and alternated between left and right. The unstimulated eye was used as the reference input. Responses were recorded from both eyes. B-wave amplitudes (from the 0.1 cd·s/m^2^ flash) were quantified from the scotopic ERG data using Diagnosys Espion software. Following ERG recording, SD-OCT imaging was performed in mice as previously described^40^. Each mouse was positioned in a custom imaging cassette that allowed multi-axis rotation for optimal eye alignment, and the cornea was kept hydrated with hypromellose 0.3% eye gel. After imaging, all retinal layers from the inner limiting membrane to the retinal pigment epithelium were segmented by a blinded retina specialist. Thickness metrics for each retinal sublayer and the total retina were then extracted using the optic nerve head (ONH) as a fiduciary reference point. Retinal thickness was sampled at predefined eccentricities from the ONH and these values were averaged to yield representative thickness measurements for each eye. Layer-specific thickness values such as outer nuclear layer (ONL) thickness were used for subsequent quantitative analysis.

Data preprocessing and statistical modeling were performed using Python, primarily utilizing the pandas and statsmodels libraries. To ensure data robustness, outliers in b-wave electroretinogram (ERG) measurements and outer nuclear layer (ONL) thickness assessed by optical coherence tomography (OCT) were identified and excluded using the standard 1.5× interquartile range (IQR) rule. This outlier detection was applied within subgroups defined by reading session, eye, genotype, and sex. For ERG and OCT measurements at P42, generalized estimating equations (GEE) were employed to model each outcome variable (ONL thickness and ERG amplitude) as a function of genotype (Cre+ versus Cre−), adjusting for sex differences due to imbalance between groups. GEE was chosen because it accounts for the correlated nature of repeated measurements within subjects, in this case measurements from both eyes of the same mouse.

### Electroretinogram (ERG) in P23H mice treated with BRD7635

Mice were dark-adapted for 24 hr prior to recordings. Under a dim safety red light, mice had their pupils dilated with the topical administration of 1% tropicamide ophthalmic solution (Akorn, 17478-102-12) and 10% phenylephrine ophthalmic solution (MWI Animal Health, 054243). Corneal hydration was maintained with the application of Eye Lube Plus Lubricating Gel (OptixCare). Mice were anesthetized by isoflurane inhalation and placed on a heated Diagnosys Celeris rodent-ERG device (Diagnosys LLC). Ocular stimulator electrodes were placed on the corneas, the reference electrode was positioned subdermally between the ears, and a ground electrode was placed in the hind leg. The eyes were stimulated with a green-light flash stimulus (peak emission 544 nm, bandwidth approximately 160 nm) of −0.5 candela-second per meter squared (cd·s/m^2^) light intensity. The responses to 10 stimuli with an inter-stimulus interval of 10 s were averaged, and the a- and b-wave amplitudes were acquired from the averaged ERG waveform. Data were analyzed with the Espion V6 software (Diagnosys LLC). To ensure data robustness, outliers in b-wave electroretinogram (ERG) measurements were identified and excluded using the standard 1.5× interquartile range (IQR) rule. This outlier detection was applied within subgroups defined by reading session, eye, genotype, and sex. For ERG measurements at P32, generalized estimating equations (GEE) were employed as described above. b-wave amplitude was compared between the treatment group (BRD7635) and no treatment group.

### Statistical analysis

Statistical analysis was performed and presented using Graphpad Prism version 10 software. All data are presented as means ± standard deviation unless otherwise specified in the figure legends. Values of ‘N’ for each experiment can be found in the figure legends and are biological replicates unless otherwise indicated. Statistical comparisons of two groups for a single variable with normal distributions were analyzed by unpaired t-test or Mann-Whitney test if not normally distributed. Statistical comparisons of two or more groups with one independent variable were analyzed by unpaired t-test if normally distributed, or by One-way ANOVA with Tukey post-tests or Dunnett’s post-tests. Statistical comparisons of two or more groups with two independent variables were analyzed by Two-way ANOVA with Tukey and Dunnett’s post-tests. Specific statistical tests are described in their respective Methods section. *p < 0.05 **p < 0.01 ***p < 0.001 ****p < 0.0001.

### Data Availability

The original mass spectra and the protein sequence database used for searches have been deposited in the public proteomics repository MassIVE (http://massive.ucsd.edu) and are accessible at ftp://MSV000093853@massive.ucsd.edu when providing the dataset password: TMED9. If requested, also provide the username: MSV000093853. These datasets will be made public upon acceptance of the manuscript.

## Acknowledgement

We would like to thank Fabian Schulte, Simon Butterworth and Brooke Linnehan from the Quantitative Proteomics Core at the Whitehead Institute, Cambridge, MA, and Steven Gygi and Julian Mintersis, Harvard Medical School, Cambridge, MA, for their help with MS-IP experiments. Shewa Osmani from the Vavvas lab helped with analysis of the ERG wave-forms. Schematics created in BioRender. Racette, M. (2026) https://BioRender.com/8kilu25.

## Author Contributions

MRK, ACG, MKA, JLP, and AG conceived the study and designed the experiments. MKA, PB, MB, BS and RM conducted the immunofluorescence experiments assessing endogenous localization of TMEDs, arrayed CRISPR KO screen, rescue of TMED depletion cells, TMED degradation after BRD7635 treatment and localization of rhodopsin mutants. MKA, PB, and EM conducted dose-response experiments. KK, JL, and MKA performed staining of MUC1-fs mice, Western blotting, and tissue analysis of mutant MUC1 mice. ERC, KK, EG, TN, GS, VC, NE, DN, TC, SO, ZD, KTN, CRM, KP and DV performed tissue analysis, ERG and OCT of mutant rhodopsin mice. SBV, MDL, DC, NU, and SC assisted with IP-MS experiments. EG, FMA, and HY assisted with immunoprecipitation experiments. JZ and JCI helped with data analysis. MR and ERC assisted with management of the mouse colonies. YM, SK and SI performed simulation of TMEDs with GRASP55. All other experiments detailed in this manuscript were conceived and conducted by MRK, ACG, and CDDM in consultation with the co-authors. MRK, ACG, JLP, and AG wrote the manuscript and all co-authors contributed to review and revision of the manuscript.

## Declaration of interests

A.G. serves as an advisor, consultant or board member to several biotechnology companies based on agreements reviewed and managed by Mass General Brigham and the Broad Institute of MIT and Harvard in accordance with their conflict-of-interest policies. S.A.C. is a member of the scientific advisory boards of Kymera, PTM BioLabs, Seer and PrognomIQ.

## Funding

This work was made possible by support from US National Institutes of Health (NIH) grants DK095045 (A.G.), DK099465 (A.G.), T32GM007753 (A.C.G), T32GM144273 (A.C.G), T32GM007226 (A.C.G.), F30DK127546 (A.C.G.), the Cure Alzheimer’s Fund, and the Slim Initiative in Genomic Medicine for the Americas (SIGMA), a collaboration of the Broad Institute with the Carlos Slim Foundation. This work was supported in part by grants from the National Cancer Institute (NCI) Clinical Proteomic Tumor Analysis Consortium grants NIH/NCI U24-CA210986 and NIH/NCI U01 CA214125 (to SAC). MRK was supported by a KRESCENT post-doctoral fellowship from The Kidney Foundation of Canada and a Ben J. Lipps post-doctoral fellowship from the American Society of Nephrology.

## Supplementary Figures/Tables

**Figure S1. Intracellularly accumulated mutant rhodopsin interacts and co-localizes with TMED9.**

(**A**) Phylogenetic tree depicting all TMED proteins and their relationships. Scale bar depicts the number of amino acid substitutions per residue. Numbers at tree branches depict the bootstrap support values in %. (**B**) (Left) Schematic of rhodopsin (RHO) with RP-causing missense (green) or nonsense (black) mutants indicated. (Right) Co-IP of TMED9 upon GFP IP from dox-inducible HEK293T GFP-tagged RHO-WT and RHO mutant-expressing cell lines. Green font, rhodopsin mutants exhibiting intracellular accumulation. Input on the right. N = 3 biological replicates. (**C**) IF images of RHO-GFP (green) in dox-inducible HEK293T RHO-WT-GFP and RHO-mutant-GFP expressing cell lines. Scale = 20 µm. (**D**) Representative IF images of RHO-GFP (green) and endogenous TMED9 (red) in dox-inducible HEK293T RHO-P23H and RHO-WT expressing cell lines. N = 4 biological replicates. Scale = 20 μm. (**E**) Quantification of (D). RHO-GFP colocalization with endogenous TMED9 by normalized area of overlap. Means ± SD. N = 4 biological replicates. 7,600 - 8,800 cells analyzed per replicate. ** = p < 0.01.

**Figure S2. Depletion of entrapment TMEDs leads to clearance of mutant protein cargo while depletion of non-entrapment TMEDs leads to cargo accumulation.**

(**A**) Co-IP of endogenous MUC1-fs and TMED9 from P cells. N cells are a negative control for MUC1-fs antibody signal since they do not express MUC1-fs. IgG, unimmunized rabbit control IgG Co-IP. (**B**) Co-IP of endogenous TMED9 with other TMEDs from P cells. IgG, unimmunized rabbit control IgG Co-IP. Lysate, lysate input. (**C-D**) Western blots of endogenous TMEDs (**C**) and endogenous MUC1-fs (**D**) in CRISPR/Cas9 stable TMED depletion P cells. TMED depletion lines (gTMED**X**-1 and -2) were generated from two separate gRNAs. Antibodies used are therefore knock-out validated, see Table S7. Stable depletion lines used in Figure 1J are indicated (red, blue, or bolded black). Blue, entrapment TMEDs; Red, non-entrapment TMEDs. Primary MUC1-fs band denoted by arrowhead. NTC, non-targeting control. (**E**) Representative Western blots of HEK RHO-P23H-GFP cells after acute knock-down of TMED5, TMED7, or TMED9. Anti-GFP antibody was used to detect RHO-P23H. GAPDH was used as loading control. (**F**) Quantification of (E). N = 3 biological replicates. Means ± SD. * = p < 0.05, ** = p < 0.01. (**G)** IF images of RHO-P23H HEK293T cells with TMED5, TMED7, or TMED9 knockout. Scale = 50 μm. (**H**) Analysis from (G) of GFP/Hoechst intensity normalized to gNTC. Means ± SD. ** = p < 0.01, *** = p < 0.001. N = 3 biological replicates. 60,000 - 80,000 cells analyzed per genotype per replicate.

**Figure S3. Entrapment TMEDs (2, 7, 9 and 10) prevent anterograde trafficking of mutant cargo whereas non-entrapment TMEDs 1 and 5 do not.**

(**A**) Schematic of the secretory pathway and markers used to label organelles within the secretory pathway. ER, endoplasmic reticulum; CNX, Calnexin; E. endo., early endosome; L.endo., late endosome. Created in BioRender. (**B**) Human P cell images in three channels (SEC31A left, TMED9 middle, Merge right). Nuclei are identified based on Hoechst staining, cells based on all other channels and cytoplasm is defined as area of cell - nucleus. “Spots” are identified based on protein signal. Protein abundance and localization in a cell was calculated (see Methods). (**C**) Representative IF images of endogenous MUC1-fs (green) and select secretory pathway markers (red) in CRISPR/Cas9 stable non-targeting control (gNTC) and TMED7 depletion (gTMED7) P cells are shown. Early secretory structures, e.g. COPII vesicles are marked by SEC31A, late secretory pathway structures, e.g. early endosome vesicles were marked by EEA1. Scale = 20 μm. (**D**) Quantification of (C). Changes in MUC1-fs intensity per organelle area in P cells with depleted gTMED**X**. N = 3 biological replicates, 1 representative replicate shown. 6,000 - 21,000 cells analyzed per secretory marker per replicate. (**E**) Analysis of (C). Proportion of MUC1-fs overlap area with organelle marker normalized to gNTC (equals 0) shown for SEC31A, COPB2, EEA1 and LAMP1. (**F**) Western blot of MUC1-fs in P cells with stable depletion of TMEDs (gTMED**X**) following 24 h treatment with DMSO, 50 nM bortezomib (BOR), 100 nM thapsigargin (THAP), 200 ng/mL brefeldin A (Bref), or 100 nM bafilomycin A1 (BAF). Increasing apparent molecular weight of MUC1-fs bands represents its progressive glycosylation as it flows in an anterograde direction toward the lysosome, as previously described in Dvela-Levitt *et al*^17^.

**Figure S4. Small molecule BRD7635 binds to the entrapment complex and promotes its lysosomal degradation.**

(**A**) Western blots of loading control vinculin for Figure 2F. (**B**) Quantification of Figure 2F. Vinculin-normalized endogenous TMED signal in BRD7635-treated P cells as a percentage of TMED signal in DMSO-treated P cells. Blue, entrapment TMEDs; Red, non-entrapment TMED. Means ± SD. N = 3 biological replicates. (**C**) NanoDSF of purified TMED7/2/9/10 complex in solution with 10 μM, 50 μM, or 100 μM BRD7635 (left) vs. BRD4780 (right). First derivative (f′) of mean 350/330 nM fluorescence ratio is shown across a 1°C/min temperature ramp of 20°C to 90°C. Legend indicates compound dose and ΔTm, the change in TMED7/2/9/10 complex melting temperature between compound and DMSO conditions. N = 3 biological replicates. (**D-E**) IF images and quantification of TMED9 (**D**) and TMED7 (**E**) in P cells treated for 24 h with BRD7635 or DMSO plus either DMSO, brefeldin, bafilomycin or MG132. TMED9 (**D**) and TMED7 (**E**) spot intensities displayed as BRD signal/DMSO signal. N = 3 biological replicates. 9,000-18,000 cells analyzed per treatment condition per replicate. Only Golgi (Bref) or lysosomal (BAF) inhibition–but not proteasomal (MG132) inhibition–can reverse the BRD7635-mediated removal of TMEDs. Scale = 20 μm.

**Figure S5. TMEDs co-localize with markers of the early secretory pathway and Co-IPs with mutant cargo.**

(**A**) IF images of endogenous TMED7 (cyan) in stable gTMED5 and gTMED9 P cells. Scale = 50 μm. (**B**) Quantification of (A). TMED7/Hoechst intensity per well is shown. Blue, entrapment TMED9; Red, non-entrapment TMED5. Means ± SD. N = 4 technical replicates. 20,230 - 40,880 cells per genotype analyzed. *** = p < 0.001. (**C**) Representative IF images of endogenous TMED5 and TMED7 (green) in P cells overlaid with COPII (SEC31A) and *cis*-Golgi (GM130) markers (red). Scale = 10 μm. (**D**) Quantification of endogenous TMED localization in (C) by normalized area of overlap with secretory pathway markers (proportion of total) in P cells. Blue, entrapment TMEDs; red, non-entrapment TMEDs. Means ± SD. N = 3 technical replicates. 6,000 - 21,000 cells analyzed per secretory marker. (**E)** Co-IP of FLAG-MUC1-fs from HEK293T cells co-transfected with TMED1, TMED5 (MYC-tagged), or TMED2, TMED7, TMED9, or TMED10 (V5-tagged). Western blots of FLAG-MUC1-fs and MYC- or V5-tagged TMEDs are shown. C, corresponding negative control vector. Lysate inputs at the bottom. (**F**) (Top) Schematic of TMED9-MYC domain structure. (Bottom) HEK293T cell co-IP of FLAG-MUC1-fs with TMED9-MYC or TMED9-MYC domain deletion constructs. Deletion of the GOLD domain abrogated the interaction. SP, signal peptide; CC, coiled-coiled domain; TM, transmembrane domain; FL, full-length TMED9; ΔGOLD, TMED9 GOLD domain deletion; ΔCC, TMED9 coiled-coil domain deletion; ΔC-tail, TMED9 C-terminal cytosolic tail domain deletion; L, Lysate input; S, unbound IP supernatant.

**Figure S6. The integral Golgi protein GRASP55 binds the TMED entrapment complex.**

(**A**) Schematic of TMED5 and TMED7 domains. (**B**) Alignment of TMED5 and TMED7 protein sequences using Snapgene. (**C**) Endogenous COPB2 Co-IP probed for TMED5. Rabbit IgG, negative control. (**D**) Endogenous GRASP55 Co-IP probed for TMED2 (left), TMED9 (middle) and TMED10 (right). Rabbit IgG, negative control. Vinculin as loading control. (**E**) IF images of MUC1-WT (green) in GRASP55 negative (-) cells (2,200 - 2,600 cells per replicate) vs. GRASP55 positive (+) cells (red, 710 - 960 cells per replicate). Size bar = 20 μm. (**F**) Quantification of (E). No significant difference in MUC1-WT intensity between GRASP- cells and GRASP+ cells was observed. N = 3 biological replicates. (**G**) (Top) Schematic of Golgi-IP to isolate intact Golgi organelles. (Bottom) Control for the quality of Golgi-IP lysate by Western blot, probed for Calreticulin (ER marker) and GM130 (Golgi marker). A cell line expressing the empty vector backbone (EV) was used as negative control. The input sample was obtained after homogenization prior to the IP. (**H**) Western blot of entrapment TMEDs in Golgi-IP compared to whole-cell lysate and GOLGI-IP Input (lysate after homogenization). TMED7 and TMED2 were probed on the same blot using fluorescent antibodies (anti-mouse for TMED2 and anti-rabbit for TMED7). N = 2 biological replicates. (**I**) Alphafold3 prediction for TMED**7**-GRASP55 (left) and TMED**5**-GRASP55 (right) interactions. Red arrows indicate binding interfaces detailed in Figure 5B. The colors indicate the confidence scores for these predictions, blue indicates high confidence (and orange indicates low). (**J**) IP-MS experiment showing interactors that preferentially bind to the TMED7 CC-domain. We compared TMED7 full-length (blue) versus a TMED7 chimera with a TMED5 CC-domain (red). N= 4 technical replicates. (**K**) GO term analysis using Enrichr of proteins preferentially binding the TMED7 with its own TMED7 CC-domain. X-axis shows GO Cellular Component 2026 analysis adj. p-values. (**L**) Quantification of Figure 5E. Log_2_ fold change of MYC compared to empty vector. Mean ± SEM. N = 3 biological replicates. 5,000 - 10,000 cells analyzed per condition per replicate. * = p < 0.05. (**M**) MYC Co-IP experiment of TMED7 and TMED7-TMED5 CC chimera probing for GRASP55 and TMEDs 2, 9, 10. N = 3 biological replicates. Right, lysate inputs. Vinculin as loading control.

**Figure S7. Natural history of disease progression and dose-responsive clearance of mutant cargo by BRD7635.**

(**A**) Machine Learning image analysis methods for kidney tissue. Tissue section images were acquired in three channels. (**B**) Different areas within the tissue section were segmented using a “Linear Classification” approach to select three populations of “Manhattan Blocks” based on intensity and texture parameters of the three channels. (**C**) One 20x field of view showing the outer medulla area (green). (**D**) Within the outer medulla area, potential tubules were identified using the “find image region” feature (rainbow colors show different potential tubules). (**E**) Real tubules were identified using a “Linear Classification” approach based on morphology, intensity and texture parameters of the periphery of each “potential tubule”. (**F**) Outer medulla (rainbow colors for “outer medulla Manhattan Blocks”) for four mouse tissue sections. Note tubular enlargement in 5-month (but not in 3-month) old MUC1^fs/+^ mice. (**G**) Box-and-whisker plots of mean tubular width (µm) by age (months) in MUC1^+/+^ (blue) and MUC1^fs/+^ (red) mice. Note enlargement of tubules in MUC1^fs/+^ kidneys compared to wild-type kidneys starting at 5 months of age. (**H**) PK measurements in mice: BRD7635 plasma concentrations (ng/mL) in three female (top, M1-3)) and three male C57BL/6J mice (bottom, M1-3) following administration by oral gavage of a single dose of BRD7635 (10.0 mg/kg in saline). (**I**) IF images of MUC1-fs (green) and NCC (red) in mouse kidney cortical cross sections from representative 4 month-old male *Muc1^fs/+^* and *Muc1^+/+^*mice following 7-day BRD7635 or vehicle treatment at indicated doses. N = 3-5 mice per group. Scale = 20 μm. (**J**) Quantification of (I). MUC1-fs intensity in NCC-positive cells as a percentage of Vehicle-treated *Muc1^fs/+^* mice. Each dot is an individual mouse, with 3 technical replicates per mouse. Means ± SD. **** = p < 0.0001, Vehicle treated vs BRD7635.

## Supplementary Tables

**Table S1. Significant interactors of MYC-TMED9 in HEK293T cells co-transfected with FLAG-MUC1-fs by Log_2_FC.** Log_2_FC, Log_2_ fold change in protein abundance between MYC-TMED9 and empty vector condition. Significant interactors chosen by adjusted p-value > 0.05.

**Table S2. Arrayed CRISPR/Cas9 knockout screen of MYC-TMED9 interactors in P cells by mean normalized MUC1-fs intensity.** Mean normalized MUC1-fs intensity per target interactor gRNA is shown. MUC1-fs intensity normalized to mean MUC1 and non-targeting control (NTC) gRNA. Adjusted p-value maximum 4.0.

**Table S3. In-vitro pharmacology study for BRD7635 at 10 mM.** % Inhibition >50% of control was considered significant.

**Table S4. IP-MS of TMED5-MYC and TMED7-MYC.** Proteins (with gene names) that preferably bind TMED5 (negative logFC value) or TMED7 (positive logFC value). Proteins are sorted with logFC value and were analyzed using the Limma package in R.

**Table S5. Secretory pathway markers.** List of secretory pathway antibodies used throughout the manuscript in immunofluorescence experiments.

**Table S6. IP-MS of TMED7-MYC and TMED7 with TMED5CC-MYC.** Proteins that preferentially bind TMED7 compared to TMED7 with a TMED5 CC domain. Proteins (with gene names) are sorted with logFC value and were analyzed using the Limma package in R.

**Table S7. TMED antibodies.** List of antibodies detecting TMEDs used for Western blot and immunofluorescence throughout the manuscript.

## Notes

### Competing Interest Statement

The authors have declared no competing interest.

