## Supplementary figures and images for "A cargo receptor entrapment complex is a therapeutic node for genetically and clinically distinct proteinopathies"

### Supplemental Figures

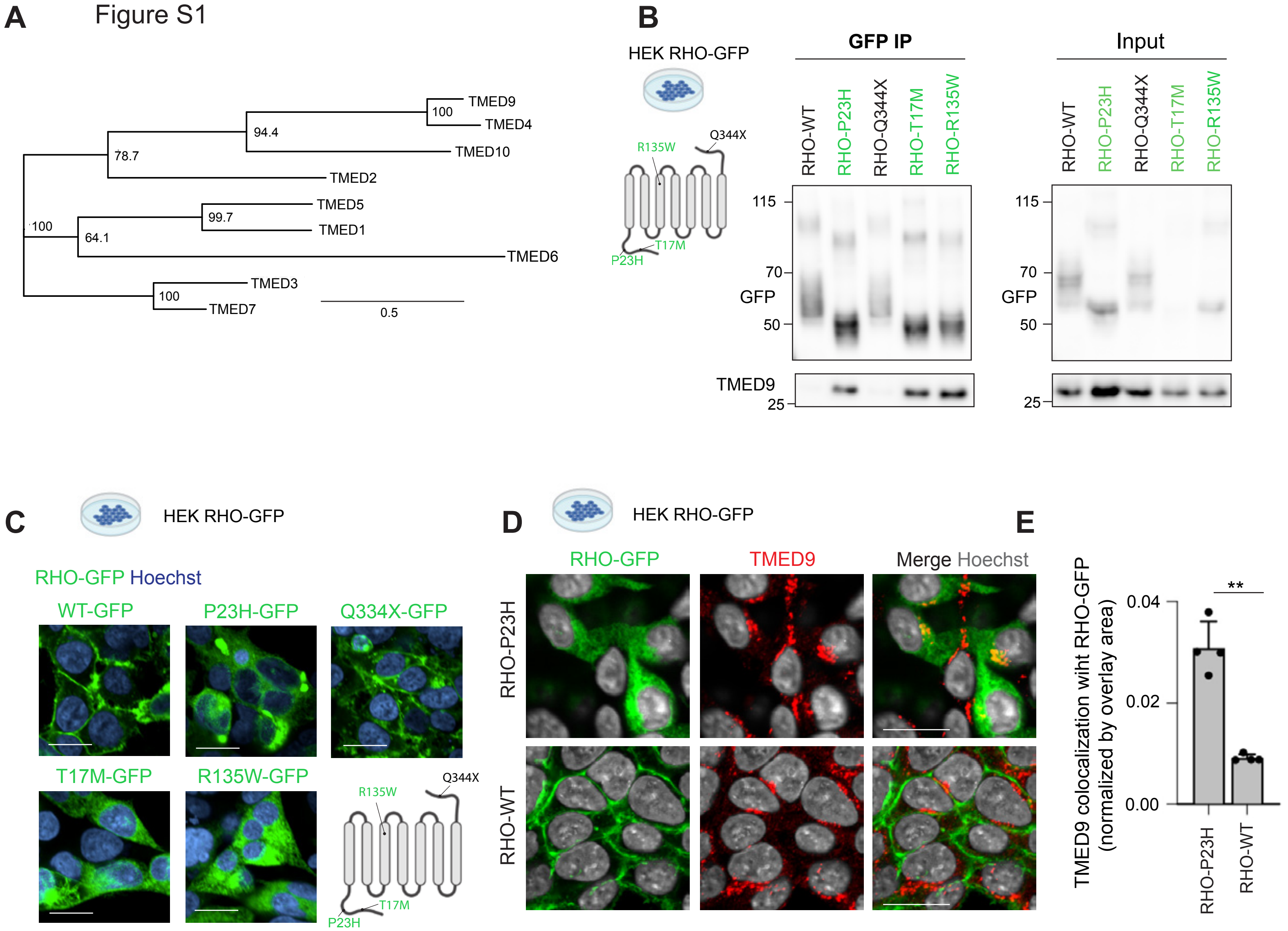

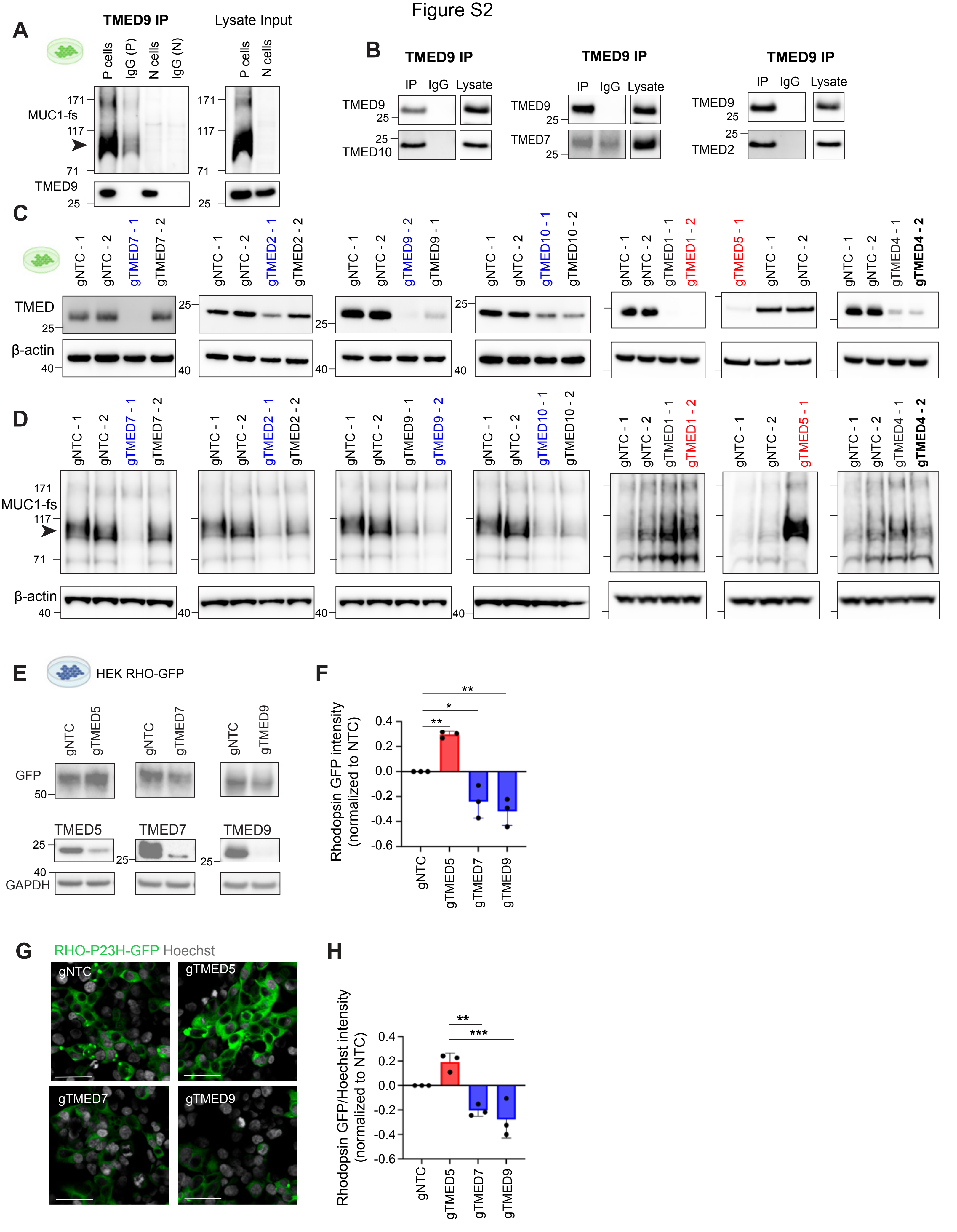

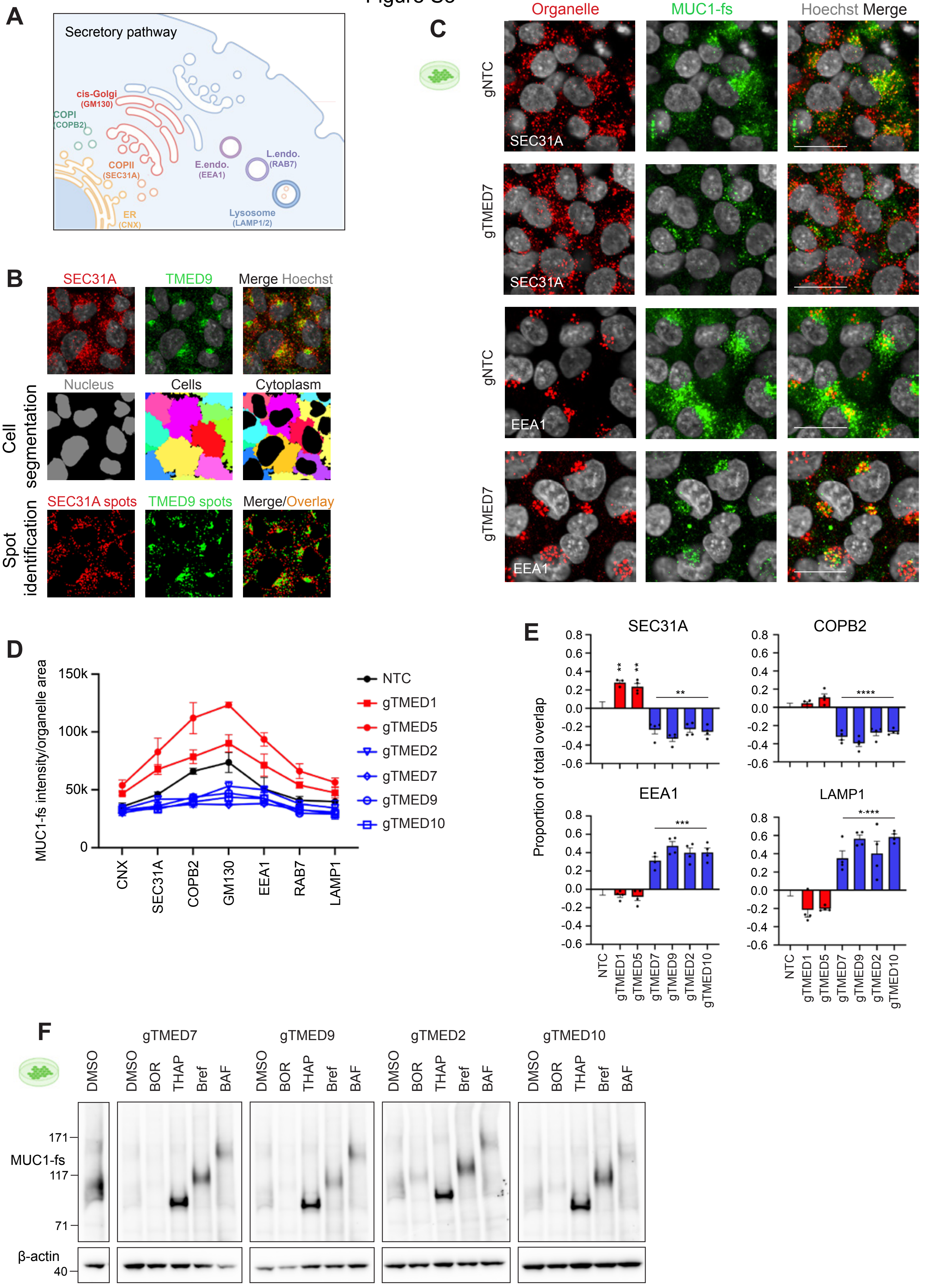

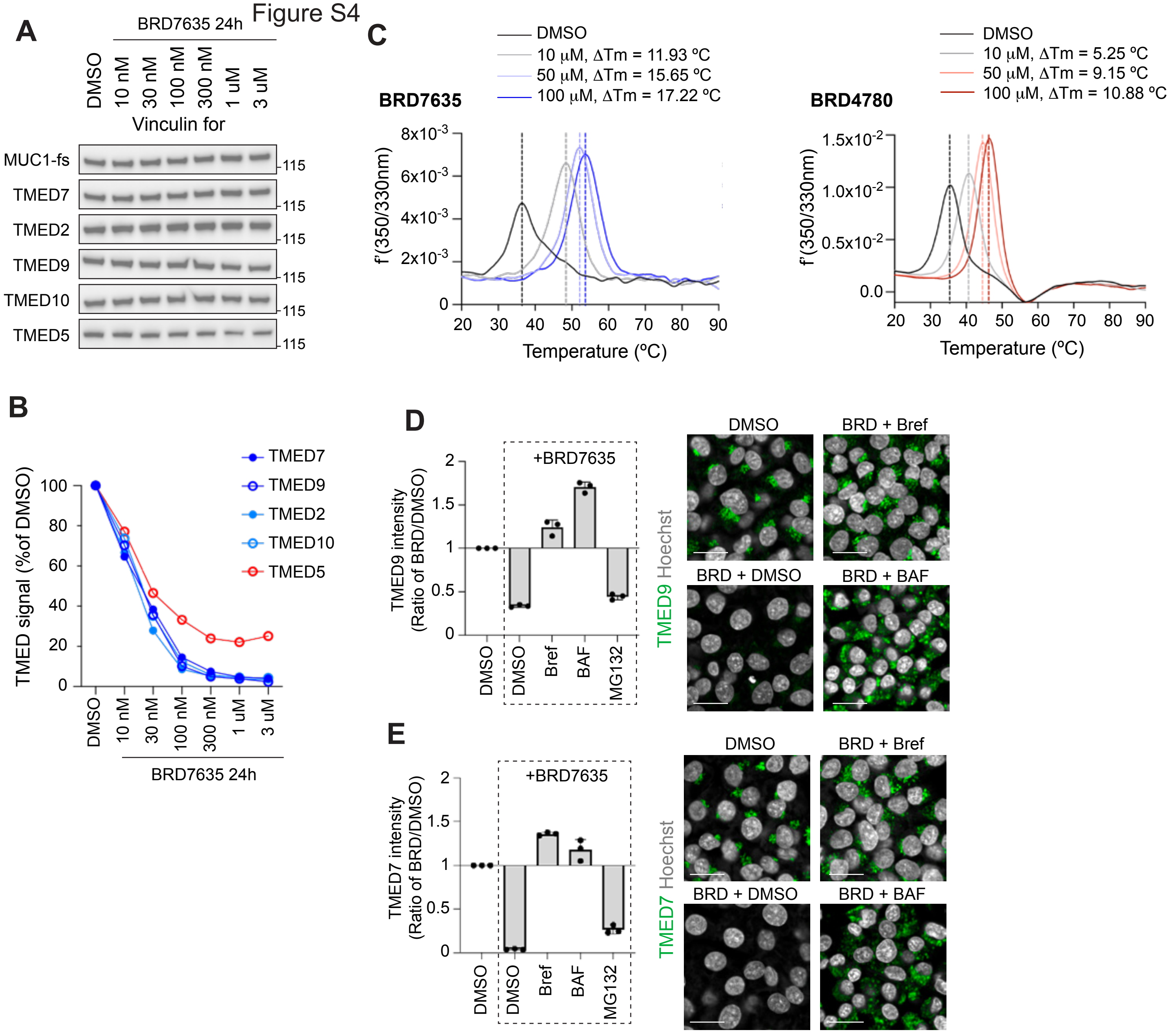

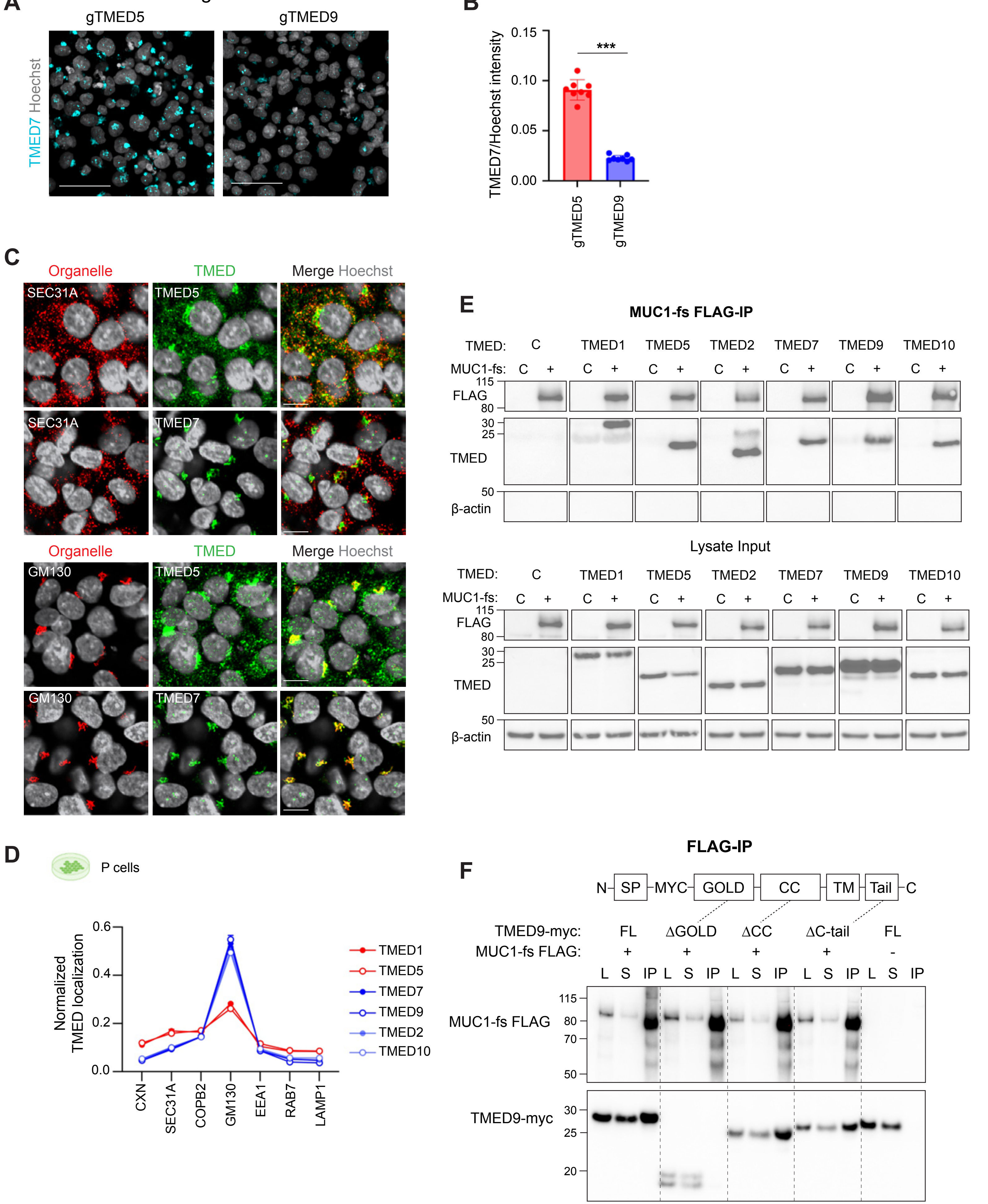

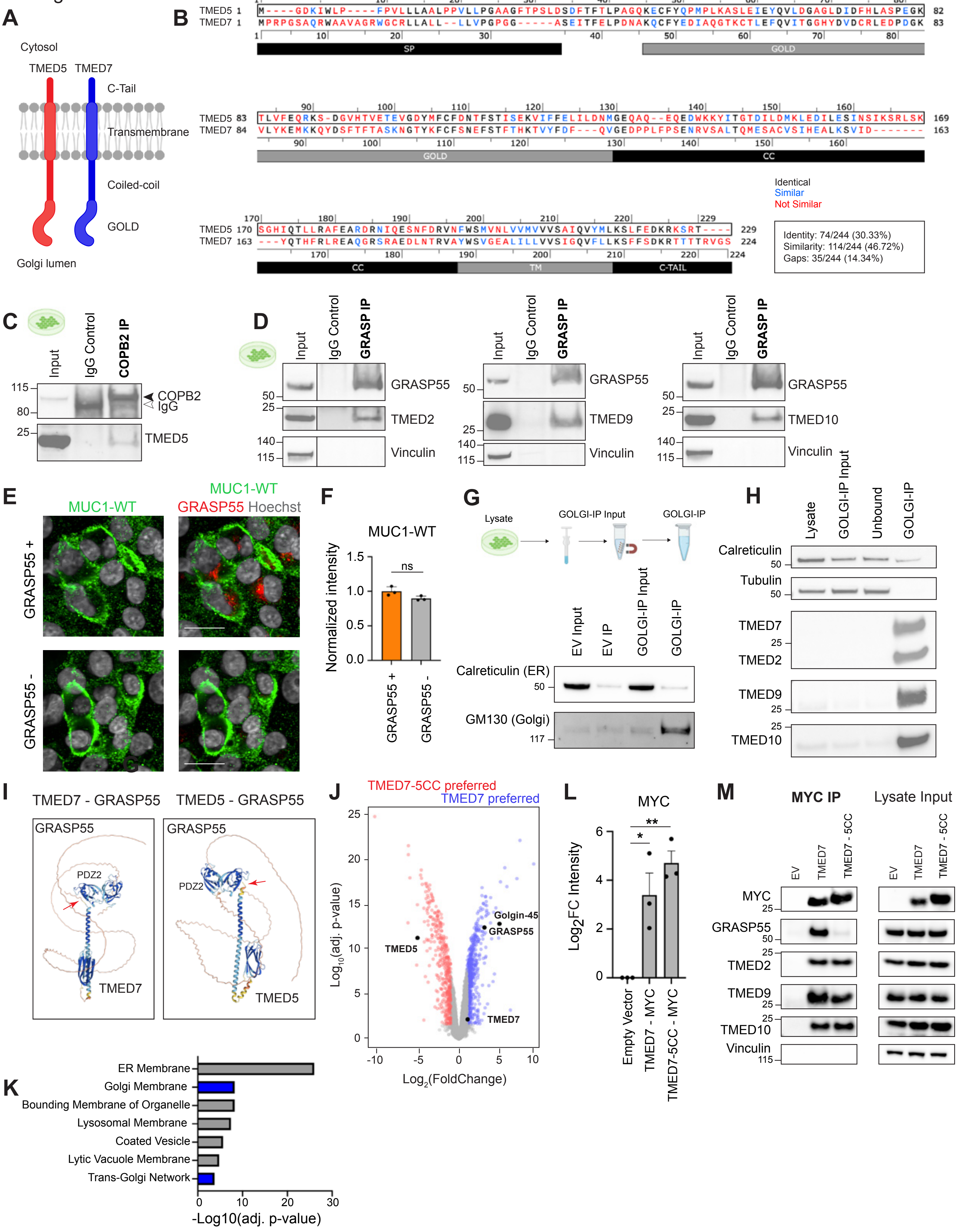

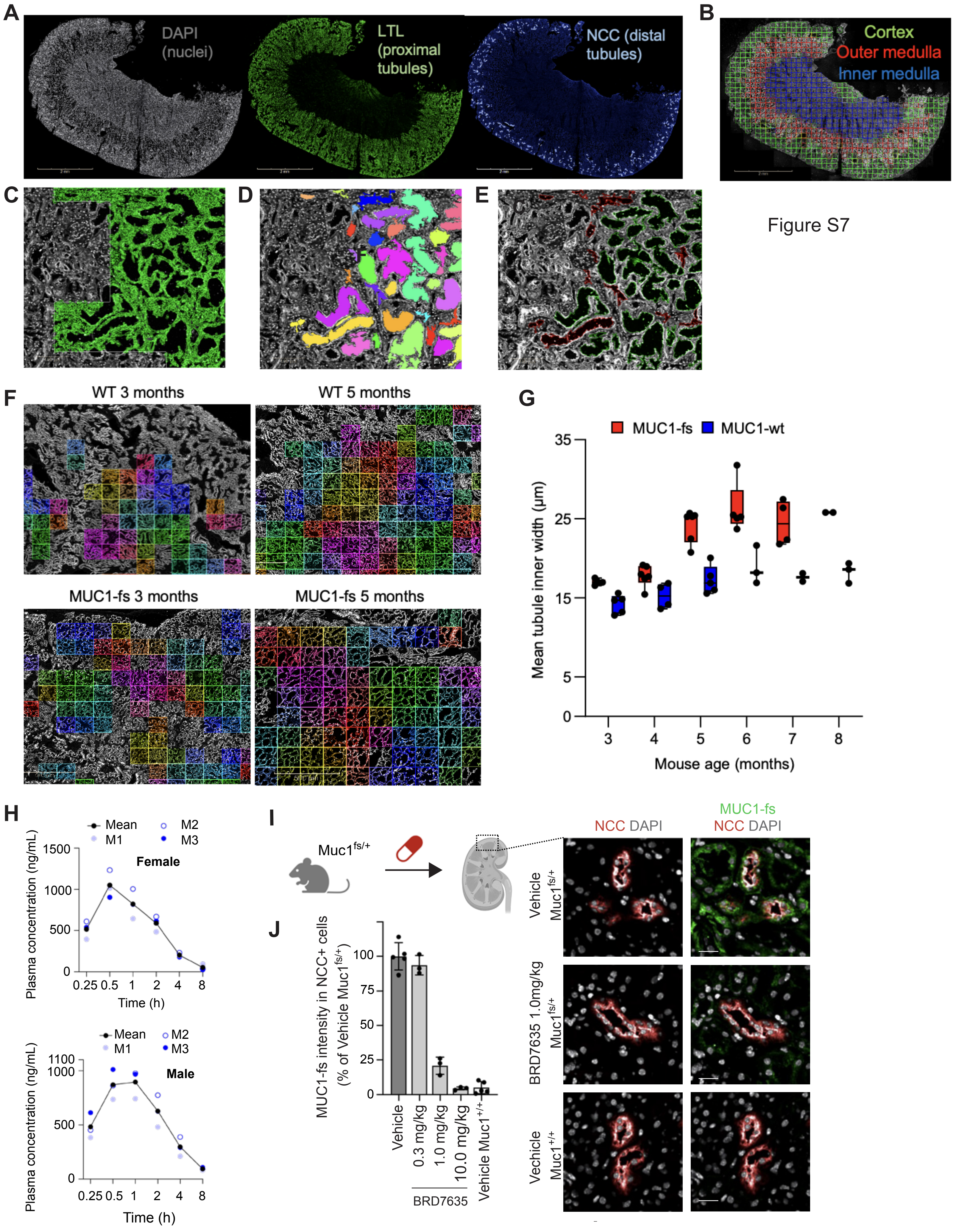
