## Supplemental Tables for "A cargo receptor entrapment complex is a therapeutic node for genetically and clinically distinct proteinopathies"

**Table S1. Significant interactors of MYC-TMED9 in HEK293T cells co-transfected with FLAG-MUC1-fs by $Log_2FC$.**

$Log_2FC$, $Log_2$ fold change in protein abundance between MYC-TMED9 and empty vector condition. Significant interactors chosen by adjusted p-value > 0.05.

| Interactor (Gene Name) | Log2FC | p-value | adjusted p-value |
|---|---|---|---|
| RAB5IF | 2.84 | 2.24E-07 | 7.76E-05 |
| ATP2A2 | 2.62367 | 4.94E-09 | 1.02E-05 |
| STARD3 | 2.527 | 1.02E-10 | 4.23E-07 |
| SC5D | 2.32967 | 1.00E-08 | 1.20E-05 |
| SAAL1 | 2.25233 | 1.35E-07 | 5.75E-05 |
| YIPF5 | 2.21567 | 2.87E-07 | 7.94E-05 |
| VIM | 2.145 | 1.15E-08 | 1.20E-05 |
| RANBP1 | 2.13967 | 1.67E-06 | 1.48E-04 |
| CLCN5 | 2.09833 | 1.15E-06 | 1.41E-04 |
| TMED9 | 1.945 | 9.59E-08 | 4.98E-05 |
| ATP1A1 | 1.933 | 5.78E-08 | 4.79E-05 |
| RPL38 | 1.91767 | 7.08E-06 | 3.15E-04 |
| ARL6IP6 | 1.90567 | 9.76E-07 | 1.40E-04 |
| TMED10 | 1.867 | 5.30E-06 | 2.65E-04 |
| SLC7A1 | 1.863 | 2.34E-06 | 1.76E-04 |
| TMED7 | 1.83033 | 1.49E-04 | 2.71E-03 |
| TNPO1 | 1.71667 | 3.61E-06 | 2.06E-04 |
| JAK3 | 1.667 | 2.46E-06 | 1.76E-04 |
| HACD3 | 1.65467 | 1.67E-06 | 1.48E-04 |
| ZMPSTE24 | 1.65233 | 4.30E-06 | 2.24E-04 |
| SELENOI | 1.63133 | 2.38E-05 | 7.31E-04 |
| EXOC7 | 1.62667 | 4.50E-07 | 1.10E-04 |
| XPO1 | 1.57333 | 2.45E-06 | 1.76E-04 |
| SPG11 | 1.51333 | 8.40E-05 | 1.72E-03 |
| ERGIC2 | 1.49667 | 1.16E-06 | 1.41E-04 |
| ERLEC1 | 1.48733 | 6.65E-05 | 1.47E-03 |
| EXOC5 | 1.474 | 8.18E-07 | 1.39E-04 |
| SET | 1.44467 | 2.64E-03 | 1.99E-02 |
| EXOC6 | 1.428 | 5.06E-07 | 1.17E-04 |
| IPO9 | 1.42767 | 4.41E-06 | 2.26E-04 |
| XPO6 | 1.42533 | 6.25E-06 | 2.91E-04 |
| CYBA | 1.42133 | 6.01E-07 | 1.31E-04 |
| VKORC1L1 | 1.409 | 1.64E-05 | 5.76E-04 |
| DYM | 1.386 | 9.49E-07 | 1.40E-04 |
| UBR3 | 1.37267 | 3.94E-07 | 1.02E-04 |
| RNF5 | 1.37033 | 8.08E-08 | 4.79E-05 |
| ERGIC3 | 1.34633 | 6.17E-06 | 2.91E-04 |
| LENG1 | 1.34133 | 1.49E-05 | 5.38E-04 |
| RABL3 | 1.32767 | 2.76E-07 | 7.94E-05 |

| | | | |
|---|---|---|---|
| YIF1B | 1.31267 | 6.35E-05 | 1.44E-03 |
| FBXO28 | 1.26467 | 2.40E-05 | 7.31E-04 |
| MZT2B | 1.262 | 6.32E-06 | 2.92E-04 |
| PRKDC | 1.25333 | 1.79E-07 | 6.74E-05 |
| TNPO3 | 1.251 | 6.57E-06 | 2.96E-04 |
| MANEA | 1.24133 | 1.28E-04 | 2.43E-03 |
| COPG2 | 1.23733 | 6.94E-08 | 4.79E-05 |
| CIP2A | 1.226 | 7.63E-07 | 1.38E-04 |
| NPEPPS | 1.22267 | 2.83E-05 | 8.33E-04 |
| ATXN10 | 1.22033 | 3.03E-05 | 8.68E-04 |
| INTS12 | 1.209 | 3.28E-06 | 1.99E-04 |
| STT3A | 1.209 | 1.74E-06 | 1.49E-04 |
| RNF185 | 1.20733 | 1.68E-05 | 5.81E-04 |
| SLC30A9 | 1.18733 | 8.67E-07 | 1.39E-04 |
| PPP4R3B | 1.182 | 3.19E-06 | 1.98E-04 |
| ARCN1 | 1.18 | 7.38E-06 | 3.22E-04 |
| CSE1L | 1.17733 | 4.36E-05 | 1.10E-03 |
| ATP1B3 | 1.17167 | 8.01E-05 | 1.66E-03 |
| FAM114A2 | 1.16367 | 9.15E-05 | 1.86E-03 |
| PKDREJ | 1.16333 | 2.18E-05 | 6.91E-04 |
| TMED4 | 1.15367 | 1.35E-06 | 1.48E-04 |
| GLMN | 1.14833 | 4.65E-05 | 1.13E-03 |
| VPS50 | 1.144 | 1.52E-04 | 2.73E-03 |
| SLC30A7 | 1.144 | 1.34E-04 | 2.53E-03 |
| PPIB | 1.143 | 3.80E-05 | 1.02E-03 |
| KIF14 | 1.14133 | 1.39E-07 | 5.75E-05 |
| CU | 1.14 | 2.58E-07 | 7.94E-05 |
| HOOK2 | 1.13933 | 3.83E-06 | 2.06E-04 |
| VDAC2 | 1.139 | 7.46E-06 | 3.22E-04 |
| TNPO2 | 1.13867 | 5.55E-06 | 2.68E-04 |
| EXOC2 | 1.13733 | 1.97E-06 | 1.63E-04 |
| UBE4A | 1.13667 | 5.56E-06 | 2.68E-04 |
| TUBGCP2 | 1.121 | 1.39E-05 | 5.18E-04 |
| HOOK3 | 1.11867 | 5.00E-06 | 2.53E-04 |
| RPN1 | 1.116 | 1.49E-06 | 1.48E-04 |
| XPOT | 1.11467 | 2.57E-05 | 7.68E-04 |
| XPO5 | 1.10867 | 5.02E-05 | 1.20E-03 |
| HEATR3 | 1.10867 | 9.16E-06 | 3.88E-04 |
| NGLY1 | 1.108 | 1.96E-03 | 1.64E-02 |
| NCAPH | 1.10767 | 1.64E-06 | 1.48E-04 |
| RINT1 | 1.106 | 6.94E-07 | 1.33E-04 |
| LMAN1 | 1.101 | 5.11E-04 | 6.02E-03 |
| VPS8 | 1.09633 | 1.31E-05 | 5.04E-04 |
| RCN1 | 1.095 | 2.22E-03 | 1.76E-02 |
| RLIM | 1.09367 | 1.55E-06 | 1.48E-04 |
| TMCO1 | 1.093 | 1.45E-06 | 1.48E-04 |
| NCAPD2 | 1.09 | 2.06E-06 | 1.65E-04 |
| TTYH3 | 1.08833 | 3.33E-06 | 1.99E-04 |
| ASS1 | 1.08033 | 1.49E-05 | 5.38E-04 |

| | | | |
|---|---|---|---|
| EI24 | 1.08033 | 2.99E-06 | 1.91E-04 |
| EXOC8 | 1.077 | 6.15E-05 | 1.41E-03 |
| TBC1D8B | 1.06033 | 8.08E-06 | 3.46E-04 |
| SURF4 | 1.05867 | 2.76E-06 | 1.83E-04 |
| TMEM161A | 1.049 | 1.37E-05 | 5.16E-04 |
| DDOST | 1.046 | 1.51E-05 | 5.39E-04 |
| UBE3B | 1.042 | 2.16E-05 | 6.90E-04 |
| ZDHHC13 | 1.03367 | 4.31E-06 | 2.24E-04 |
| CAND1 | 1.03033 | 4.91E-05 | 1.18E-03 |
| XPO4 | 1.02933 | 1.00E-04 | 1.97E-03 |
| ANO10 | 1.029 | 1.73E-04 | 2.94E-03 |
| VAPA | 1.023 | 1.62E-05 | 5.74E-04 |
| BTAF1 | 1.02233 | 1.69E-05 | 5.81E-04 |
| MKS1 | 1.01333 | 5.37E-04 | 6.24E-03 |
| COG6 | 1.011 | 5.48E-06 | 2.68E-04 |
| C19orf25 | 1.01067 | 2.10E-06 | 1.65E-04 |
| YTHDF3 | 1.00733 | 8.62E-03 | 4.41E-02 |
| COPB2 | 1.00733 | 4.44E-05 | 1.10E-03 |
| NCDN | 0.99633 | 4.74E-05 | 1.15E-03 |
| HSPA5 | 0.995 | 7.32E-05 | 1.56E-03 |
| HEATR1 | 0.99433 | 3.88E-06 | 2.06E-04 |
| C1orf112 | 0.98967 | 2.12E-04 | 3.29E-03 |
| STT3B | 0.985 | 3.85E-06 | 2.06E-04 |
| NDUFS6 | 0.98 | 3.28E-04 | 4.46E-03 |
| OXA1L | 0.97633 | 1.60E-06 | 1.48E-04 |
| ARMC9 | 0.97067 | 9.54E-05 | 1.91E-03 |
| SMC4 | 0.96433 | 3.72E-06 | 2.06E-04 |
| COPE | 0.96367 | 6.33E-07 | 1.31E-04 |
| TMED5 | 0.963 | 4.27E-04 | 5.31E-03 |
| SMC2 | 0.95967 | 1.06E-06 | 1.41E-04 |
| SMG9 | 0.95367 | 2.78E-06 | 1.83E-04 |
| IPO7 | 0.95 | 1.94E-05 | 6.36E-04 |
| BZW2 | 0.94767 | 1.80E-04 | 3.01E-03 |
| KPNB1 | 0.941 | 7.03E-07 | 1.33E-04 |
| SGSM3 | 0.93867 | 2.11E-06 | 1.65E-04 |
| DERL1 | 0.93267 | 3.19E-05 | 8.95E-04 |
| EXOC6B | 0.92967 | 4.02E-05 | 1.06E-03 |
| SLC39A14 | 0.92867 | 3.02E-03 | 2.18E-02 |
| TBC1D9 | 0.91933 | 2.14E-05 | 6.87E-04 |
| MTFR2 | 0.91767 | 1.06E-04 | 2.07E-03 |
| CANX | 0.917 | 1.61E-03 | 1.41E-02 |
| TBL2 | 0.91333 | 2.57E-06 | 1.81E-04 |
| NUP210 | 0.909 | 3.49E-05 | 9.41E-04 |
| ARFGAP2 | 0.90833 | 1.23E-03 | 1.17E-02 |
| INTS1 | 0.90633 | 5.61E-06 | 2.68E-04 |
| HMOX2 | 0.906 | 1.95E-05 | 6.36E-04 |
| NUP93 | 0.90267 | 6.56E-06 | 2.96E-04 |
| UBE3C | 0.90233 | 1.30E-06 | 1.48E-04 |
| NDUFA10 | 0.902 | 5.30E-05 | 1.25E-03 |

| | | | |
|---|---|---|---|
| DNAJC30 | 0.902 | 1.37E-06 | 1.48E-04 |
| PTDSS1 | 0.89933 | 1.43E-04 | 2.63E-03 |
| EMD | 0.89233 | 1.68E-06 | 1.48E-04 |
| CIAO1 | 0.89133 | 1.84E-04 | 3.04E-03 |
| DYNC2LI1 | 0.89033 | 6.10E-03 | 3.49E-02 |
| TMED1 | 0.879 | 2.10E-04 | 3.29E-03 |
| MON2 | 0.87833 | 9.06E-07 | 1.39E-04 |
| PKD2 | 0.87633 | 8.05E-05 | 1.66E-03 |
| NCAPG | 0.87633 | 1.45E-05 | 5.36E-04 |
| RMND1 | 0.87567 | 2.29E-04 | 3.46E-03 |
| ATP1A3 | 0.87333 | 3.16E-04 | 4.35E-03 |
| GOLGA5 | 0.869 | 8.04E-05 | 1.66E-03 |
| DOCK1 | 0.86667 | 4.96E-04 | 5.95E-03 |
| GCN1 | 0.86633 | 4.27E-05 | 1.09E-03 |
| ABHD3 | 0.86067 | 1.47E-05 | 5.38E-04 |
| ATP1B1 | 0.858 | 3.10E-05 | 8.74E-04 |
| RHBDD1 | 0.85733 | 1.58E-04 | 2.79E-03 |
| TMEM109 | 0.84967 | 1.04E-05 | 4.22E-04 |
| DNAJC16 | 0.84867 | 2.02E-04 | 3.22E-03 |
| SUN2 | 0.848 | 2.65E-04 | 3.84E-03 |
| YIPF4 | 0.848 | 2.18E-04 | 3.34E-03 |
| EVI5 | 0.847 | 1.18E-06 | 1.41E-04 |
| RB1 | 0.846 | 1.76E-06 | 1.49E-04 |
| GPR180 | 0.84467 | 1.95E-05 | 6.36E-04 |
| DOCK4 | 0.84367 | 3.06E-06 | 1.93E-04 |
| EXOC3 | 0.83867 | 7.39E-05 | 1.56E-03 |
| DNAJC3 | 0.836 | 2.14E-03 | 1.74E-02 |
| HLA-B | 0.83267 | 2.87E-05 | 8.34E-04 |
| TIMELESS | 0.83033 | 1.05E-06 | 1.41E-04 |
| EXOC1 | 0.82833 | 1.46E-06 | 1.48E-04 |
| ATM | 0.82767 | 6.53E-05 | 1.46E-03 |
| SOGA1 | 0.826 | 5.33E-03 | 3.18E-02 |
| FAM96B | 0.826 | 4.05E-05 | 1.06E-03 |
| CLCC1 | 0.82567 | 1.62E-04 | 2.81E-03 |
| ST8 | 0.814 | 8.43E-04 | 8.95E-03 |
| SLC1A5 | 0.81367 | 1.02E-03 | 1.03E-02 |
| TRAM1 | 0.81167 | 3.26E-05 | 9.08E-04 |
| NUP205 | 0.809 | 7.13E-06 | 3.15E-04 |
| SDF2L1 | 0.80767 | 4.58E-05 | 1.13E-03 |
| NEFM | 0.79633 | 1.97E-04 | 3.16E-03 |
| PGRMC1 | 0.79633 | 2.23E-06 | 1.71E-04 |
| WFS1 | 0.79167 | 1.49E-03 | 1.33E-02 |
| AMFR | 0.79067 | 4.66E-04 | 5.64E-03 |
| FANCD2 | 0.78933 | 2.78E-06 | 1.83E-04 |
| COPG1 | 0.787 | 5.38E-04 | 6.24E-03 |
| COPB1 | 0.78233 | 7.26E-05 | 1.56E-03 |
| MZT2A | 0.78033 | 3.06E-04 | 4.28E-03 |
| SEC61A1 | 0.78 | 2.85E-06 | 1.85E-04 |
| RAN | 0.77767 | 2.42E-05 | 7.33E-04 |

| | | | |
|---|---|---|---|
| COG8 | 0.77733 | 4.51E-05 | 1.11E-03 |
| ITPRIPL1 | 0.77667 | 7.72E-04 | 8.37E-03 |
| NLR | 0.77667 | 1.72E-04 | 2.94E-03 |
| NR6A1 | 0.77567 | 2.93E-04 | 4.12E-03 |
| DAB2IP | 0.77267 | 2.72E-06 | 1.83E-04 |
| SLC39A6 | 0.77067 | 1.83E-05 | 6.13E-04 |
| IPO5 | 0.76733 | 3.47E-04 | 4.59E-03 |
| UBAC2 | 0.767 | 9.05E-07 | 1.39E-04 |
| VDAC3 | 0.764 | 6.53E-05 | 1.46E-03 |
| COPZ1 | 0.75867 | 1.50E-03 | 1.34E-02 |
| NDUFA7 | 0.755 | 1.48E-04 | 2.70E-03 |
| VRK2 | 0.75433 | 1.70E-05 | 5.81E-04 |
| TBC1D9B | 0.75267 | 1.01E-05 | 4.21E-04 |
| SEC23IP | 0.74933 | 2.86E-04 | 4.06E-03 |
| SEL1L | 0.74733 | 7.02E-05 | 1.53E-03 |
| ZW10 | 0.746 | 2.69E-04 | 3.88E-03 |
| GANAB | 0.745 | 1.14E-06 | 1.41E-04 |
| CAND2 | 0.741 | 1.93E-04 | 3.15E-03 |
| TMED2 | 0.73467 | 8.57E-03 | 4.41E-02 |
| OTOA | 0.731 | 3.37E-06 | 1.99E-04 |
| OS9 | 0.729 | 1.10E-03 | 1.08E-02 |
| TM9SF3 | 0.728 | 1.02E-03 | 1.03E-02 |
| PHKA1 | 0.72133 | 2.27E-05 | 7.02E-04 |
| EXOC4 | 0.72133 | 2.43E-06 | 1.76E-04 |
| ORC2 | 0.72033 | 2.16E-04 | 3.33E-03 |
| B4GAT1 | 0.71733 | 2.96E-04 | 4.16E-03 |
| PIGS | 0.71667 | 1.15E-05 | 4.60E-04 |
| NCAPD3 | 0.71633 | 3.41E-06 | 1.99E-04 |
| DNAAF5 | 0.71233 | 6.56E-03 | 3.71E-02 |
| PGM3 | 0.70833 | 4.19E-05 | 1.07E-03 |
| CEPT1 | 0.70233 | 1.57E-04 | 2.79E-03 |
| COG5 | 0.701 | 3.33E-05 | 9.16E-04 |
| SMPD4 | 0.69933 | 2.75E-04 | 3.94E-03 |
| MARS | 0.69733 | 2.71E-05 | 8.03E-04 |
| NUP160 | 0.69667 | 3.72E-06 | 2.06E-04 |
| TMX2 | 0.69533 | 2.88E-03 | 2.12E-02 |
| TMEM11 | 0.693 | 1.09E-04 | 2.11E-03 |
| B3GAT3 | 0.69233 | 3.37E-05 | 9.22E-04 |
| ATP2B1 | 0.69067 | 1.27E-05 | 4.96E-04 |
| HIP1R | 0.68633 | 9.51E-03 | 4.68E-02 |
| LRPPRC | 0.68633 | 5.81E-05 | 1.34E-03 |
| LEMD2 | 0.686 | 1.38E-04 | 2.58E-03 |
| USP8 | 0.68467 | 1.28E-03 | 1.19E-02 |
| AKAP11 | 0.68467 | 3.07E-05 | 8.73E-04 |
| KIAA0355 | 0.684 | 8.52E-03 | 4.40E-02 |
| APMAP | 0.68333 | 1.84E-03 | 1.56E-02 |
| TTC12 | 0.683 | 1.33E-03 | 1.22E-02 |
| HLA-C | 0.683 | 1.51E-04 | 2.73E-03 |
| LBR | 0.682 | 2.22E-05 | 6.94E-04 |

| | | | |
|---|---|---|---|
| DNAJB12 | 0.68167 | 1.24E-05 | 4.91E-04 |
| ATP11B | 0.68067 | 3.23E-04 | 4.42E-03 |
| TMEM168 | 0.679 | 8.61E-03 | 4.41E-02 |
| THADA | 0.677 | 1.85E-04 | 3.04E-03 |
| ELMO2 | 0.67667 | 7.88E-05 | 1.65E-03 |
| ALDH3A2 | 0.67633 | 3.30E-05 | 9.14E-04 |
| GOSR1 | 0.67367 | 2.07E-04 | 3.28E-03 |
| MUC1-fs | 0.668 | 6.31E-03 | 3.61E-02 |
| SCCPDH | 0.66133 | 6.57E-05 | 1.46E-03 |
| CLCN7 | 0.66133 | 5.66E-05 | 1.32E-03 |
| NUP43 | 0.65833 | 2.12E-04 | 3.29E-03 |
| MTOR | 0.65667 | 2.08E-04 | 3.28E-03 |
| LTN1 | 0.652 | 4.41E-05 | 1.10E-03 |
| STX5 | 0.65 | 4.94E-04 | 5.94E-03 |
| NUP85 | 0.64867 | 3.83E-06 | 2.06E-04 |
| ERLIN2 | 0.64733 | 5.68E-04 | 6.57E-03 |
| EMC2 | 0.64567 | 1.24E-04 | 2.37E-03 |
| COPA | 0.64167 | 5.34E-05 | 1.25E-03 |
| DYNC2H1 | 0.63267 | 1.46E-04 | 2.68E-03 |
| ATP2B4 | 0.63033 | 3.67E-04 | 4.75E-03 |
| KNTC1 | 0.62767 | 3.09E-04 | 4.30E-03 |
| RABGAP1 | 0.627 | 1.25E-03 | 1.17E-02 |
| CDK12 | 0.62633 | 1.81E-05 | 6.13E-04 |
| IMMT | 0.62567 | 2.86E-05 | 8.34E-04 |
| MTCH1 | 0.62467 | 6.67E-05 | 1.47E-03 |
| FNDC3B | 0.624 | 2.42E-04 | 3.60E-03 |
| BZW1 | 0.624 | 1.76E-04 | 2.97E-03 |
| CHCHD3 | 0.623 | 1.60E-04 | 2.81E-03 |
| CCT3 | 0.62233 | 2.45E-05 | 7.37E-04 |
| TTC27 | 0.62067 | 4.02E-05 | 1.06E-03 |
| PGRMC2 | 0.618 | 1.83E-04 | 3.04E-03 |
| HOOK1 | 0.61633 | 1.53E-04 | 2.74E-03 |
| MTCH2 | 0.61467 | 6.20E-05 | 1.41E-03 |
| HLA-A | 0.61367 | 4.47E-03 | 2.84E-02 |
| GBF1 | 0.60967 | 6.91E-04 | 7.59E-03 |
| CLTA | 0.60567 | 1.24E-04 | 2.37E-03 |
| NDUFV1 | 0.60433 | 1.68E-03 | 1.46E-02 |
| COG7 | 0.60267 | 8.28E-04 | 8.85E-03 |
| ATP6V1H | 0.602 | 3.72E-03 | 2.53E-02 |
| TMEM9 | 0.602 | 8.30E-04 | 8.86E-03 |
| SAMM50 | 0.597 | 8.20E-04 | 8.80E-03 |
| ATP6V1A | 0.59633 | 1.61E-04 | 2.81E-03 |
| TMEM201 | 0.595 | 1.92E-03 | 1.61E-02 |
| INA | 0.592 | 2.58E-04 | 3.75E-03 |
| TFRC | 0.59033 | 2.89E-03 | 2.12E-02 |
| PIK3R2 | 0.58967 | 3.33E-04 | 4.47E-03 |
| CARM1 | 0.58667 | 5.03E-03 | 3.07E-02 |
| DHCR7 | 0.58233 | 3.91E-03 | 2.61E-02 |
| LRBA | 0.58167 | 6.32E-04 | 7.18E-03 |

| | | | |
|---|---|---|---|
| PPP3CA | 0.58033 | 3.41E-03 | 2.39E-02 |
| PIGK | 0.58 | 4.20E-04 | 5.24E-03 |
| THEM6 | 0.58 | 4.14E-05 | 1.07E-03 |
| KIAA0391 | 0.58 | 3.90E-05 | 1.04E-03 |
| SACM1L | 0.573 | 2.21E-05 | 6.94E-04 |
| BSG | 0.57267 | 9.02E-04 | 9.39E-03 |
| TBC1D17 | 0.571 | 1.03E-05 | 4.22E-04 |
| SMOC1 | 0.57 | 5.14E-05 | 1.22E-03 |
| COQ8A | 0.56667 | 2.38E-04 | 3.58E-03 |
| NOP9 | 0.566 | 7.25E-05 | 1.56E-03 |
| MAGED1 | 0.56033 | 9.43E-06 | 3.95E-04 |
| CERS1 | 0.55833 | 5.09E-04 | 6.02E-03 |
| HSP90B1 | 0.552 | 1.28E-05 | 4.96E-04 |
| UCK2 | 0.55 | 9.10E-04 | 9.44E-03 |
| KTN1 | 0.549 | 2.48E-04 | 3.65E-03 |
| FASTKD5 | 0.548 | 4.50E-04 | 5.56E-03 |
| TMEM57 | 0.54533 | 1.75E-04 | 2.97E-03 |
| TELO2 | 0.54433 | 1.43E-04 | 2.63E-03 |
| SLIRP | 0.543 | 2.23E-04 | 3.39E-03 |
| KPNA2 | 0.542 | 2.23E-04 | 3.39E-03 |
| EPRS | 0.54033 | 1.56E-04 | 2.78E-03 |
| UTP20 | 0.53833 | 6.59E-04 | 7.38E-03 |
| CCDC47 | 0.537 | 9.53E-05 | 1.91E-03 |
| IRS4 | 0.537 | 2.06E-05 | 6.67E-04 |
| DNAJC11 | 0.535 | 3.03E-05 | 8.68E-04 |
| GALNT2 | 0.53233 | 3.31E-04 | 4.46E-03 |
| C6orf203 | 0.52733 | 6.62E-04 | 7.39E-03 |
| RPN2 | 0.52467 | 3.50E-03 | 2.43E-02 |
| SLC35B2 | 0.52333 | 1.20E-03 | 1.14E-02 |
| EEF1E1 | 0.52267 | 3.66E-04 | 4.75E-03 |
| GPX8 | 0.52033 | 1.17E-03 | 1.12E-02 |
| HSP90AB1 | 0.51867 | 2.56E-04 | 3.74E-03 |
| UBE2G2 | 0.51633 | 9.59E-04 | 9.83E-03 |
| SRPRB | 0.51633 | 1.63E-04 | 2.83E-03 |
| DHCR24 | 0.515 | 1.97E-04 | 3.16E-03 |
| DDX20 | 0.51467 | 3.86E-04 | 4.93E-03 |
| TFIP11 | 0.512 | 1.82E-04 | 3.04E-03 |
| COG4 | 0.50767 | 3.44E-05 | 9.33E-04 |
| ATP2C1 | 0.50233 | 6.12E-04 | 7.00E-03 |
| UNC45A | 0.50033 | 4.54E-04 | 5.58E-03 |
| RANBP6 | 0.49733 | 3.79E-04 | 4.86E-03 |
| SYNGAP1 | 0.49667 | 8.27E-03 | 4.31E-02 |
| DOCK3 | 0.495 | 3.69E-04 | 4.77E-03 |
| STK11IP | 0.49033 | 1.00E-03 | 1.02E-02 |
| PAXBP1 | 0.49033 | 5.71E-05 | 1.32E-03 |
| NDUFA2 | 0.49 | 6.48E-04 | 7.27E-03 |
| VPS41 | 0.48867 | 2.55E-04 | 3.74E-03 |
| FANCI | 0.48833 | 4.63E-04 | 5.64E-03 |
| MLST8 | 0.48567 | 3.97E-04 | 5.03E-03 |

| | | | |
|---|---|---|---|
| DIAPH3 | 0.48267 | 3.17E-04 | 4.35E-03 |
| IARS | 0.48267 | 1.78E-04 | 2.99E-03 |
| EFHD2 | 0.482 | 5.81E-04 | 6.68E-03 |
| HSD17B12 | 0.48033 | 1.87E-03 | 1.58E-02 |
| FANCA | 0.47533 | 4.36E-03 | 2.80E-02 |
| VPS16 | 0.475 | 4.65E-04 | 5.64E-03 |
| ESYT1 | 0.47133 | 2.90E-04 | 4.10E-03 |
| STAT1 | 0.469 | 3.31E-04 | 4.46E-03 |
| CLTC | 0.46833 | 1.38E-03 | 1.25E-02 |
| ICMT | 0.46767 | 2.59E-03 | 1.96E-02 |
| USP34 | 0.465 | 9.25E-04 | 9.55E-03 |
| NDUFV2 | 0.46333 | 7.56E-04 | 8.21E-03 |
| PRKAR1A | 0.46267 | 3.90E-03 | 2.61E-02 |
| KIAA0368 | 0.462 | 7.76E-03 | 4.11E-02 |
| ORC5 | 0.46167 | 1.14E-03 | 1.11E-02 |
| PDCD6 | 0.46133 | 2.41E-03 | 1.86E-02 |
| NDUFS2 | 0.46067 | 6.96E-03 | 3.82E-02 |
| PIGA | 0.46 | 5.26E-04 | 6.15E-03 |
| BRAT1 | 0.458 | 1.44E-03 | 1.29E-02 |
| PDS5A | 0.456 | 1.61E-03 | 1.41E-02 |
| NUP133 | 0.45233 | 1.96E-03 | 1.64E-02 |
| NEK6 | 0.45067 | 4.13E-04 | 5.19E-03 |
| VAC14 | 0.44967 | 5.01E-04 | 5.97E-03 |
| NELFB | 0.44967 | 2.46E-04 | 3.63E-03 |
| ALG1 | 0.448 | 2.21E-03 | 1.76E-02 |
| TCP1 | 0.448 | 1.26E-03 | 1.18E-02 |
| DYNC1LI1 | 0.447 | 1.36E-04 | 2.56E-03 |
| AUP1 | 0.44333 | 5.30E-04 | 6.18E-03 |
| USP13 | 0.44133 | 3.77E-03 | 2.55E-02 |
| TOMM22 | 0.441 | 2.38E-03 | 1.85E-02 |
| OPA1 | 0.44067 | 1.14E-03 | 1.11E-02 |
| NDUFA5 | 0.44033 | 5.45E-03 | 3.22E-02 |
| NEFL | 0.44033 | 5.80E-04 | 6.68E-03 |
| APBB2 | 0.43967 | 9.37E-05 | 1.90E-03 |
| NDUFA13 | 0.43867 | 3.76E-04 | 4.84E-03 |
| HAT1 | 0.43867 | 1.46E-04 | 2.68E-03 |
| PRAF2 | 0.43733 | 5.67E-03 | 3.32E-02 |
| CCT8 | 0.437 | 2.76E-04 | 3.94E-03 |
| PRMT3 | 0.43567 | 1.70E-03 | 1.47E-02 |
| DNAJC7 | 0.43433 | 1.30E-03 | 1.20E-02 |
| FASTKD1 | 0.43367 | 8.45E-03 | 4.37E-02 |
| PHGDH | 0.433 | 2.79E-03 | 2.06E-02 |
| ASPH | 0.433 | 2.87E-04 | 4.07E-03 |
| HEATR6 | 0.43267 | 2.37E-03 | 1.85E-02 |
| SSR1 | 0.43233 | 6.63E-03 | 3.72E-02 |
| IPO11 | 0.43133 | 1.69E-03 | 1.46E-02 |
| SYMPK | 0.431 | 2.35E-03 | 1.85E-02 |
| SMG8 | 0.43033 | 6.85E-04 | 7.57E-03 |
| NF1 | 0.429 | 1.72E-04 | 2.94E-03 |

| | | | |
|---|---|---|---|
| MMS19 | 0.42867 | 2.64E-03 | 1.99E-02 |
| NDUFA9 | 0.42567 | 2.25E-04 | 3.40E-03 |
| KDM4F | 0.42533 | 3.36E-03 | 2.36E-02 |
| TANGO6 | 0.422 | 8.32E-03 | 4.33E-02 |
| GK | 0.42167 | 3.97E-03 | 2.62E-02 |
| PPP2R1B | 0.42133 | 8.98E-03 | 4.53E-02 |
| WDR75 | 0.41967 | 3.15E-04 | 4.35E-03 |
| NUP188 | 0.41967 | 1.94E-04 | 3.15E-03 |
| VPS18 | 0.418 | 4.55E-03 | 2.86E-02 |
| CCT4 | 0.41733 | 1.11E-04 | 2.15E-03 |
| CASKIN2 | 0.41667 | 2.52E-03 | 1.92E-02 |
| ARHGAP29 | 0.41633 | 9.55E-05 | 1.91E-03 |
| TBC1D15 | 0.41533 | 8.52E-04 | 9.02E-03 |
| CDKAL1 | 0.41333 | 1.54E-03 | 1.36E-02 |
| DNAJB2 | 0.41233 | 6.69E-03 | 3.72E-02 |
| PDE3B | 0.412 | 4.03E-03 | 2.65E-02 |
| JPH1 | 0.41133 | 3.26E-04 | 4.46E-03 |
| NDUFS3 | 0.411 | 3.61E-03 | 2.48E-02 |
| SELENOK | 0.40733 | 1.02E-03 | 1.03E-02 |
| NR2F2 | 0.407 | 9.52E-03 | 4.68E-02 |
| ANAPC10 | 0.405 | 2.20E-03 | 1.75E-02 |
| AIMP1 | 0.40333 | 3.98E-04 | 5.03E-03 |
| PSMC2 | 0.403 | 5.35E-03 | 3.18E-02 |
| RAB5C | 0.403 | 1.20E-03 | 1.14E-02 |
| LARS | 0.40267 | 7.23E-03 | 3.94E-02 |
| MDN1 | 0.401 | 2.60E-04 | 3.78E-03 |
| VAPB | 0.39867 | 7.89E-03 | 4.16E-02 |
| DARS | 0.398 | 1.39E-03 | 1.25E-02 |
| ATP9A | 0.39667 | 1.15E-03 | 1.11E-02 |
| CALU | 0.39633 | 6.74E-04 | 7.50E-03 |
| RAF1 | 0.39333 | 6.18E-04 | 7.04E-03 |
| NUP107 | 0.39333 | 3.36E-04 | 4.50E-03 |
| COX6C | 0.39267 | 8.74E-04 | 9.18E-03 |
| RSRC1 | 0.39267 | 2.40E-04 | 3.60E-03 |
| P3H3 | 0.39233 | 2.17E-03 | 1.75E-02 |
| TRIP13 | 0.39133 | 1.13E-03 | 1.11E-02 |
| IDH3G | 0.38867 | 2.96E-03 | 2.15E-02 |
| DTWD2 | 0.38733 | 1.49E-03 | 1.34E-02 |
| TMEM165 | 0.38467 | 1.99E-03 | 1.65E-02 |
| SSR4 | 0.38433 | 2.91E-03 | 2.13E-02 |
| FARSA | 0.384 | 8.01E-04 | 8.64E-03 |
| ATP5B | 0.38333 | 1.05E-03 | 1.04E-02 |
| March7 | 0.38267 | 1.81E-03 | 1.55E-02 |
| POM121 | 0.382 | 3.96E-04 | 5.03E-03 |
| VPS52 | 0.38133 | 1.35E-03 | 1.24E-02 |
| TMEM209 | 0.381 | 4.29E-03 | 2.79E-02 |
| CDC42 | 0.37933 | 1.12E-03 | 1.10E-02 |
| NCAPG2 | 0.378 | 3.66E-04 | 4.75E-03 |
| USP24 | 0.37667 | 5.23E-03 | 3.14E-02 |

| | | | |
|---|---|---|---|
| CHST14 | 0.37633 | 1.52E-03 | 1.35E-02 |
| UBE2I | 0.37533 | 3.03E-03 | 2.18E-02 |
| CLASP2 | 0.37433 | 2.57E-03 | 1.95E-02 |
| ARMCX3 | 0.37367 | 5.75E-03 | 3.35E-02 |
| SPTLC1 | 0.37367 | 5.21E-04 | 6.12E-03 |
| NUP98 | 0.37133 | 1.15E-03 | 1.11E-02 |
| CARMIL1 | 0.371 | 8.59E-03 | 4.41E-02 |
| MPP6 | 0.36967 | 2.16E-03 | 1.75E-02 |
| COX4I1 | 0.369 | 1.21E-03 | 1.15E-02 |
| ARFGAP3 | 0.36833 | 4.64E-04 | 5.64E-03 |
| AHCTF1 | 0.36767 | 5.01E-03 | 3.07E-02 |
| BAG2 | 0.36767 | 1.54E-03 | 1.36E-02 |
| TSC2 | 0.365 | 5.81E-03 | 3.37E-02 |
| TUBA1C | 0.36467 | 4.29E-03 | 2.79E-02 |
| TUBA1B | 0.364 | 5.55E-03 | 3.28E-02 |
| DYNC1H1 | 0.36367 | 8.99E-04 | 9.38E-03 |
| GFPT1 | 0.36267 | 8.56E-04 | 9.03E-03 |
| GTF2I | 0.36233 | 3.65E-03 | 2.50E-02 |
| RARS | 0.36167 | 2.21E-03 | 1.76E-02 |
| DYNC1I2 | 0.36 | 7.80E-04 | 8.43E-03 |
| MELK | 0.35867 | 4.19E-04 | 5.24E-03 |
| GEMIN4 | 0.35833 | 3.95E-03 | 2.62E-02 |
| DNAJB11 | 0.357 | 3.87E-03 | 2.60E-02 |
| NIPSNAP1 | 0.35633 | 1.36E-03 | 1.24E-02 |
| ARL1 | 0.35433 | 1.04E-03 | 1.04E-02 |
| MYO19 | 0.354 | 1.37E-03 | 1.25E-02 |
| CORO7 | 0.34833 | 7.17E-03 | 3.91E-02 |
| UQCRH | 0.34667 | 1.99E-03 | 1.66E-02 |
| COG3 | 0.342 | 6.96E-03 | 3.82E-02 |
| RHOT2 | 0.34167 | 5.63E-03 | 3.30E-02 |
| YY1 | 0.34133 | 9.19E-03 | 4.58E-02 |
| LRIG2 | 0.34 | 5.19E-03 | 3.13E-02 |
| PCBP2 | 0.33967 | 3.45E-03 | 2.41E-02 |
| FAF2 | 0.339 | 7.35E-03 | 3.98E-02 |
| NDUFA3 | 0.33767 | 6.69E-03 | 3.72E-02 |
| NR2F6 | 0.33633 | 6.69E-03 | 3.72E-02 |
| DNAJA3 | 0.33533 | 7.47E-04 | 8.14E-03 |
| MTUS1 | 0.335 | 5.20E-03 | 3.13E-02 |
| SLC16A1 | 0.333 | 4.94E-03 | 3.03E-02 |
| GPAT4 | 0.33267 | 6.40E-04 | 7.24E-03 |
| ATP5D | 0.331 | 1.84E-03 | 1.56E-02 |
| KARS | 0.32767 | 7.96E-03 | 4.19E-02 |
| NDUFS7 | 0.32767 | 2.97E-03 | 2.15E-02 |
| ULK3 | 0.327 | 6.34E-03 | 3.62E-02 |
| ACAD9 | 0.327 | 4.58E-03 | 2.87E-02 |
| RNF213 | 0.326 | 6.66E-03 | 3.72E-02 |
| CNOT11 | 0.32567 | 4.17E-03 | 2.72E-02 |
| SEC22B | 0.324 | 1.30E-03 | 1.21E-02 |
| RAB11FIP2 | 0.32367 | 9.62E-03 | 4.70E-02 |

| | | | |
|---|---|---|---|
| COG1 | 0.322 | 3.22E-03 | 2.28E-02 |
| RHOT1 | 0.32167 | 2.02E-03 | 1.67E-02 |
| HELLS | 0.31967 | 6.50E-03 | 3.68E-02 |
| NDUFB10 | 0.31833 | 7.06E-03 | 3.86E-02 |
| LRWD1 | 0.318 | 4.41E-03 | 2.82E-02 |
| CNOT10 | 0.316 | 5.18E-03 | 3.13E-02 |
| DGKA | 0.31567 | 5.02E-04 | 5.97E-03 |
| AAR2 | 0.31433 | 4.20E-03 | 2.73E-02 |
| UQCRC2 | 0.314 | 4.39E-03 | 2.82E-02 |
| SUCLA2 | 0.31367 | 2.08E-03 | 1.70E-02 |
| MTHFD1 | 0.31133 | 9.69E-03 | 4.71E-02 |
| POM121C | 0.30867 | 5.93E-03 | 3.42E-02 |
| RPTOR | 0.30867 | 4.18E-03 | 2.73E-02 |
| ARHGEF40 | 0.30833 | 5.80E-03 | 3.37E-02 |
| JAK1 | 0.30533 | 9.08E-03 | 4.56E-02 |
| NDUFS1 | 0.30167 | 5.83E-03 | 3.38E-02 |
| CCT7 | 0.30033 | 7.02E-03 | 3.84E-02 |
| TIMM50 | 0.299 | 3.85E-03 | 2.59E-02 |
| CU | 0.29867 | 7.37E-03 | 3.98E-02 |
| CDK17 | 0.29867 | 2.00E-03 | 1.66E-02 |
| DPM1 | 0.298 | 5.03E-03 | 3.07E-02 |
| UBE2O | 0.29633 | 8.38E-03 | 4.34E-02 |
| PPM1F | 0.29467 | 4.31E-03 | 2.79E-02 |
| SENP1 | 0.29267 | 6.96E-03 | 3.82E-02 |
| IGF2R | 0.292 | 1.61E-03 | 1.41E-02 |
| TARDBP | 0.291 | 5.42E-03 | 3.22E-02 |
| CLASP1 | 0.29033 | 5.06E-03 | 3.08E-02 |
| RANBP2 | 0.29033 | 4.15E-03 | 2.71E-02 |
| PIK3R2 | 0.28533 | 3.08E-03 | 2.20E-02 |
| PIK3C3 | 0.283 | 6.76E-03 | 3.75E-02 |
| IRAK1 | 0.28167 | 7.84E-03 | 4.14E-02 |
| DHRS7B | 0.28067 | 6.91E-03 | 3.82E-02 |
| UBIAD1 | 0.28067 | 3.51E-03 | 2.43E-02 |
| DNM2 | 0.27833 | 8.85E-03 | 4.49E-02 |
| RBM4 | 0.27667 | 6.66E-03 | 3.72E-02 |
| AGPS | 0.276 | 5.41E-03 | 3.21E-02 |
| DSP | 0.27567 | 9.26E-03 | 4.60E-02 |
| NDUFB4 | 0.275 | 6.36E-03 | 3.62E-02 |
| KNSTRN | 0.274 | 3.72E-03 | 2.53E-02 |
| ACSL3 | 0.271 | 3.10E-03 | 2.21E-02 |
| USP9X | 0.27067 | 4.66E-03 | 2.90E-02 |
| RER1 | 0.27 | 6.05E-03 | 3.47E-02 |
| AAAS | 0.26633 | 3.10E-03 | 2.21E-02 |
| CYC1 | 0.26233 | 4.84E-03 | 2.99E-02 |
| PPP6R3 | 0.262 | 1.38E-03 | 1.25E-02 |
| HCK | 0.26133 | 4.02E-03 | 2.65E-02 |
| CKAP4 | 0.25967 | 5.79E-03 | 3.37E-02 |
| PNMA8A | 0.25867 | 7.51E-03 | 4.02E-02 |
| DROSHA | 0.25833 | 8.97E-03 | 4.53E-02 |

| | | | |
|---|---|---|---|
| LPCAT4 | 0.25633 | 3.92E-03 | 2.61E-02 |
| CABIN1 | 0.25233 | 6.59E-03 | 3.72E-02 |
| RPS27 | 0.252 | 9.10E-03 | 4.57E-02 |
| CDK20 | 0.25133 | 5.23E-03 | 3.14E-02 |
| CCT6A | 0.251 | 2.48E-03 | 1.90E-02 |
| POLG | 0.24967 | 8.35E-03 | 4.34E-02 |
| ARF5 | 0.24833 | 8.76E-03 | 4.45E-02 |
| NDC1 | 0.247 | 8.19E-03 | 4.28E-02 |
| RBM26 | 0.24667 | 4.04E-03 | 2.66E-02 |
| HSP90AA1 | 0.24433 | 9.63E-03 | 4.70E-02 |
| NSDHL | 0.24433 | 3.08E-03 | 2.20E-02 |
| RNASEL | 0.24033 | 1.02E-02 | 4.92E-02 |
| SLC27A4 | 0.23967 | 8.41E-03 | 4.35E-02 |
| REXO2 | 0.23833 | 5.20E-03 | 3.13E-02 |
| PE | 0.21933 | 6.10E-03 | 3.49E-02 |
| RBM27 | 0.208 | 6.98E-03 | 3.82E-02 |
| SOX9 | 0.198 | 9.58E-03 | 4.70E-02 |

**Table S2. Arrayed CRISPR/Cas9 knockout screen of MYC-TMED9 interactors in P cells by mean normalized MUC1-fs intensity.**

Mean normalized MUC1-fs intensity per target interactor gRNA is shown. MUC1-fs intensity normalized to mean MUC1 and non-targeting control (NTC) gRNA. Adjusted p-value maximum 4.0.

| Target | gRNA clone (GPP) | Mean normalized MUC1-fs intensity | -log10(adj. p-value) |
|---|---|---|---|
| MUC1 | BRDN0001065880 | -99.73 | 4.00E+00 |
| CSE1L | BRDN0004620574 | -64.89 | 4.00E+00 |
| TMED9 | BRDN0003481199 | -61.66 | 4.00E+00 |
| ATP1A1 | BRDN0001489778 | -57.31 | 4.00E+00 |
| TMED10 | BRDN0003581071 | -51.23 | 3.28E+00 |
| TMED7 | BRDN0003481231 | -48.14 | 2.70E+00 |
| ATP1A1 | BRDN0001488278 | -46.59 | 2.41E+00 |
| COPB2 | BRDN0004308235 | -40.79 | 1.44E+00 |
| XPO5 | BRDN0003789292 | -27.96 | 6.06E-02 |
| RPN1 | BRDN0002662586 | -25.89 | 1.30E-02 |
| ATP1A1 | BRDN0001488366 | -21.62 | 9.59E-04 |
| SLC30A7 | BRDN0001504553 | -21.29 | 9.32E-04 |
| CLCN5 | BRDN0003233263 | -21.12 | 9.06E-04 |
| CSE1L | BRDN0004620733 | -19.43 | 7.73E-04 |
| TMED10 | BRDN0003230535 | -17.21 | 6.14E-04 |
| CAND1 | BRDN0003789268 | -16.39 | 5.61E-04 |
| XPO1 | BRDN0001053803 | -16.37 | 5.61E-04 |
| CSE1L | BRDN0005318341 | -16.12 | 5.34E-04 |
| NPEPPS | BRDN0001485127 | -13.27 | 4.01E-04 |
| XPO5 | BRDN0003789291 | -12.89 | 3.75E-04 |
| SELENOI | BRDN0001501953 | -12.52 | 3.75E-04 |
| RNF185 | BRDN0003675322 | -12.14 | 3.48E-04 |
| ERGIC2 | BRDN0003491628 | -11.73 | 1.36E-04 |
| FBXO28 | BRDN0003418810 | -10.14 | 2.69E-04 |
| COPB2 | BRDN0004310474 | -10.06 | 2.69E-04 |
| UBE4A | BRDN0004618624 | -8.51 | 2.16E-04 |
| DDOST | BRDN0002661282 | -8 | 2.16E-04 |
| SELENOI | BRDN0001501610 | -5.69 | 1.36E-04 |
| EXOC6 | BRDN0003482793 | -5.37 | 1.36E-04 |
| RNF5 | BRDN0003487834 | -4.68 | 1.10E-04 |
| HOOK3 | BRDN0003512502 | -4.47 | 1.10E-04 |
| VIM | BRDN0005315643 | -3.67 | 8.32E-05 |
| VKORC1L1 | BRDN0001483648 | -3.67 | 8.32E-05 |
| LMAN1 | BRDN0003492359 | -3.55 | 8.32E-05 |
| YTHDF3 | BRDN0003493281 | -3.33 | 8.32E-05 |
| UBR3 | BRDN0003233536 | -2.89 | 5.67E-05 |

| | | | |
|---|---|---|---|
| SLC30A9 | BRDN0004309772 | -2.75 | 5.67E-05 |
| ATF6 | BRDN0001484163 | -1.84 | 4.34E-05 |
| ATXN10 | BRDN0002661502 | -1.24 | 4.34E-05 |
| SET | BRDN0000733796 | -1.22 | 4.34E-05 |
| ARCN1 | BRDN0005316639 | -0.84 | 4.34E-05 |
| VDAC2 | BRDN0000583713 | -0.24 | 4.34E-05 |
| ATP1B3 | BRDN0001484128 | -0.23 | 4.34E-05 |
| RINT1 | BRDN0003488288 | 2.5 | 5.67E-05 |
| NPEPPS | BRDN0001488390 | 3.34 | 8.32E-05 |
| FBXO28 | BRDN0003492069 | 4.55 | 1.10E-04 |
| SPG11 | BRDN0003675609 | 5.81 | 1.36E-04 |
| SAAL1 | BRDN0003229359 | 6.54 | 1.63E-04 |
| ARCN1 | BRDN0004309486 | 6.64 | 1.63E-04 |
| HACD3 | BRDN0003491840 | 7.04 | 1.89E-04 |
| ERGIC2 | BRDN0003482450 | 7.67 | 1.89E-04 |
| GLMN | BRDN0003229841 | 7.92 | 2.16E-04 |
| RABL3 | BRDN0003440034 | 8.01 | 2.16E-04 |
| STT3A | BRDN0002661226 | 8.02 | 2.16E-04 |
| ATF6 | BRDN0001487465 | 8.41 | 2.16E-04 |
| PPIB | BRDN0001483815 | 9.23 | 2.42E-04 |
| STT3A | BRDN0002662166 | 9.98 | 2.69E-04 |
| ZDHHC13 | BRDN0003481518 | 9.98 | 2.69E-04 |
| YIF1B | BRDN0003514201 | 10.28 | 2.95E-04 |
| ZMPSTE24 | BRDN0003099273 | 11.19 | 3.22E-04 |
| SET | BRDN0000733365 | 11.76 | 3.48E-04 |
| TMED9 | BRDN0003481863 | 11.85 | 3.48E-04 |
| RNF185 | BRDN0003577634 | 12.33 | 3.75E-04 |
| PPIB | BRDN0001482792 | 12.4 | 3.75E-04 |
| CAND1 | BRDN0003789538 | 12.42 | 3.75E-04 |
| VKORC1L1 | BRDN0001488893 | 12.71 | 3.75E-04 |
| HOOK2 | BRDN0004308366 | 12.8 | 3.75E-04 |
| EXOC6 | BRDN0003579623 | 13.06 | 4.01E-04 |
| MANEA | BRDN0003786468 | 13.39 | 4.01E-04 |
| TMED7 | BRDN0003482908 | 13.54 | 4.28E-04 |
| ARL6IP6 | BRDN0003462870 | 14.36 | 4.54E-04 |
| CLCN5 | BRDN0003233375 | 14.46 | 4.54E-04 |
| VKORC1L1 | BRDN0001479757 | 14.88 | 4.81E-04 |
| HOOK3 | BRDN0003676604 | 15.15 | 5.08E-04 |
| MKS1 | BRDN0003581731 | 15.15 | 5.08E-04 |
| VDAC2 | BRDN0000583494 | 15.2 | 5.08E-04 |
| XPO1 | BRDN0003482103 | 16.05 | 5.34E-04 |
| RLIM | BRDN0003789316 | 16.23 | 5.61E-04 |
| EI24 | BRDN0003578771 | 17 | 5.87E-04 |
| DYM | BRDN0003481135 | 18.38 | 6.93E-04 |
| SLC30A7 | BRDN0001496874 | 18.75 | 7.20E-04 |
| KIAA1524 | BRDN0001065354 | 19.04 | 7.46E-04 |
| SELENOI | BRDN0001503933 | 19.39 | 7.73E-04 |
| UBE3B | BRDN0003512819 | 19.5 | 7.73E-04 |
| RNF5 | BRDN0003677146 | 20.31 | 8.53E-04 |

| | | | |
|---|---|---|---|
| DYM | BRDN0003581707 | 20.93 | 9.06E-04 |
| LENG1 | BRDN0003490376 | 21.31 | 9.32E-04 |
| STARD3 | BRDN0001501816 | 21.81 | 3.60E-03 |
| NGLY1 | BRDN0004437338 | 21.95 | 3.68E-03 |
| PRKDC | BRDN0001149021 | 21.98 | 3.68E-03 |
| RLIM | BRDN0003581834 | 22.39 | 3.89E-03 |
| SLC7A1 | BRDN0001482549 | 22.77 | 4.10E-03 |
| RCN1 | BRDN0003786294 | 22.86 | 4.16E-03 |
| VIM | BRDN0005318202 | 23.24 | 4.35E-03 |
| HACD3 | BRDN0003481785 | 23.33 | 4.40E-03 |
| EXOC5 | BRDN0003229230 | 23.36 | 4.43E-03 |
| TMCO1 | BRDN0003491792 | 23.36 | 4.43E-03 |
| KIAA1524 | BRDN0001491034 | 23.9 | 7.97E-03 |
| CYBA | BRDN0001483865 | 24.22 | 8.56E-03 |
| COG6 | BRDN0003676872 | 24.79 | 9.57E-03 |
| ZDHHC13 | BRDN0003481212 | 25.18 | 3.65E-03 |
| TBC1D8B | BRDN0003483033 | 25.39 | 1.09E-02 |
| MANEA | BRDN0003489569 | 25.6 | 1.16E-02 |
| ATP1B3 | BRDN0001484585 | 26.55 | 2.48E-02 |
| NGLY1 | BRDN0004437152 | 26.62 | 2.63E-02 |
| RABL3 | BRDN0003439449 | 26.76 | 2.93E-02 |
| XPO4 | BRDN0004437906 | 27.2 | 3.96E-02 |
| HACD3 | BRDN0005318390 | 27.46 | 4.66E-02 |
| XPOT | BRDN0000583493 | 27.53 | 4.87E-02 |
| JAK3 | BRDN0001065886 | 27.72 | 5.47E-02 |
| DDOST | BRDN0002662287 | 27.74 | 5.53E-02 |
| GLMN | BRDN0003228099 | 28 | 6.43E-02 |
| ERLEC1 | BRDN0003441061 | 28.74 | 9.44E-02 |
| XPO4 | BRDN0004437208 | 29.03 | 1.08E-01 |
| ERGIC3 | BRDN0003482036 | 29.66 | 1.43E-01 |
| VDAC2 | BRDN0000583490 | 29.98 | 1.63E-01 |
| PRKDC | BRDN0001145772 | 30.04 | 1.67E-01 |
| ATXN10 | BRDN0002661454 | 30.38 | 1.89E-01 |
| PRKDC | BRDN0001149438 | 30.81 | 2.20E-01 |
| VAPA | BRDN0003577536 | 30.95 | 2.31E-01 |
| RPN1 | BRDN0002661706 | 31.25 | 2.54E-01 |
| XPOT | BRDN0000583586 | 31.43 | 2.69E-01 |
| ASS1 | BRDN0001482897 | 31.46 | 2.72E-01 |
| CUX1 | BRDN0004308398 | 31.52 | 2.77E-01 |
| RPN1 | BRDN0002662056 | 31.62 | 2.85E-01 |
| EI24 | BRDN0003579090 | 31.75 | 2.96E-01 |
| GLMN | BRDN0003228143 | 31.88 | 3.09E-01 |
| KIAA1524 | BRDN0001065265 | 32.28 | 3.45E-01 |
| ZMPSTE24 | BRDN0003462398 | 32.33 | 3.49E-01 |
| SLC7A1 | BRDN0001489586 | 32.49 | 3.65E-01 |
| JAK3 | BRDN0001065702 | 32.81 | 3.96E-01 |
| MKS1 | BRDN0003481567 | 32.9 | 4.05E-01 |
| UBE4A | BRDN0004308101 | 33.21 | 4.37E-01 |
| YTHDF3 | BRDN0003419243 | 33.83 | 5.04E-01 |

| | | | |
|---|---|---|---|
| STT3A | BRDN0002661250 | 34.09 | 5.32E-01 |
| TMED4 | BRDN0003292311 | 34.65 | 5.98E-01 |
| MZT2A | BRDN0003486425 | 35.07 | 6.48E-01 |
| PKDREJ | BRDN0003098737 | 36.05 | 7.70E-01 |
| RINT1 | BRDN0003674647 | 36.06 | 7.72E-01 |
| SET | BRDN0000733691 | 36.21 | 7.91E-01 |
| SAAL1 | BRDN0003230674 | 36.46 | 8.24E-01 |
| XPO6 | BRDN0003227864 | 36.54 | 8.35E-01 |
| TBC1D8B | BRDN0003581050 | 36.88 | 8.81E-01 |
| BTAF1 | BRDN0004309909 | 37.19 | 9.23E-01 |
| VAPA | BRDN0003578188 | 37.98 | 1.03E+00 |
| CYBA | BRDN0001481752 | 38.62 | 1.12E+00 |
| ERGIC3 | BRDN0003492871 | 38.9 | 1.17E+00 |
| PKDREJ | BRDN0003098757 | 39.34 | 1.23E+00 |
| STARD3 | BRDN0001502427 | 39.5 | 1.25E+00 |
| UBR3 | BRDN0003234374 | 39.98 | 1.33E+00 |
| ARL6IP6 | BRDN0003233203 | 40.33 | 1.38E+00 |
| ANO10 | BRDN0003791621 | 40.8 | 1.45E+00 |
| HEATR3 | BRDN0003692951 | 41.12 | 1.51E+00 |
| FAM114A2 | BRDN0003228996 | 41.26 | 1.53E+00 |
| TNPO2 | BRDN0003789585 | 41.28 | 1.53E+00 |
| VPS50 | BRDN0003675517 | 42.5 | 1.73E+00 |
| XPO6 | BRDN0003228050 | 42.91 | 1.79E+00 |
| MZT2B | BRDN0003675755 | 44.36 | 2.04E+00 |
| STARD3 | BRDN0001500209 | 45.07 | 2.16E+00 |
| ASS1 | BRDN0001481696 | 45.68 | 2.26E+00 |
| UBE3B | BRDN0003513235 | 46.03 | 2.32E+00 |
| XPO1 | BRDN0001053798 | 46.18 | 2.35E+00 |
| SC5D | BRDN0003675041 | 47.32 | 2.56E+00 |
| CYBA | BRDN0001485647 | 47.33 | 2.56E+00 |
| VIM | BRDN0004618501 | 47.46 | 2.58E+00 |
| YIF1B | BRDN0003511856 | 47.56 | 2.60E+00 |
| NCAPH | BRDN0001496173 | 48.69 | 2.81E+00 |
| RCN1 | BRDN0003487870 | 48.82 | 2.83E+00 |
| CUX1 | BRDN0005315860 | 50.11 | 3.08E+00 |
| SLC7A1 | BRDN0001480840 | 50.31 | 3.11E+00 |
| HEATR3 | BRDN0003482193 | 50.39 | 3.13E+00 |
| TUBGCP2 | BRDN0004619161 | 52.15 | 3.47E+00 |
| LENG1 | BRDN0003488622 | 53.71 | 3.79E+00 |
| SPG11 | BRDN0003786553 | 54.9 | 4.00E+00 |
| TMEM161A | BRDN0003483166 | 55.08 | 4.00E+00 |
| EXOC7 | BRDN0003418047 | 55.23 | 4.00E+00 |
| DDOST | BRDN0002662308 | 55.55 | 4.00E+00 |
| RNF185 | BRDN0003790052 | 56.14 | 4.00E+00 |
| LMAN1 | BRDN0003579603 | 58.1 | 4.00E+00 |
| RPL38 | BRDN0004308732 | 58.24 | 4.00E+00 |
| INTS12 | BRDN0004435252 | 58.74 | 4.00E+00 |
| TNPO1 | BRDN0001053805 | 59.99 | 1.92E-01 |
| XPOT | BRDN0004620183 | 61.03 | 4.00E+00 |

| | | | |
|---|---|---|---|
| JAK3 | BRDN0001065699 | 61.11 | 4.00E+00 |
| Exoc2 | BRDN0003827348 | 61.8 | 4.00E+00 |
| TTYH3 | BRDN0003676678 | 62.76 | 4.00E+00 |
| ATXN10 | BRDN0002661891 | 65.89 | 4.00E+00 |
| EXOC7 | BRDN0003417181 | 66.31 | 4.00E+00 |
| EXOC8 | BRDN0004435602 | 66.61 | 4.00E+00 |
| IPO9 | BRDN0003485505 | 67.97 | 4.00E+00 |
| MZT2B | BRDN0003786680 | 71.44 | 4.00E+00 |
| INTS12 | BRDN0004620016 | 72.19 | 4.00E+00 |
| EXOC5 | BRDN0003229792 | 72.54 | 4.00E+00 |
| SLC30A7 | BRDN0001496936 | 72.62 | 4.00E+00 |
| ERLEC1 | BRDN0003439516 | 76.49 | 4.00E+00 |
| ANO10 | BRDN0003485205 | 79.11 | 4.00E+00 |
| HOOK2 | BRDN0003438935 | 79.23 | 4.00E+00 |
| NCAPH | BRDN0001497174 | 79.49 | 4.00E+00 |
| LOC101929950 | BRDN0004437028 | 80.1 | 4.00E+00 |
| EXOC2 | BRDN0003484202 | 80.28 | 4.00E+00 |
| VPS50 | BRDN0003485240 | 81.27 | 4.00E+00 |
| ASS1 | BRDN0001485125 | 81.76 | 4.00E+00 |
| PPIB | BRDN0001480019 | 82 | 4.00E+00 |
| IPO9 | BRDN0003485217 | 82.74 | 4.00E+00 |
| SC5D | BRDN0003676178 | 82.93 | 4.00E+00 |
| TGIF2-C20orf24 | BRDN0003227710 | 85.7 | 4.00E+00 |
| TUBGCP2 | BRDN0004309201 | 88.47 | 4.00E+00 |
| FAM114A2 | BRDN0003230048 | 89.41 | 4.00E+00 |
| EXOC2 | BRDN0003485674 | 90.83 | 4.00E+00 |
| NCAPD2 | BRDN0004309346 | 94.86 | 4.00E+00 |
| TGIF2-C20orf24 | BRDN0004437554 | 101.06 | 4.00E+00 |
| RPL38 | BRDN0004310303 | 103.2 | 4.00E+00 |
| C19orf25 | BRDN0003439510 | 103.97 | 4.00E+00 |
| TMEM161A | BRDN0003481163 | 105.1 | 4.00E+00 |
| PPP4R3B | BRDN0004437764 | 106.52 | 4.00E+00 |
| C20orf24 | BRDN0003485359 | 115.01 | 4.00E+00 |
| TNPO3 | BRDN0003228034 | 130.57 | 4.00E+00 |
| TNPO3 | BRDN0003227889 | 154 | 4.00E+00 |
| SURF4 | BRDN0003486326 | 160.69 | 4.00E+00 |
| ATP2A2 | BRDN0001482972 | 164.54 | 4.00E+00 |
| ATP2A2 | BRDN0001482206 | 177.14 | 4.00E+00 |
| NCAPD2 | BRDN0004310515 | 178.53 | 4.00E+00 |
| NCAPH | BRDN0001497233 | 189.16 | 4.00E+00 |
| SURF4 | BRDN0003486926 | 414.46 | 4.00E+00 |
| ATP2A2 | BRDN0001483123 | 423.9 | 4.00E+00 |

**Table S3: In-vitro pharmacology study for BRD7635 at 10 mM.**

% Inhibition >50% of control was considered significant.

| **Compound** | **% Inhibition of Control Specific Binding** |
|---|---|
| 5-HT transporter (h) (antagonist radioligand) | Non-significant |
| *5-HT1A(h) (agonist radioligand)* | Non-significant |
| *5-HT1B (h) (antagonist radioligand)* | Non-significant |
| *5-HT2A(h) (agonist radioligand)* | Non-significant |
| *5-HT2B(h) (agonist radioligand)* | Non-significant |
| *5-HT3(h) (antagonist radioligand)* | Non-significant |
| *alpha1A(h) (antagonist radioligand)* | Non-significant |
| *alpha2A(h) (antagonist radioligand)* | Non-significant |
| *A2A(h) (agonist radioligand)* | Non-significant |
| *AR(h) (agonist radioligand)* | Non-significant |
| *beta1(h) (agonist radioligand)* | Non-significant |
| *beta2(h) (antagonist radioligand)* | Non-significant |
| *BZD (central)(h) (agonist radioligand)* | Non-significant |
| *KV channel (antagonist radioligand)* | Non-significant |
| *Cav1.2 (L-type) Human Calcium Ion Channel Binding (Dihydropyridine Site) LeadHunter Assay - FR* | Non-significant |
| *CB2(h) (agonist radioligand)* | Non-significant |
| *CB1(h) (agonist radioligand)* | Non-significant |
| *CCK1 (CCKA) (h) (agonist radioligand)* | Non-significant |
| *D1(h) (antagonist radioligand)* | Non-significant |
| *D2S(h) (agonist radioligand)* | Non-significant |
| *delta (DOP) (h) (agonist radioligand)* | Non-significant |
| *dopamine transporter(h) (antagonist radioligand)* | Non-significant |
| *ETA(h) (agonist radioligand)* | Non-significant |
| *Glutamate (NMDA NR1/NR2A) Human Ion Channel [3H] CGP-39653 Binding* | Non-significant |
| *GR (h) (agonist radioligand)* | Non-significant |
| *H1(h) (antagonist radioligand)* | Non-significant |
| *H2(h) (antagonist radioligand)* | Non-significant |
| *kappa (h) (KOP) (agonist radioligand)* | Non-significant |
| *M1(h) (antagonist radioligand)* | Non-significant |
| *M2 (h) (antagonist radioligand)* | Non-significant |
| *M3(h) (antagonist radioligand)* | Non-significant |
| *MAO-A (antagonist radioligand)* | Non-significant |
| *μ (MOP) (h) (agonist radioligand)* | Non-significant |

| | |
|---|---|
| *N neuronal alpha4beta2 (h) (agonist radioligand)* | Non-significant |
| *norepinephrine transporter(h) (antagonist radioligand)* | Non-significant |
| *Potassium Channel hERG (human)- [3H] Dofetilide* | Non-significant |
| *Sodium Channel Site2 (Non-selective) Rat Ion Channel Batrachotoxin Mass Spectrometry Binding* | Non-significant |
| *V1a(h) (agonist radioligand)* | Non-significant |
| *acetylcholinesterase (h)* | Non-significant |
| *PDE3A (h)* | Non-significant |
| *PDE4D2 (h)* | Non-significant |
| *COX1(h)* | Non-significant |
| *COX2(h)* | Non-significant |
| *Lck Human TK Kinase Enzymatic Radiometric Assay [Km ATP]* | Non-significant |

**Table S4. IP-MS of TMED5-MYC and TMED7-MYC Co-IP.**

Proteins (with gene names) that preferably bind TMED5 (negative logFC value) or TMED7 (positive logFC value). Proteins are sorted with logFC value and were analyzed using the Limma package in R.

### TMED5 preferred

| Gene | logFC | Average expression | Adj. p value |
|---|---|---|---|
| TMED5 | -6.589298 | 25.2499502 | 7.76E-12 |
| UQCRB | -5.9129959 | 18.9814001 | 1.89E-13 |
| PET100 | -5.1439785 | 18.7250609 | 4.91E-11 |
| MPDU1 | -5.1269568 | 18.719387 | 1.68E-13 |
| COA1 | -4.8918114 | 16.7913166 | 3.08E-08 |
| ATAD3B | -4.6930918 | 23.1765167 | 1.30E-07 |
| NDUFA11 | -4.3999072 | 19.7747412 | 1.82E-09 |
| RAB1B | -4.3733848 | 20.0498528 | 3.70E-12 |
| LSM4 | -3.8870387 | 19.4626641 | 1.76E-07 |
| MOSPD2 | -3.8769765 | 19.9548157 | 1.07E-11 |
| NAP1L1;NAP1L4 | -3.6874492 | 19.1748986 | 4.48E-11 |
| TVP23B | -3.6367894 | 18.2226646 | 6.01E-12 |
| TMEM14C | -3.4626828 | 18.164629 | 1.11E-12 |
| ATAD3A | -3.3112969 | 24.4413885 | 4.54E-12 |
| NDUFS4 | -2.9341676 | 18.9056801 | 1.02E-11 |
| AURKA | -2.8150021 | 18.467099 | 1.32E-11 |
| DHX29 | -2.7890245 | 15.3030143 | 2.60E-09 |

| | | | |
|---|---|---|---|
| PEX6 | -2.7727003 | 17.9346349 | 1.05E-11 |
| LONP2 | -2.7567724 | 18.1174756 | 1.49E-10 |
| SLC25A46 | -2.7398815 | 17.9236952 | 8.42E-12 |
| FRG1 | -2.7308269 | 18.5400336 | 4.50E-10 |
| ADGRB3 | -2.6964314 | 16.0642267 | 8.36E-11 |
| MCAT | -2.6444485 | 16.0815544 | 6.93E-11 |
| INA | -2.6052762 | 19.5912961 | 6.20E-04 |
| ZBED1 | -2.5583081 | 18.0182785 | 6.70E-05 |
| HVCN1 | -2.5259859 | 17.8273871 | 1.40E-08 |
| ASPM | -2.5062038 | 17.8458027 | 9.99E-12 |
| ODR4 | -2.4544214 | 17.8285419 | 9.99E-12 |
| STX17 | -2.4456288 | 18.8648203 | 1.37E-06 |
| IGHV1-3;IGHV1-46;IGHV1-69;IGHV1-69D;IGHV1-8 | -2.4292294 | 17.8201446 | 1.18E-11 |
| NDUFS3 | -2.3934893 | 19.7706461 | 1.25E-07 |
| NDST2 | -2.3850666 | 16.168015 | 2.15E-07 |
| SMO | -2.3780275 | 16.1703614 | 1.27E-08 |
| SLC25A16 | -2.3504856 | 18.8810419 | 1.01E-10 |
| SFXN2 | -2.3033567 | 20.8994095 | 1.12E-05 |
| APOA1 | -2.2939439 | 18.7753881 | 3.94E-06 |
| ZNHIT6 | -2.2858367 | 17.772347 | 4.99E-11 |
| SERPINA12 | -2.2621666 | 17.7644569 | 2.41E-09 |
| AGPAT5 | -2.2419423 | 17.2609777 | 4.33E-10 |

| | | | |
|---|---|---|---|
| TCTN1 | -2.2298746 | 16.2197457 | 5.07E-05 |
| VPS13B | -2.2071094 | 17.7461045 | 2.29E-07 |
| GABPA | -2.181266 | 17.4176118 | 1.11E-04 |
| THUMPD3 | -2.1642976 | 17.7318339 | 6.17E-10 |
| MFN2 | -2.1465773 | 20.1023364 | 0.10026966 |
| SLC25A11 | -2.1242666 | 25.0088395 | 2.19E-09 |
| LYZ | -2.112728 | 18.7616991 | 1.42E-09 |
| PTPRG | -2.1059154 | 16.2610654 | 3.95E-10 |
| DDX28 | -2.1042839 | 18.068612 | 2.83E-08 |
| MTCH1 | -2.053462 | 22.5800876 | 4.44E-07 |
| ZNF568 | -2.0367761 | 16.3267136 | 2.17E-09 |
| CHRNA3 | -2.0318215 | 16.2857634 | 4.58E-09 |
| CCDC8 | -2.0250214 | 17.793864 | 2.27E-07 |
| ATP9A | -2.0126326 | 19.0411319 | 4.88E-07 |
| PDAP1 | -2.010518 | 18.113003 | 3.80E-11 |
| GDF11 | -2.0027525 | 16.295453 | 6.46E-10 |
| DCD | -2.0016171 | 22.0668457 | 0.00689375 |
| WDR35 | -2.0008005 | 16.2961037 | 1.97E-08 |
| IGHG1 | -1.9945766 | 19.4959446 | 1.74E-04 |
| NDUFAF4 | -1.9923611 | 18.8963282 | 3.25E-09 |
| ANTXR2 | -1.9649461 | 16.3080552 | 1.00E-08 |
| SLC2A1 | -1.9541388 | 17.1831653 | 3.32E-08 |
| PLCH1 | -1.948956 | 16.3133852 | 1.69E-09 |

| | | | |
|---|---|---|---|
| SLC25A12 | -1.9392796 | 21.7490006 | 3.73E-10 |
| YME1L1 | -1.9294537 | 21.5686149 | 1.15E-07 |
| ISOC1 | -1.9098749 | 21.213353 | 8.34E-05 |
| ATP5MK | -1.9093903 | 22.8004407 | 5.85E-07 |
| ABCB7 | -1.8924674 | 18.854389 | 9.37E-09 |
| SFXN3 | -1.8705434 | 19.6041804 | 4.44E-07 |
| MCM7 | -1.8703043 | 21.4972891 | 6.87E-08 |
| SCD5 | -1.8688637 | 18.0706954 | 3.15E-04 |
| NDUFV2 | -1.8634361 | 18.9996972 | 3.06E-07 |
| SLC25A30 | -1.8479546 | 17.9262899 | 1.88E-07 |
| PLXDC2 | -1.8428967 | 17.7577554 | 1.89E-08 |
| TTC28 | -1.8340744 | 16.629757 | 2.61E-07 |
| UCP2 | -1.8314349 | 16.2186849 | 3.81E-07 |
| SFXN4 | -1.8256337 | 21.6228363 | 1.44E-07 |
| ADGRL3 | -1.8196867 | 16.356475 | 5.09E-07 |
| FGA | -1.7963585 | 16.2353016 | 3.13E-10 |
| SLC27A4 | -1.7950568 | 21.3307474 | 4.38E-10 |
| SLC43A2 | -1.7777695 | 16.1605935 | 7.01E-08 |
| NDUFS6 | -1.7677245 | 18.1419208 | 2.82E-09 |
| GOLGB1 | -1.7572353 | 18.7692001 | 2.47E-10 |
| KLHL26 | -1.7504507 | 16.3795537 | 1.12E-07 |
| C1QTNF3 | -1.7449903 | 16.6106887 | 2.04E-08 |
| IDH1 | -1.7414639 | 17.5908894 | 6.51E-10 |

| | | | |
|---|---|---|---|
| RPS6KB1 | -1.7379525 | 17.5897189 | 1.12E-09 |
| CALM1;CALM2;CALM3 | -1.7262144 | 17.5858062 | 7.75E-10 |
| SYDE1 | -1.7261519 | 17.4051236 | 1.03E-08 |
| TRIOBP | -1.7183516 | 20.5465841 | 4.82E-10 |
| NDUFS1 | -1.7058308 | 19.0742929 | 4.43E-08 |
| MOCS3 | -1.6991573 | 17.5767872 | 4.82E-10 |
| APOOL | -1.6970136 | 20.0579926 | 2.31E-08 |
| METTL9 | -1.6880621 | 16.4003499 | 7.52E-07 |
| ZMYM4 | -1.6862011 | 18.1851109 | 3.14E-05 |
| COMTD1 | -1.6796162 | 18.8060239 | 2.90E-09 |
| KLHL18 | -1.6722027 | 16.7937733 | 9.69E-08 |
| NDUFA7 | -1.6711135 | 16.8459171 | 3.81E-08 |
| ANO8 | -1.6686594 | 16.4068174 | 7.52E-06 |
| MAOA | -1.6679506 | 17.5330588 | 6.36E-08 |
| NDUFA5 | -1.6573567 | 20.8975243 | 6.32E-08 |
| GTF3C5 | -1.6565802 | 16.8318452 | 5.09E-08 |
| NXT1;NXT2 | -1.6440063 | 17.5584035 | 3.23E-07 |
| SCO2 | -1.6309798 | 20.9233172 | 7.44E-08 |
| CANT1 | -1.6251536 | 17.6105793 | 2.82E-08 |
| DNAJA2 | -1.6247466 | 18.8393374 | 8.78E-05 |
| PIK3R2 | -1.6164782 | 17.1999896 | 1.65E-05 |
| PJA2 | -1.6037183 | 18.2171669 | 1.21E-06 |
| SLC25A13 | -1.5905302 | 23.5079305 | 1.17E-09 |

| | | | |
|---|---|---|---|
| ABCB10 | -1.5823309 | 18.8037184 | 2.43E-09 |
| UTP25 | -1.5817062 | 15.5094062 | 6.42E-05 |
| LPCAT4 | -1.5765771 | 17.4555245 | 7.78E-09 |
| DIRAS1 | -1.5732772 | 17.5923723 | 9.43E-09 |
| PHKG2 | -1.57192 | 20.0939565 | 3.03E-09 |
| WDR45 | -1.5669719 | 17.9182639 | 0.00112704 |
| PHKA2 | -1.5668228 | 21.0193079 | 0.00449218 |
| ZNF146 | -1.5664056 | 17.9741125 | 4.31E-09 |
| TLCD3A | -1.5636462 | 17.6280267 | 1.87E-07 |
| RUVBL2 | -1.5591772 | 21.059079 | 6.36E-05 |
| C3ORF33 | -1.5587397 | 18.9894946 | 3.74E-04 |
| ABCD1 | -1.5494193 | 22.4042307 | 2.42E-05 |
| ZDHHC9 | -1.5437949 | 15.7389584 | 1.69E-04 |
| CBR4 | -1.5403395 | 17.7094992 | 3.40E-05 |
| NDUFAF3 | -1.5396641 | 19.8266253 | 2.60E-08 |
| KRT3 | -1.5368201 | 19.7364804 | 0.01570408 |
| PHKA1 | -1.5356072 | 19.8140403 | 0.00151237 |
| SLC25A15 | -1.5345207 | 19.8229975 | 4.87E-07 |
| SLC38A1 | -1.5306502 | 19.192369 | 0.0019123 |
| ADAMTS15 | -1.5286064 | 16.1415912 | 1.17E-06 |
| STOM | -1.5283844 | 17.5198629 | 1.50E-06 |
| TUBB2B | -1.5277439 | 21.9572805 | 0.04357614 |
| HSD17B10 | -1.5256171 | 18.1047755 | 6.73E-05 |

| | | | |
|---|---|---|---|
| PHKB | -1.5214684 | 20.8148639 | 1.29E-08 |
| ARPC5L | -1.5190836 | 16.55958 | 5.80E-07 |
| SLITRK5 | -1.5178765 | 19.0594946 | 1.39E-08 |
| RTN4 | -1.5128937 | 18.2965181 | 3.64E-08 |
| RHBDF1 | -1.5105906 | 16.459507 | 3.21E-07 |
| NDUFA12 | -1.5060887 | 18.3658123 | 1.21E-09 |
| PIP | -1.5053189 | 19.4893207 | 0.00530145 |
| SLC25A5 | -1.496885 | 28.2397401 | 2.80E-08 |
| SLC25A19 | -1.4951348 | 22.0554929 | 6.28E-09 |
| AMIGO1 | -1.4914957 | 16.465872 | 1.15E-06 |
| ATP5MF | -1.4850219 | 22.2035586 | 6.39E-04 |
| GOLPH3 | -1.4835454 | 15.6436363 | 1.60E-06 |
| TBC1D4 | -1.472611 | 17.2445651 | 0.0010637 |
| NDUFS8 | -1.4652163 | 20.8401585 | 1.96E-06 |
| SLC44A1 | -1.4615845 | 16.4758424 | 1.93E-08 |
| SLC25A20 | -1.4601895 | 17.2824435 | 2.91E-06 |
| PYGL | -1.4451011 | 19.5876207 | 0.01963884 |
| SLC25A4 | -1.4433379 | 23.6561579 | 1.40E-08 |
| TIMM23 | -1.4412503 | 20.8975173 | 1.83E-08 |
| VANGL2 | -1.4323227 | 16.0476968 | 4.95E-04 |
| DSC1 | -1.4317459 | 19.455346 | 0.00689375 |
| NDUFS2 | -1.4299613 | 20.9720748 | 1.99E-06 |
| RADX | -1.4298476 | 17.9064986 | 0.01014882 |

| | | | |
|---|---|---|---|
| SEMA3E | -1.4287004 | 16.4868038 | 5.31E-08 |
| SLC25A21 | -1.4206835 | 19.5940098 | 1.33E-08 |
| RPS13 | -1.4185691 | 20.886606 | 5.46E-04 |
| VIM | -1.4151454 | 25.7223452 | 1.49E-06 |
| LATS1 | -1.4134644 | 17.4815562 | 8.08E-07 |
| RHOF | -1.4080809 | 16.8394404 | 6.92E-06 |
| MRPS15 | -1.4058327 | 17.450455 | 2.56E-06 |
| THBS1 | -1.4018126 | 17.5850691 | 1.17E-04 |
| MRM2 | -1.3878228 | 18.2809232 | 6.84E-06 |
| ACOT8 | -1.3857687 | 21.188513 | 2.11E-06 |
| SNU13 | -1.3839021 | 18.065575 | 7.36E-07 |
| SPINT2 | -1.3805737 | 16.502846 | 9.65E-05 |
| ATP5PB | -1.3794075 | 20.7212348 | 6.20E-06 |
| ELSPBP1 | -1.3738005 | 16.5051037 | 1.12E-06 |
| GALNT5 | -1.3721479 | 16.5056546 | 1.09E-05 |
| PODXL | -1.3683397 | 16.506924 | 5.41E-05 |
| C17ORF80 | -1.3678504 | 17.4663515 | 6.75E-09 |
| SLC35E1 | -1.3654913 | 20.6773696 | 0.06189419 |
| GPAT3 | -1.3571233 | 18.1377419 | 2.15E-06 |
| IRS4 | -1.3530647 | 21.8414937 | 9.81E-05 |
| UBE2N;UBE2NL | -1.3514558 | 17.4185135 | 7.35E-08 |
| PGAM5 | -1.3500921 | 21.8519556 | 5.07E-09 |
| NAT14 | -1.3486979 | 17.7628866 | 4.57E-05 |

| | | | |
|---|---|---|---|
| PTGES2 | -1.3458899 | 17.0947696 | 4.13E-06 |
| COG3 | -1.3445811 | 17.4585951 | 1.02E-07 |
| SLC25A25 | -1.3434989 | 18.8514021 | 6.72E-05 |
| DMXL2 | -1.3433404 | 18.0846221 | 2.85E-07 |
| COQ2 | -1.3413887 | 17.457531 | 7.71E-08 |
| DLD | -1.337796 | 17.4563334 | 1.14E-07 |
| RAB39A | -1.3360045 | 19.6547561 | 7.28E-08 |
| RDH10 | -1.3272465 | 16.6634724 | 7.53E-05 |
| AFG2A | -1.3181927 | 17.2197251 | 3.16E-04 |
| TIMMDC1 | -1.3127733 | 22.0908942 | 1.01E-08 |
| SPAG5 | -1.3125207 | 17.6562033 | 3.37E-06 |
| DNAJA1 | -1.3106164 | 24.80791 | 1.01E-07 |
| PDCD10 | -1.3076707 | 16.527147 | 3.09E-06 |
| RHBDD2 | -1.3058935 | 19.6518942 | 4.14E-08 |
| COX7A2 | -1.3024014 | 19.829723 | 9.10E-06 |
| NXF1 | -1.3009578 | 19.1470623 | 2.86E-04 |
| SLC25A6 | -1.2980509 | 20.9112732 | 1.50E-05 |
| SLC25A29 | -1.293569 | 19.1699143 | 5.47E-07 |
| NDUFS7 | -1.2930543 | 20.249624 | 4.37E-05 |
| HSCB | -1.2892161 | 16.6536088 | 9.73E-08 |
| CUL7 | -1.2846807 | 17.5700955 | 1.64E-06 |
| UQCC3 | -1.2823994 | 20.7714097 | 6.55E-08 |
| ZSWIM8 | -1.2816387 | 17.4376143 | 4.02E-07 |

| | | | |
|---|---|---|---|
| SLC25A33 | -1.277031 | 20.3394781 | 1.92E-05 |
| SLC25A28 | -1.2760785 | 16.9245547 | 2.01E-06 |
| PLEKHH3 | -1.2737279 | 16.4133564 | 1.44E-06 |
| MAP7 | -1.2725916 | 17.4405226 | 2.93E-08 |
| GLUD1 | -1.2716249 | 21.0551031 | 6.94E-08 |
| EXOC7 | -1.270882 | 20.2386891 | 0.12289453 |
| CHP1 | -1.2696184 | 20.2368398 | 4.00E-08 |
| CDC27 | -1.2654816 | 17.7658735 | 7.12E-06 |
| NDUFA8 | -1.2639977 | 19.2126814 | 1.68E-08 |
| MUCL1 | -1.2636706 | 17.431625 | 6.95E-04 |
| SEMA3D | -1.2627327 | 16.5421263 | 1.75E-04 |
| MRPL9 | -1.2583601 | 17.8037421 | 1.38E-07 |
| HIGD1A | -1.2551188 | 19.0281865 | 4.42E-08 |
| RP2 | -1.2530768 | 16.8158363 | 5.76E-06 |
| MRPL14 | -1.2529222 | 17.2965629 | 8.67E-04 |
| SPG7 | -1.2526382 | 17.285153 | 4.28E-07 |
| YWHAB | -1.2513261 | 17.4275101 | 2.23E-06 |
| SLC27A6 | -1.2504204 | 18.221093 | 7.78E-06 |
| SSR1 | -1.2450926 | 23.081525 | 1.43E-06 |
| MYO1B | -1.2438677 | 17.4013493 | 1.30E-06 |
| VPS33A | -1.2423389 | 17.0459475 | 2.95E-05 |
| FAM210A | -1.2381909 | 19.8767649 | 4.97E-08 |
| MED1 | -1.2354963 | 17.1594812 | 0.00213908 |

| | | | |
|---|---|---|---|
| SERAC1 | -1.2338586 | 17.8935493 | 2.12E-06 |
| SLC35A5 | -1.2304049 | 16.5529023 | 1.89E-07 |
| RCE1 | -1.2297033 | 16.5531361 | 1.72E-05 |
| KIAA1549 | -1.2296335 | 16.5531594 | 2.83E-07 |
| MRPL24 | -1.2180322 | 17.8359978 | 4.47E-05 |
| MICOS13 | -1.2131933 | 21.1099818 | 6.00E-05 |
| USP11 | -1.2112548 | 16.5592856 | 4.00E-07 |
| ELP6 | -1.2108658 | 16.7486802 | 6.00E-06 |
| CETN3 | -1.2070176 | 17.4127406 | 1.52E-08 |
| SLC25A10 | -1.2055691 | 23.8737018 | 3.21E-07 |
| ARL13B | -1.2049597 | 17.8325665 | 9.24E-07 |
| TRRAP | -1.2047854 | 17.9020246 | 0.00252424 |
| UGDH | -1.2043715 | 15.6141317 | 5.70E-04 |
| MYO9A | -1.1969324 | 17.5093604 | 7.40E-06 |
| NF1 | -1.1961003 | 18.0882982 | 2.85E-04 |
| SLC16A10 | -1.1950498 | 18.5832418 | 5.24E-07 |
| SLC25A14 | -1.194598 | 19.3862351 | 8.39E-05 |
| SYNE3 | -1.191181 | 16.5659769 | 8.41E-08 |
| TMEM165 | -1.1877502 | 19.3614679 | 2.02E-06 |
| RAB6A | -1.1862887 | 17.405831 | 1.39E-08 |
| EXOSC4 | -1.1853658 | 17.0391519 | 0.00297502 |
| LETMD1 | -1.1847926 | 17.0816351 | 4.87E-07 |
| DNAJC25 | -1.182808 | 17.7284575 | 2.42E-06 |

| | | | |
|---|---|---|---|
| DOCK11 | -1.182263 | 19.3312395 | 6.30E-04 |
| PNKD | -1.1822303 | 16.7727639 | 1.18E-08 |
| LGR4 | -1.176234 | 16.5709592 | 1.60E-07 |
| MRPL23 | -1.1715418 | 17.40442 | 2.66E-07 |
| HEATR6 | -1.1702887 | 17.5037854 | 1.73E-06 |
| DYNC2LI1 | -1.1699867 | 17.9899002 | 3.11E-04 |
| GPAT4 | -1.1697892 | 19.4029677 | 3.52E-08 |
| AZGP1 | -1.1677354 | 14.8404116 | 0.03219891 |
| ATP5MG | -1.1643699 | 21.8431706 | 2.24E-08 |
| C3ORF38 | -1.1641649 | 19.9631238 | 0.00279002 |
| MRPL3 | -1.1637415 | 17.3983153 | 2.72E-07 |
| CHAMP1 | -1.1637089 | 17.0156108 | 0.05748924 |
| MSMO1 | -1.1619494 | 16.5757208 | 2.02E-06 |
| HSPD1 | -1.1599098 | 23.6664312 | 2.05E-07 |
| TIMM21 | -1.159328 | 20.1842033 | 7.88E-06 |
| ABCB6 | -1.1488172 | 17.6768574 | 1.53E-05 |
| UVRAG | -1.1472515 | 16.3480345 | 6.68E-07 |
| GPD2 | -1.1465969 | 19.7858132 | 3.34E-06 |
| DDX11 | -1.1457982 | 17.8570928 | 8.26E-07 |
| DENND6A | -1.1450961 | 16.4085911 | 3.94E-07 |
| EEF1B2 | -1.1442831 | 17.2557777 | 0.00126155 |
| RNF167 | -1.1409457 | 18.0144612 | 3.09E-05 |
| MAIP1 | -1.1387619 | 20.8685517 | 0.00337321 |

| | | | |
|---|---|---|---|
| SLC25A26 | -1.1378362 | 16.922251 | 3.98E-08 |
| AHSA1 | -1.1333438 | 19.2882145 | 1.34E-07 |
| CSNK1G3 | -1.1323976 | 16.4867465 | 5.80E-07 |
| ARL1 | -1.1313685 | 21.8768016 | 1.81E-08 |
| TIMM29 | -1.1313593 | 20.6894993 | 2.68E-04 |
| TMEM183A;TMEM183BP | -1.1302659 | 17.1669648 | 1.28E-05 |
| ALG1 | -1.1297253 | 19.7623272 | 2.23E-06 |
| ATP5F1B | -1.1280507 | 20.339986 | 7.32E-06 |
| POR | -1.1239931 | 18.4321842 | 3.73E-07 |
| ESYT2 | -1.1204374 | 21.36949 | 6.38E-07 |
| WDR6 | -1.1145664 | 18.1388057 | 2.57E-05 |
| TMA7;TMA7B | -1.1128271 | 18.2206152 | 0.0096371 |
| HSD17B11 | -1.10946 | 22.153938 | 4.27E-06 |
| RHOT2 | -1.108913 | 18.2392199 | 1.63E-07 |
| APOD | -1.1084271 | 17.3106428 | 8.43E-07 |
| IGHA1;IGHA2 | -1.1083168 | 18.2594642 | 0.01418213 |
| SLC25A23 | -1.107323 | 17.7885894 | 1.84E-07 |
| RPS3 | -1.1049479 | 25.9355767 | 6.96E-08 |
| POLR3B | -1.1035215 | 18.3805342 | 0.00127764 |
| RPL4 | -1.103158 | 24.0275521 | 4.59E-04 |
| TCP11L1 | -1.1027785 | 17.3863563 | 3.98E-08 |
| POMGNT1 | -1.1015814 | 18.7704597 | 1.48E-07 |
| EDF1 | -1.1006207 | 17.5128982 | 9.54E-08 |

| | | | |
|---|---|---|---|
| GOLGA7 | -1.1001028 | 16.4552358 | 1.09E-07 |
| ATP5F1D | -1.0926884 | 17.5839242 | 5.82E-04 |
| GTF2H4 | -1.0907712 | 17.7081456 | 3.91E-08 |
| KDELR2 | -1.0890719 | 18.3054131 | 0.01517073 |
| MAD2L2 | -1.0870097 | 17.372738 | 2.82E-08 |
| N4BP3 | -1.0859347 | 16.8500843 | 4.06E-05 |
| STAG1 | -1.0856631 | 16.8883108 | 5.73E-04 |
| KEAP1 | -1.084778 | 16.7265687 | 1.81E-05 |
| SFXN1 | -1.0844933 | 24.0881434 | 4.21E-08 |
| BMP8A;BMP8B | -1.0822634 | 16.6022828 | 1.69E-04 |
| PIGQ | -1.0812505 | 20.2284895 | 8.32E-06 |
| TLN1 | -1.0772849 | 19.7982221 | 3.79E-04 |
| VPS50 | -1.0772274 | 16.8639808 | 1.73E-05 |
| NDUFV1 | -1.0762229 | 19.1350474 | 2.32E-06 |
| DECR2 | -1.0752292 | 17.6779544 | 9.02E-06 |
| NOC2L | -1.0748296 | 17.4295512 | 5.38E-05 |
| IMPA1 | -1.0744591 | 18.105748 | 1.11E-06 |
| BSG | -1.0741516 | 20.7502886 | 1.78E-07 |
| ACSL3 | -1.0739497 | 21.1211074 | 6.26E-06 |
| COX10 | -1.0727064 | 16.7053706 | 5.82E-04 |
| ABCC4 | -1.072434 | 16.6369221 | 9.49E-06 |
| KIF14 | -1.0713269 | 19.4723563 | 8.61E-07 |
| LPCAT1 | -1.0691612 | 18.8697563 | 4.53E-06 |

| | | | |
|---|---|---|---|
| MTHFD1L | -1.0691218 | 17.3557124 | 0.00486505 |
| SLC16A1 | -1.0688756 | 24.6920841 | 6.21E-07 |
| NDUFB1 | -1.065496 | 17.2751684 | 1.46E-06 |
| PTCD2 | -1.064947 | 15.6761489 | 0.0032062 |
| EIF2B2 | -1.0644181 | 16.6082312 | 9.10E-07 |
| POLD3 | -1.0640745 | 17.3650929 | 2.13E-08 |
| FASTKD5 | -1.0637489 | 20.9776244 | 1.01E-04 |
| RAB10 | -1.0620475 | 23.0951023 | 3.76E-06 |
| ECSIT | -1.060455 | 17.0262704 | 1.47E-07 |
| MRPL41 | -1.0599911 | 18.2181603 | 7.76E-06 |
| ATP2C1 | -1.0598196 | 18.800828 | 4.85E-07 |
| PCDH9 | -1.0588737 | 15.8562153 | 1.10E-05 |
| VAMP7 | -1.0575322 | 17.7524776 | 4.08E-07 |
| RAF1 | -1.0558619 | 21.1610268 | 6.82E-05 |
| OSGEP | -1.0556582 | 17.4364476 | 6.17E-05 |
| NUP188 | -1.0529073 | 19.3915907 | 2.71E-06 |
| IBA57 | -1.0520618 | 19.5119677 | 9.86E-08 |
| ACSL4 | -1.051488 | 18.8042305 | 0.00139766 |
| DNAJA3 | -1.0503656 | 20.9474307 | 1.23E-05 |
| COA3 | -1.0494764 | 21.1045451 | 5.57E-04 |
| FEZ2 | -1.0476582 | 19.0108776 | 5.93E-04 |
| RPL34 | -1.046413 | 21.9646702 | 0.00103545 |
| DHRS7B | -1.0453186 | 18.8267132 | 8.08E-07 |

| | | | |
|---|---|---|---|
| PARL | -1.0427059 | 21.275181 | 1.21E-04 |
| HRNR | -1.0356733 | 21.5703234 | 0.00720717 |
| ACSL1 | -1.0337483 | 18.3763357 | 1.81E-06 |
| PUS7 | -1.032669 | 18.1539925 | 0.00472623 |
| ABHD14B | -1.0322559 | 18.7475059 | 1.25E-05 |
| DENND4C | -1.0309866 | 18.3095219 | 8.26E-07 |
| CMTM8 | -1.029762 | 18.8624534 | 1.06E-07 |
| MBOAT1 | -1.0280692 | 17.4850875 | 9.36E-06 |
| SLC25A38 | -1.0278313 | 17.275179 | 5.29E-04 |
| HSD17B12 | -1.0259428 | 23.0142035 | 2.12E-06 |
| GCAT | -1.0240696 | 19.4120007 | 4.80E-06 |
| ERCC3 | -1.0220598 | 18.4809398 | 9.94E-05 |
| KRT9 | -1.0198833 | 25.9832291 | 0.00592011 |
| RAB31 | -1.0197976 | 15.1897109 | 0.00808987 |
| SAMD1 | -1.0188787 | 17.1396824 | 3.14E-05 |
| ATP5PD | -1.0183611 | 19.2868575 | 3.78E-07 |
| MAPK14 | -1.0144585 | 14.6327259 | 0.00307389 |
| MRPL45 | -1.0140527 | 18.1515479 | 9.39E-04 |
| DECR1 | -1.0097362 | 17.132254 | 5.62E-05 |
| CDC45 | -1.0094799 | 19.3055127 | 0.0011681 |
| SLC35C1 | -1.009361 | 17.9705278 | 2.66E-07 |
| MAP7D3 | -1.0074023 | 16.7635848 | 4.27E-04 |
| MAP1B | -1.0064147 | 16.4758945 | 5.25E-04 |

| | | | |
|---|---|---|---|
| IMPA2 | -1.0060351 | 17.6869333 | 0.00168294 |
| SLC25A39 | -1.0034417 | 16.4126751 | 2.88E-05 |
| CDIPT | -1.0032212 | 22.2294178 | 1.59E-05 |
| SLC25A1 | -1.0004204 | 25.1668241 | 1.85E-07 |

### TMED7 preferred

| Gene | logFC | Average expression | Adj. p value |
|---|---|---|---|
| METTL2A;METTL2B | 8.21700968 | 22.3899625 | 1.07E-12 |
| TMED7 | 7.11807398 | 27.6029425 | 2.51E-10 |
| LRPAP1 | 5.36975361 | 19.6284435 | 2.59E-12 |
| PBXIP1 | 5.26383896 | 18.7176502 | 5.33E-13 |
| SMIM14 | 4.80819702 | 16.8380425 | 2.78E-12 |
| KRT13 | 4.77975557 | 20.3295871 | 6.93E-09 |
| CLTB | 4.70859607 | 18.3319398 | 1.68E-13 |
| TUSC3 | 4.55125817 | 19.3197672 | 7.79E-10 |
| SIGMAR1 | 4.49796977 | 18.4623605 | 1.35E-12 |
| KLHL22 | 4.43995578 | 18.4430225 | 1.11E-12 |
| JMJD8 | 4.400829 | 18.4299802 | 3.92E-11 |
| TOR3A | 4.35568085 | 19.0362503 | 1.82E-07 |
| CLTA | 4.30332696 | 18.9650941 | 3.79E-13 |
| FZD2 | 4.1621943 | 18.6862993 | 3.79E-13 |
| FZD1 | 4.14343787 | 18.3441832 | 1.41E-12 |

| | | | |
|---|---|---|---|
| MANF | 4.12846547 | 19.1813371 | 3.15E-10 |
| FRAS1 | 4.12555613 | 18.3382226 | 9.33E-12 |
| CLCC1 | 4.11573869 | 19.0131542 | 1.89E-13 |
| ICAM5 | 4.0860176 | 18.3250431 | 4.07E-12 |
| NICOL1 | 4.00643922 | 18.298517 | 4.38E-10 |
| HAPLN3 | 3.97527188 | 18.6283353 | 1.01E-08 |
| NRM | 3.96142957 | 18.2835138 | 1.87E-12 |
| DNAJB9 | 3.85303707 | 18.2473829 | 2.22E-12 |
| CLINT1 | 3.793368 | 21.0465685 | 2.94E-11 |
| NAGLU | 3.78235231 | 18.6021395 | 3.67E-04 |
| SUN1 | 3.75306516 | 19.3572211 | 2.12E-11 |
| TPST2 | 3.7202641 | 17.1726866 | 1.28E-04 |
| MIA3 | 3.70690039 | 18.1986707 | 3.85E-12 |
| TMED3 | 3.68618546 | 21.0211681 | 9.19E-13 |
| MARCHF6 | 3.67249421 | 18.187202 | 4.81E-12 |
| CDK5RAP1 | 3.65199941 | 18.9425166 | 4.81E-12 |
| GALNS | 3.58865519 | 18.1592556 | 1.38E-10 |
| SLC39A1 | 3.5599344 | 18.3282862 | 1.57E-12 |
| ARSK | 3.55418281 | 18.1477648 | 4.81E-12 |
| IKBIP | 3.53897734 | 18.1426963 | 2.55E-11 |
| OS9 | 3.53577251 | 19.0035439 | 1.90E-10 |
| SUMF2 | 3.53100781 | 19.0574818 | 4.67E-12 |
| MAGT1 | 3.52045597 | 23.3532047 | 2.41E-06 |

| | | | |
|---|---|---|---|
| PLPP5 | 3.51468702 | 18.1345996 | 6.73E-11 |
| TMEM87A | 3.46615095 | 18.1184209 | 6.44E-11 |
| ATP11C | 3.34168504 | 18.7780408 | 2.86E-10 |
| GPM6A | 3.26097764 | 18.0500298 | 1.02E-11 |
| MTFR1 | 3.24798086 | 18.0456975 | 7.76E-12 |
| CRELD2 | 3.24268866 | 18.0439335 | 2.10E-11 |
| HLA-E | 3.22153982 | 18.0368838 | 1.18E-11 |
| SLC35C2 | 3.21273477 | 18.0339488 | 1.85E-10 |
| NGLY1 | 3.19978409 | 18.0296319 | 2.46E-11 |
| ATP6V0A2 | 3.18718043 | 18.0254307 | 8.14E-11 |
| CCDC134 | 3.17721596 | 18.9848442 | 8.35E-10 |
| KRT4 | 3.17295749 | 19.539406 | 4.87E-06 |
| LMAN1 | 3.17290572 | 20.9539782 | 1.22E-11 |
| ASAH1 | 3.14437313 | 18.1772848 | 3.85E-12 |
| TMEM30A | 3.12431525 | 19.5787952 | 2.37E-11 |
| GPC5 | 3.10227822 | 17.8519793 | 1.91E-11 |
| SLC12A9 | 3.09490365 | 18.9036797 | 3.66E-12 |
| APPBP2 | 3.09326708 | 17.9941263 | 2.46E-11 |
| CDH2 | 3.08704011 | 19.1588442 | 1.53E-10 |
| CCDC167 | 3.08173546 | 17.9668828 | 6.74E-12 |
| GLT8D2 | 3.08036701 | 17.9898262 | 1.83E-10 |
| YIPF3 | 3.0765694 | 17.9885604 | 1.22E-11 |
| ALG9 | 3.06799887 | 19.2432174 | 1.72E-12 |

| | | | |
|---|---|---|---|
| CTNNB1 | 3.06427331 | 21.4337928 | 4.02E-08 |
| MSRB3 | 3.03993864 | 17.4909104 | 2.12E-11 |
| CREB3L2 | 3.03681572 | 18.4969679 | 2.22E-12 |
| SEC63 | 3.01040811 | 20.2971599 | 1.30E-11 |
| CCN1 | 2.97475415 | 17.9546219 | 4.70E-11 |
| UBE2J1 | 2.97420541 | 20.1322008 | 1.81E-10 |
| FTO | 2.95281098 | 15.8309961 | 3.18E-09 |
| PRKACB | 2.95261337 | 17.9472417 | 2.43E-11 |
| MFSD5 | 2.94621422 | 17.9451086 | 3.74E-10 |
| ATP6V0A1 | 2.91499708 | 18.8021519 | 3.34E-11 |
| ATP6AP1 | 2.90654383 | 19.8778673 | 3.85E-12 |
| CANX | 2.88284985 | 22.1746836 | 2.82E-09 |
| GPC6 | 2.87882894 | 21.0574642 | 1.53E-10 |
| PDIA6 | 2.87219034 | 21.9047736 | 7.46E-06 |
| PCNX3 | 2.85956211 | 17.8666641 | 2.82E-09 |
| CKAP4 | 2.85938793 | 20.9500003 | 7.49E-06 |
| NUDT9 | 2.85354733 | 18.7708405 | 5.23E-10 |
| LIPA | 2.83224074 | 17.9071175 | 3.13E-10 |
| ITGB1 | 2.82182678 | 17.9036462 | 2.56E-11 |
| ULBP3 | 2.8122943 | 17.946106 | 4.69E-11 |
| PPM1L | 2.79817526 | 17.5353799 | 7.98E-05 |
| NCSTN | 2.79066799 | 20.5170836 | 4.18E-12 |
| BLZF1 | 2.78822557 | 17.8924458 | 3.14E-11 |

| | | | |
|---|---|---|---|
| XXYLT1 | 2.74673484 | 17.8786155 | 1.83E-10 |
| PDIA3 | 2.74471013 | 20.2722361 | 3.65E-10 |
| SLC9A8 | 2.74211647 | 18.6678988 | 3.66E-12 |
| FZD3 | 2.74190991 | 18.3219754 | 2.51E-11 |
| ERP29 | 2.73919925 | 18.8240222 | 4.37E-09 |
| AGA | 2.73737025 | 17.6787033 | 2.73E-10 |
| SELENOM | 2.73049877 | 18.8505232 | 2.18E-09 |
| CHMP2B | 2.72530013 | 17.8714706 | 2.00E-10 |
| EDEM1 | 2.71738774 | 18.0599674 | 4.42E-12 |
| NCBP2AS2 | 2.69163699 | 18.9886305 | 2.51E-07 |
| HEXB | 2.67331538 | 18.1504502 | 3.18E-08 |
| FAM241B | 2.66741755 | 17.8521764 | 1.28E-10 |
| LAMB2 | 2.66627935 | 18.7259229 | 4.13E-07 |
| SLC2A6 | 2.66241303 | 17.8505082 | 6.83E-09 |
| STC2 | 2.65344729 | 21.4863447 | 2.37E-11 |
| SLC41A3 | 2.6446557 | 17.8445891 | 6.44E-11 |
| NPC1 | 2.6030814 | 19.0296269 | 2.51E-11 |
| DCBLD1 | 2.59016084 | 17.8264242 | 5.83E-11 |
| NECTIN2 | 2.58963581 | 18.5978684 | 1.60E-10 |
| TMEM259 | 2.58521605 | 17.8247759 | 6.44E-11 |
| CTNNA1 | 2.58489844 | 20.1380185 | 1.09E-10 |
| MXRA8 | 2.58303929 | 17.5659401 | 2.86E-10 |
| CDC20 | 2.57467938 | 18.0966579 | 9.07E-08 |

| | | | |
|---|---|---|---|
| BRI3BP | 2.57046046 | 19.6344336 | 4.82E-10 |
| PTK7 | 2.55747367 | 19.800444 | 4.66E-09 |
| SIL1 | 2.55286676 | 18.7734837 | 1.54E-11 |
| WFS1 | 2.54572304 | 20.4427698 | 1.91E-10 |
| SEL1L | 2.52701485 | 20.5116048 | 3.60E-08 |
| METRNL | 2.52487788 | 17.8046632 | 6.93E-11 |
| POGLUT1 | 2.52259448 | 18.7615025 | 1.37E-07 |
| ERP44 | 2.52044841 | 20.319254 | 6.22E-11 |
| MBTPS2 | 2.51404632 | 17.8010527 | 1.13E-09 |
| HLA-C | 2.49698919 | 19.3869797 | 5.32E-09 |
| DCBLD2 | 2.48961767 | 17.7309778 | 2.40E-10 |
| TAP2 | 2.47615525 | 19.5284123 | 5.94E-07 |
| CALR | 2.46516593 | 20.4993676 | 1.39E-08 |
| GRAMD1B | 2.4621048 | 16.9318595 | 3.14E-11 |
| NEMP1 | 2.46142769 | 20.6525562 | 3.85E-11 |
| SLC52A2 | 2.4556262 | 17.9523715 | 1.36E-07 |
| CCPG1 | 2.45331462 | 19.7038217 | 3.85E-11 |
| ALG12 | 2.43582276 | 18.6069497 | 3.90E-10 |
| CXADR | 2.43397778 | 17.7743632 | 9.37E-09 |
| CNNM1 | 2.43213988 | 17.7737505 | 2.86E-10 |
| POFUT2 | 2.42992178 | 18.0245122 | 1.07E-10 |
| ITGA5 | 2.4218459 | 18.1194605 | 2.78E-11 |
| MENT | 2.41654463 | 17.5426719 | 4.45E-11 |

| | | | |
|---|---|---|---|
| SYVN1 | 2.4161328 | 17.7684148 | 1.85E-10 |
| MANEA | 2.41286155 | 19.6620631 | 5.85E-11 |
| C1GALT1 | 2.40632767 | 17.7651465 | 1.50E-10 |
| HSP90B1 | 2.39378578 | 23.2394903 | 3.13E-10 |
| SLC4A2 | 2.38673014 | 18.9902441 | 9.85E-04 |
| ANG | 2.37464394 | 17.7545852 | 4.82E-10 |
| TMX1 | 2.3606911 | 19.6334966 | 1.18E-11 |
| RGPD4 | 2.34656443 | 17.7452254 | 4.76E-09 |
| PIGG | 2.34590193 | 19.2759268 | 4.45E-11 |
| LSR | 2.34556563 | 18.6382817 | 1.22E-10 |
| MAN1A1 | 2.3396001 | 20.2386898 | 1.83E-08 |
| MRPL21 | 2.32408771 | 16.2357055 | 2.51E-11 |
| ATP6AP2 | 2.3174445 | 19.2790568 | 1.46E-10 |
| UBR4 | 2.31620074 | 21.6460866 | 6.87E-12 |
| COL18A1 | 2.31006357 | 18.6162245 | 0.00100403 |
| PTGFRN | 2.30510653 | 18.2824965 | 1.72E-09 |
| PIGO | 2.29374998 | 19.3148242 | 1.61E-09 |
| MPZL1 | 2.28565159 | 16.2485176 | 3.14E-11 |
| ATP11B | 2.2766472 | 17.7219196 | 1.35E-09 |
| MIB2 | 2.27218529 | 17.7204323 | 1.85E-10 |
| ZNF391 | 2.27197875 | 17.7203635 | 1.19E-08 |
| INHBE | 2.270989 | 19.6607154 | 2.16E-10 |
| P3H1 | 2.27037608 | 19.0495001 | 2.85E-09 |

| | | | |
|---|---|---|---|
| CHST10 | 2.26298213 | 18.2963624 | 4.46E-11 |
| HLA-DQB1 | 2.25937413 | 17.5919505 | 3.14E-11 |
| TRPM7 | 2.25284226 | 17.8979527 | 1.74E-08 |
| BPNT2 | 2.25236364 | 19.4864431 | 1.76E-07 |
| NAXE | 2.23280834 | 17.7073067 | 4.22E-09 |
| ATP1B3 | 2.22412102 | 20.9938714 | 4.11E-07 |
| TOR1A | 2.22410777 | 19.0546144 | 1.00E-08 |
| HLA-B | 2.21802802 | 21.3917703 | 3.53E-11 |
| SP9 | 2.21573865 | 17.7016168 | 1.02E-09 |
| TEX264 | 2.21071994 | 17.6999439 | 1.54E-09 |
| ITGAV | 2.20247812 | 18.5654016 | 2.16E-10 |
| ERLEC1 | 2.20207986 | 20.4944929 | 1.22E-10 |
| PCBD1;PCBD2 | 2.19528236 | 18.451992 | 2.86E-10 |
| LAMP1 | 2.18890593 | 18.8210881 | 2.34E-09 |
| LRP11 | 2.18812376 | 17.0921604 | 5.91E-07 |
| NMU | 2.18257801 | 18.2055242 | 5.58E-07 |
| TXNDC15 | 2.17197983 | 18.5816883 | 9.87E-10 |
| C1GALT1C1 | 2.16624106 | 19.8459323 | 4.45E-11 |
| CLSTN1 | 2.14890793 | 17.5638729 | 4.82E-08 |
| CTSV | 2.14080119 | 17.8635652 | 1.08E-08 |
| TAP1 | 2.13835098 | 19.6342026 | 3.61E-11 |
| CLTC | 2.13458752 | 22.8576722 | 1.52E-07 |
| SASS6 | 2.12793329 | 18.0308213 | 4.65E-10 |

| | | | |
|---|---|---|---|
| LRIG2 | 2.12158854 | 17.7481759 | 3.53E-11 |
| MFGE8 | 2.11899611 | 20.7015897 | 4.63E-09 |
| P4HA2 | 2.11327294 | 18.4986814 | 2.17E-08 |
| RB1CC1 | 2.11186525 | 17.9687163 | 3.72E-10 |
| DGCR2 | 2.10432176 | 18.9265392 | 4.50E-09 |
| LAMB1 | 2.09858895 | 18.559772 | 0.02368873 |
| B3GALT6 | 2.09337958 | 20.0354528 | 6.50E-08 |
| TAPBP | 2.09054018 | 18.0343665 | 2.02E-07 |
| CRTAP | 2.08981453 | 19.383848 | 3.55E-09 |
| TMX4 | 2.08719677 | 19.5924634 | 7.81E-10 |
| GPC4 | 2.08378149 | 20.5236547 | 1.55E-10 |
| B3GLCT | 2.07733917 | 18.9449614 | 4.38E-10 |
| POFUT1 | 2.07640446 | 19.2167237 | 1.42E-07 |
| TXNDC5 | 2.07600652 | 21.533692 | 1.54E-04 |
| P3H4 | 2.07594117 | 19.5227098 | 1.90E-09 |
| DDR2 | 2.07498898 | 18.1438824 | 2.96E-10 |
| KCMF1 | 2.06853445 | 17.0450918 | 2.65E-06 |
| SULF1 | 2.06627934 | 18.5262455 | 9.94E-05 |
| TMEM39A | 2.06481403 | 20.1821349 | 3.19E-08 |
| CHD7;CHD9 | 2.06079085 | 16.395328 | 3.09E-11 |
| WNT5B | 2.05880367 | 17.8203979 | 1.55E-10 |
| SEC11A | 2.05807276 | 21.1240926 | 6.01E-10 |
| DPY19L4 | 2.05692367 | 17.6486785 | 2.47E-07 |

| | | | |
|---|---|---|---|
| TXNDC12 | 2.05411986 | 19.0315987 | 3.73E-10 |
| SPPL2B | 2.05401786 | 18.4941528 | 7.08E-09 |
| MOXD1 | 2.04455231 | 20.068783 | 8.20E-10 |
| MPZL3 | 2.0420745 | 17.9422038 | 3.89E-10 |
| RBM41 | 2.02844816 | 15.9453993 | 6.40E-08 |
| ADAM10 | 2.02379082 | 20.1625861 | 6.93E-11 |
| HLA-A | 2.02378953 | 23.1285949 | 2.11E-08 |
| CNOT2 | 2.00937101 | 17.9512403 | 1.90E-09 |
| TMEM43 | 1.98790169 | 21.9213659 | 3.52E-07 |
| HHIP | 1.98252578 | 16.3790901 | 1.13E-06 |
| FKBP7 | 1.97763212 | 17.6222479 | 2.21E-09 |
| STIM1 | 1.96397162 | 19.5589541 | 1.33E-06 |
| NMB | 1.96254447 | 19.5624089 | 6.76E-09 |
| SMTN | 1.96167012 | 18.0681224 | 8.70E-07 |
| TCTN3 | 1.95233182 | 17.6138145 | 1.54E-09 |
| ACVR2A | 1.95203186 | 17.7015776 | 1.03E-09 |
| PRKCSH | 1.95163814 | 19.0574542 | 7.25E-07 |
| GJA1 | 1.94490029 | 18.8522228 | 6.31E-08 |
| NUCB2 | 1.94470451 | 17.6112721 | 2.60E-08 |
| FRYL | 1.9375627 | 16.3645472 | 4.82E-10 |
| B4GALT3 | 1.93142496 | 18.2235738 | 1.58E-09 |
| TMEM219 | 1.93097231 | 18.4319081 | 1.12E-09 |
| SLC47A1 | 1.93088512 | 17.8281462 | 1.74E-09 |

| | | | |
|---|---|---|---|
| NDFIP2 | 1.92526439 | 16.6151794 | 3.49E-07 |
| SPON1 | 1.92505508 | 19.0456795 | 3.82E-10 |
| ART5 | 1.91751884 | 18.1590519 | 7.50E-10 |
| WNT5A | 1.91504055 | 18.6393058 | 3.17E-06 |
| MFSD14B | 1.90574927 | 17.598287 | 1.70E-09 |
| GPR89A;GPR89B | 1.89549872 | 20.927104 | 6.12E-10 |
| DAG1 | 1.89528184 | 18.9711142 | 3.03E-09 |
| SEL1L3 | 1.8951318 | 18.0328045 | 2.40E-07 |
| NDST1 | 1.88960689 | 18.2582239 | 2.06E-09 |
| SGCB | 1.88827443 | 18.6752155 | 3.69E-09 |
| COL14A1 | 1.8879755 | 17.5923624 | 9.59E-10 |
| UGGT1 | 1.87756107 | 22.2477201 | 1.19E-09 |
| NRP2 | 1.87743807 | 17.437555 | 4.14E-09 |
| CHID1 | 1.87172541 | 21.1542894 | 4.98E-07 |
| CYB5D2 | 1.86754584 | 19.1486177 | 2.96E-09 |
| LRRC8D | 1.86698656 | 17.5853661 | 3.85E-09 |
| MAP3K5 | 1.86695198 | 16.3880841 | 8.40E-09 |
| FAT1 | 1.86171804 | 17.5836099 | 1.36E-05 |
| DPY19L3 | 1.86166831 | 17.7014855 | 5.76E-09 |
| SLC39A6 | 1.85284729 | 18.8880971 | 7.05E-09 |
| ASIC1 | 1.84540443 | 19.3467849 | 3.36E-08 |
| DPY19L1 | 1.84421757 | 19.0109021 | 8.06E-10 |
| PCYOX1L | 1.84249306 | 17.5772016 | 2.20E-04 |

| | | | |
|---|---|---|---|
| GPAA1 | 1.83748098 | 20.1690913 | 4.36E-09 |
| CLGN | 1.83461838 | 20.3199945 | 1.85E-10 |
| OGFOD3 | 1.83159691 | 20.7550264 | 9.56E-09 |
| FAM3C | 1.83139941 | 18.1792174 | 5.30E-08 |
| CHST6 | 1.82927943 | 18.0809262 | 8.01E-07 |
| SLC35E2B | 1.82600081 | 17.5717042 | 4.16E-09 |
| NOMO1 | 1.82568263 | 21.4188792 | 1.96E-04 |
| SNX3 | 1.8235849 | 17.6473685 | 6.68E-09 |
| SOAT1 | 1.82310023 | 19.1590841 | 1.80E-09 |
| GCLM | 1.8215007 | 17.9628914 | 7.15E-04 |
| IL17RA | 1.82111089 | 17.5958655 | 2.26E-10 |
| DNAJC10 | 1.81909909 | 20.7252854 | 6.01E-10 |
| AACS | 1.81809675 | 17.5690695 | 1.51E-09 |
| WLS | 1.81667263 | 20.7518924 | 6.01E-10 |
| TXNDC11 | 1.81480407 | 18.9775896 | 1.99E-09 |
| GRAMD1A | 1.81413423 | 20.2416485 | 1.51E-08 |
| SLC7A8 | 1.80763417 | 17.565582 | 7.03E-09 |
| CNNM4 | 1.7941518 | 19.5847393 | 2.06E-09 |
| PAFAH1B1 | 1.79064304 | 21.6353337 | 1.99E-09 |
| CLPTM1 | 1.79025179 | 19.5604065 | 5.41E-10 |
| DHFR2 | 1.7855471 | 19.3716412 | 2.11E-09 |
| WDR70 | 1.78489891 | 17.5706289 | 7.20E-08 |
| SUN2 | 1.78326216 | 20.9763512 | 7.23E-08 |

| | | | |
|---|---|---|---|
| SUMO1 | 1.77933108 | 16.4749725 | 6.16E-09 |
| ZZEF1 | 1.77755623 | 16.4178827 | 2.06E-07 |
| TMEM87B | 1.77508923 | 19.2144653 | 1.80E-10 |
| MESD | 1.77347084 | 19.4249487 | 4.65E-10 |
| SLC39A7 | 1.77225537 | 20.0991537 | 8.36E-10 |
| PTX3 | 1.77174475 | 17.5536188 | 4.64E-09 |
| SGMS2 | 1.7581194 | 17.339404 | 4.14E-09 |
| SLC12A2 | 1.75160865 | 19.2928548 | 5.14E-10 |
| TMEM231 | 1.75098783 | 19.4740979 | 6.51E-10 |
| TMEM185A | 1.74844496 | 16.6496658 | 9.27E-08 |
| SERPINF1 | 1.73630738 | 17.4119035 | 1.73E-04 |
| CLSTN3 | 1.73179654 | 17.7926934 | 1.06E-08 |
| STT3A | 1.73175699 | 22.3375765 | 1.37E-09 |
| LNPEP | 1.72355927 | 19.1062484 | 7.09E-09 |
| TGFBR1 | 1.72264556 | 19.900001 | 1.05E-06 |
| PACC1 | 1.72134047 | 20.039669 | 4.69E-10 |
| ALG10;ALG10B | 1.72024198 | 20.4463697 | 4.82E-10 |
| HSBP1 | 1.7202061 | 16.7715269 | 2.48E-08 |
| HEXA | 1.71028909 | 17.5331336 | 1.18E-08 |
| UGT3A2 | 1.70765334 | 17.670073 | 3.40E-10 |
| CTSL | 1.70302592 | 17.5307125 | 7.07E-08 |
| PGAP1 | 1.69483371 | 19.9892575 | 4.37E-05 |
| TTC13 | 1.69440786 | 19.4062385 | 2.50E-09 |

| | | | |
|---|---|---|---|
| DSG2 | 1.69198749 | 20.0961833 | 1.80E-08 |
| IL6ST | 1.69038848 | 17.3061725 | 9.03E-09 |
| G3BP1 | 1.69030168 | 19.444832 | 0.00671149 |
| MATN2 | 1.68927806 | 17.5261299 | 1.16E-08 |
| PLAT | 1.68843595 | 20.09451 | 1.61E-07 |
| PIGK | 1.68562571 | 20.8503223 | 4.95E-05 |
| EMILIN2 | 1.68399239 | 19.4376446 | 4.31E-09 |
| ALG8 | 1.67822358 | 21.0759474 | 6.68E-09 |
| PXDN | 1.67787446 | 19.7726895 | 1.16E-08 |
| FAM3A | 1.67721192 | 20.3394643 | 6.24E-10 |
| LONRF3 | 1.67424882 | 17.593243 | 3.87E-08 |
| SLC25A53 | 1.67343268 | 16.4525905 | 1.28E-08 |
| CHPF | 1.67334001 | 21.0646232 | 4.31E-09 |
| IGF1R | 1.67181709 | 19.0092775 | 1.65E-09 |
| IGSF1 | 1.66864317 | 18.4738332 | 2.05E-08 |
| CIP2A | 1.66473317 | 19.9747467 | 1.19E-08 |
| COLEC12 | 1.66369903 | 17.5176036 | 6.19E-09 |
| TRHDE | 1.6571562 | 16.5492088 | 5.96E-08 |
| TMEM120B | 1.65392861 | 18.8559583 | 0.00117279 |
| TMEM106C | 1.65309052 | 17.7091171 | 6.94E-07 |
| TOR1AIP2 | 1.65019514 | 20.9241727 | 7.88E-06 |
| SGSH | 1.64972456 | 17.7135594 | 9.21E-07 |
| TMEM131 | 1.64888601 | 21.3150221 | 2.76E-08 |

| | | | |
|---|---|---|---|
| PIGU | 1.64884574 | 19.2796694 | 8.06E-06 |
| HS6ST2 | 1.64722332 | 20.9634667 | 4.39E-09 |
| HS3ST3A1 | 1.64100423 | 16.2671776 | 1.74E-08 |
| RHOG | 1.63894511 | 16.4640864 | 4.82E-10 |
| MAP7D2 | 1.63780624 | 17.9494697 | 8.52E-07 |
| UBE2D2;UBE2D3 | 1.63643206 | 18.8520986 | 0.00356638 |
| DNAJC16 | 1.63611127 | 20.1319572 | 1.98E-08 |
| GALNT2 | 1.63218273 | 20.5196623 | 2.19E-09 |
| MGAT1 | 1.63151728 | 19.7973735 | 2.85E-07 |
| STIM2 | 1.63055306 | 18.3973707 | 1.96E-06 |
| GPR180 | 1.63037018 | 18.0025654 | 6.35E-06 |
| SLC6A15 | 1.63027095 | 17.6101982 | 1.97E-09 |
| EIF2AK3 | 1.62513583 | 19.5345619 | 5.82E-10 |
| PIGS | 1.62505881 | 20.5800138 | 4.38E-10 |
| PTPRS | 1.61977158 | 19.1505256 | 7.69E-09 |
| FICD | 1.61974196 | 17.5029512 | 1.47E-07 |
| TMEM35B | 1.61934271 | 15.5626535 | 3.08E-08 |
| TOR2A | 1.6164548 | 16.9821772 | 6.11E-08 |
| SDF4 | 1.61414302 | 22.3295335 | 5.85E-07 |
| TMEM214 | 1.60466584 | 17.4979258 | 7.24E-08 |
| SEMA3A | 1.60129783 | 18.1696365 | 7.69E-09 |
| SCPEP1 | 1.59969621 | 18.4200689 | 7.24E-07 |
| ECE1 | 1.581713 | 17.4902749 | 4.62E-08 |

| | | | |
|---|---|---|---|
| HLA-DRB1 | 1.57571104 | 19.8694587 | 4.37E-09 |
| ADAMTS2 | 1.5748628 | 17.4879915 | 5.20E-07 |
| NXPH4 | 1.57169417 | 18.9715716 | 2.36E-09 |
| SLC38A7 | 1.56314157 | 18.2371125 | 4.43E-08 |
| MFSD10 | 1.56275186 | 18.0956013 | 1.30E-06 |
| PRCP | 1.56086693 | 18.172017 | 6.29E-06 |
| CHST11 | 1.55863324 | 19.0614969 | 6.75E-09 |
| SFRP2 | 1.55849907 | 17.4825369 | 4.84E-08 |
| ERO1B | 1.55643024 | 19.2823025 | 8.66E-07 |
| EXT2 | 1.5562726 | 20.1015707 | 0.00109012 |
| CRISPLD1 | 1.55440487 | 18.0885994 | 4.22E-09 |
| SLC33A1 | 1.55303586 | 17.7277675 | 1.96E-06 |
| BCAP31 | 1.54985399 | 19.1341951 | 4.97E-08 |
| PIGT | 1.54787042 | 20.4543162 | 4.58E-08 |
| SMOC1 | 1.54588027 | 19.5400479 | 3.64E-07 |
| KIAA2013 | 1.54459004 | 19.4420268 | 0.00138957 |
| ST6GALNAC4 | 1.54401653 | 19.966956 | 7.88E-10 |
| BMP2 | 1.5350821 | 17.4747313 | 1.03E-08 |
| GPX8 | 1.53364252 | 21.8368644 | 4.40E-09 |
| LRFN3 | 1.53186787 | 17.6758786 | 1.52E-09 |
| FBN2 | 1.52508137 | 20.4122351 | 3.95E-10 |
| CD55 | 1.51938625 | 17.5828617 | 2.60E-07 |
| RANBP2 | 1.51841266 | 20.5315584 | 9.91E-07 |

| | | | |
|---|---|---|---|
| EMC7 | 1.51830896 | 19.8545027 | 5.26E-08 |
| PCSK7 | 1.5166353 | 17.4685823 | 1.56E-07 |
| LOX | 1.50724224 | 17.5499728 | 8.89E-08 |
| HUWE1 | 1.50521768 | 19.9263175 | 2.12E-06 |
| PPIL3 | 1.50459225 | 16.5088707 | 4.25E-08 |
| MLF1 | 1.50350946 | 19.1545666 | 3.91E-09 |
| MBTPS1 | 1.49959004 | 18.0323506 | 0.00670661 |
| INTS6 | 1.4971815 | 18.6231068 | 1.10E-07 |
| UTS2R | 1.49359152 | 17.000852 | 1.22E-08 |
| TMCC3 | 1.49231667 | 16.8271889 | 6.68E-09 |
| TTC17 | 1.48787732 | 20.8661743 | 1.35E-09 |
| ABCA7 | 1.48608576 | 17.4583992 | 3.35E-05 |
| TASOR | 1.47563424 | 16.5185233 | 1.60E-08 |
| IL18R1 | 1.47311962 | 17.8827832 | 1.98E-08 |
| CNNM2 | 1.47000696 | 20.0531858 | 5.43E-09 |
| STT3B | 1.4695978 | 21.0152972 | 6.97E-10 |
| ADAMTS1 | 1.46830992 | 19.9090441 | 2.07E-09 |
| LOXL2 | 1.46515104 | 17.6790859 | 3.36E-08 |
| POMT1 | 1.46490077 | 19.053088 | 3.52E-08 |
| ENTPD7 | 1.46459891 | 18.4611139 | 5.33E-07 |
| SRPX | 1.46331909 | 18.6865087 | 7.20E-08 |
| ROR2 | 1.45825759 | 17.4491231 | 1.69E-07 |
| TM9SF2 | 1.45478892 | 19.0566891 | 5.25E-04 |

| | | | |
|---|---|---|---|
| RFT1 | 1.45454568 | 19.8471725 | 1.99E-09 |
| KANK2 | 1.45175504 | 17.4469556 | 3.09E-08 |
| MEF2C | 1.44911351 | 17.8412492 | 0.18687435 |
| LEPR | 1.44871061 | 17.2476781 | 3.89E-09 |
| TMEM132E | 1.44624848 | 18.5906819 | 2.80E-08 |
| POMT2 | 1.44135517 | 18.8319146 | 8.68E-08 |
| ARID5B | 1.44116945 | 17.443427 | 1.36E-08 |
| TP53I13 | 1.4394435 | 17.9970855 | 1.73E-06 |
| FKBP10 | 1.438852 | 18.1772562 | 6.25E-08 |
| LTBP3 | 1.43568878 | 17.3981505 | 1.60E-08 |
| EIF4E2 | 1.43414402 | 17.4410852 | 6.87E-08 |
| EGFL7 | 1.43281553 | 20.081029 | 8.08E-07 |
| BMPR1B | 1.43233443 | 15.9459741 | 3.36E-08 |
| TPCN1 | 1.43083059 | 17.4399808 | 1.28E-08 |
| SIDT2 | 1.42613635 | 17.574302 | 2.92E-05 |
| EPHB3 | 1.42167222 | 17.436928 | 4.43E-08 |
| XPR1 | 1.41910531 | 17.9342916 | 2.66E-09 |
| TRIM13 | 1.40899577 | 17.4327025 | 4.17E-08 |
| MINPP1 | 1.40204912 | 17.9714918 | 3.36E-08 |
| GCNT1 | 1.40100108 | 18.3368441 | 1.97E-08 |
| PCSK5 | 1.39335847 | 17.8049403 | 1.37E-06 |
| TMED1 | 1.39152716 | 20.2078287 | 3.04E-05 |
| CHST14 | 1.38898829 | 20.9596751 | 4.77E-05 |

| | | | |
|---|---|---|---|
| ABCB1 | 1.38715233 | 17.4254213 | 4.32E-08 |
| CLTCL1 | 1.38710266 | 17.4254048 | 1.55E-07 |
| TFRC | 1.38467706 | 20.85006 | 1.81E-05 |
| GLA | 1.38109172 | 19.4071204 | 2.90E-09 |
| TPP2 | 1.38071956 | 19.7580742 | 0.00634197 |
| PWP2 | 1.37775695 | 16.5511491 | 1.85E-08 |
| SLC44A2 | 1.37635601 | 17.4218226 | 4.97E-08 |
| PIGN | 1.36905754 | 17.3276002 | 4.84E-06 |
| P4HB | 1.36578112 | 21.518748 | 8.23E-07 |
| PCYOX1 | 1.36525707 | 18.221794 | 2.72E-07 |
| PDIA5 | 1.36376495 | 20.7241788 | 6.57E-05 |
| NOMO2;NOMO3 | 1.36225912 | 18.3620555 | 2.20E-05 |
| LARGE2 | 1.35888724 | 17.0442117 | 2.40E-04 |
| GLB1L2 | 1.35710389 | 19.7713729 | 4.00E-07 |
| PPT1 | 1.35541918 | 17.4148436 | 5.44E-08 |
| TMEM39B | 1.35376376 | 19.6466469 | 1.90E-09 |
| APLP2 | 1.35236535 | 19.3220932 | 1.72E-06 |
| ALKBH2 | 1.35198963 | 16.5597382 | 2.16E-08 |
| P4HA1 | 1.34776086 | 19.9187517 | 5.70E-08 |
| CBX3 | 1.34499066 | 17.4113675 | 1.81E-06 |
| TMEM41A | 1.34335407 | 17.4108219 | 4.13E-07 |
| ESRRB | 1.33384171 | 17.1024639 | 3.78E-07 |
| COL13A1 | 1.33208815 | 17.4070666 | 3.64E-08 |

| | | | |
|---|---|---|---|
| SH2B1 | 1.32353776 | 18.8492515 | 1.82E-07 |
| COLGALT1 | 1.32283249 | 19.3954908 | 2.51E-07 |
| SEMA4F | 1.32053841 | 17.4077265 | 1.69E-07 |
| IGFBP2 | 1.31910488 | 17.4027389 | 2.30E-05 |
| DNAJC3 | 1.31343384 | 21.1644972 | 4.63E-09 |
| MEGF8 | 1.31146879 | 17.5513622 | 7.73E-05 |
| MDK | 1.30799285 | 18.7165028 | 2.60E-07 |
| B2M | 1.30360275 | 17.7638736 | 1.25E-07 |
| FUCA2 | 1.29415471 | 20.1537861 | 1.90E-08 |
| GALNT12 | 1.29315989 | 18.6221052 | 3.50E-05 |
| TMEM167A | 1.29100818 | 16.5800654 | 3.60E-09 |
| TGFB1 | 1.29073907 | 19.445508 | 5.76E-09 |
| TM9SF4 | 1.28704667 | 20.0154912 | 7.20E-08 |
| LMAN2L | 1.28588831 | 19.0666311 | 2.50E-08 |
| SELENOF | 1.28511237 | 16.5311897 | 1.52E-06 |
| PLD3 | 1.28291729 | 19.7127789 | 3.94E-09 |
| FNDC10 | 1.28181903 | 16.6080393 | 1.30E-07 |
| SCARB1 | 1.28100083 | 19.5923958 | 3.00E-08 |
| TMEM59 | 1.27820238 | 16.8682339 | 3.08E-05 |
| SEC11C | 1.27635396 | 18.5761548 | 2.12E-04 |
| SPCS3 | 1.27612856 | 20.5235749 | 2.06E-07 |
| ATF6 | 1.27377858 | 17.8703516 | 6.14E-04 |
| SDK2 | 1.27369376 | 16.7752451 | 1.24E-07 |

| | | | |
|---|---|---|---|
| PGAP4 | 1.27278201 | 19.2578838 | 0.00340088 |
| ESS2 | 1.27249654 | 18.605421 | 8.90E-08 |
| UST | 1.2692494 | 18.2305328 | 7.50E-08 |
| HNRNPM | 1.26639529 | 20.6678807 | 8.50E-08 |
| GORASP2 | 1.26472648 | 17.679067 | 8.09E-09 |
| OSBPL8 | 1.26285189 | 20.2618841 | 1.73E-06 |
| ATF6B | 1.26239617 | 17.6490459 | 2.05E-07 |
| GANAB | 1.26212818 | 22.2577107 | 0.0076021 |
| FAM234B | 1.25857274 | 17.1544704 | 1.18E-06 |
| TM2D3 | 1.25846671 | 17.3825261 | 7.04E-07 |
| GATB | 1.25638183 | 16.5916075 | 1.03E-08 |
| BTN2A1 | 1.25527777 | 18.1289466 | 3.98E-05 |
| NOP9 | 1.25523365 | 18.4489263 | 2.28E-05 |
| LIMD2 | 1.25081442 | 17.3398644 | 1.43E-07 |
| PRRC1 | 1.24828152 | 16.4133642 | 1.42E-07 |
| POGLUT2 | 1.24765436 | 19.22529 | 8.60E-06 |
| PPIC | 1.24682911 | 18.718275 | 1.13E-04 |
| EDEM3 | 1.24060943 | 19.0353872 | 5.01E-07 |
| IGF2R | 1.23922389 | 20.628269 | 4.20E-09 |
| KIDINS220 | 1.23733634 | 18.3884837 | 1.80E-08 |
| B4GALT4 | 1.23311554 | 19.6429234 | 1.87E-05 |
| SFRP1 | 1.23250122 | 18.0878404 | 2.37E-04 |
| RXRB | 1.22960607 | 17.4975292 | 2.14E-05 |

| | | | |
|---|---|---|---|
| GALNT6 | 1.22854516 | 18.1240417 | 4.84E-06 |
| PIGM | 1.22369204 | 19.439956 | 1.25E-07 |
| NRP1 | 1.22318996 | 18.0573118 | 8.81E-05 |
| GNL3L | 1.22274037 | 19.7499579 | 0.03045543 |
| GALNT14 | 1.21855842 | 19.8700807 | 2.91E-08 |
| TCEAL8 | 1.21841873 | 17.577373 | 0.00331375 |
| LRIG1 | 1.21008235 | 18.1148796 | 1.72E-06 |
| PNN | 1.20976579 | 18.4355919 | 1.10E-04 |
| VKORC1L1 | 1.20944636 | 17.8585605 | 6.97E-05 |
| AKIP1 | 1.20149807 | 16.6154672 | 3.95E-06 |
| TMUB1 | 1.20103376 | 18.0344056 | 6.75E-08 |
| ERMP1 | 1.19698395 | 19.3115747 | 7.17E-06 |
| TMX2 | 1.19521435 | 18.4620858 | 4.99E-06 |
| P3H3 | 1.1944633 | 19.9325287 | 2.76E-08 |
| DSE | 1.19327486 | 19.5407206 | 1.41E-06 |
| RNF187 | 1.19306671 | 17.813985 | 1.33E-06 |
| NXPE3 | 1.19249582 | 18.2738813 | 1.31E-05 |
| FLOT2 | 1.1880805 | 16.6143746 | 7.49E-07 |
| UBL5 | 1.18780551 | 17.8321022 | 2.13E-04 |
| FUT8 | 1.18746801 | 18.7425588 | 5.21E-05 |
| EXTL3 | 1.18501487 | 18.7021697 | 8.17E-09 |
| GLCE | 1.18361243 | 17.4066971 | 2.58E-08 |
| ABCD4 | 1.18211607 | 16.3973712 | 4.44E-05 |

| | | | |
|---|---|---|---|
| ABCA3 | 1.17904381 | 19.1666953 | 1.91E-07 |
| TCEAL4 | 1.17493303 | 19.9601876 | 2.80E-08 |
| LGMN | 1.17316331 | 17.3540917 | 1.73E-06 |
| ST6GALNAC3 | 1.16854408 | 18.125949 | 2.37E-08 |
| GALNT4 | 1.16148041 | 18.6822959 | 5.56E-08 |
| TMEM161A | 1.16131285 | 21.0314789 | 4.46E-05 |
| GP1BB | 1.16115719 | 18.8974849 | 1.87E-08 |
| MAN2A2 | 1.16017106 | 19.1119291 | 8.51E-07 |
| GALNT10 | 1.15504333 | 18.3781813 | 1.93E-05 |
| PIGB | 1.15492347 | 18.9919756 | 1.38E-07 |
| YIF1B | 1.15263953 | 20.3017564 | 8.13E-08 |
| CHPF2 | 1.15102276 | 20.2952952 | 2.19E-07 |
| PCSK6 | 1.14753976 | 20.3663489 | 3.32E-07 |
| ST3GAL5 | 1.14005613 | 19.1396462 | 0.00135954 |
| ODAD2 | 1.1393262 | 17.4390067 | 2.85E-06 |
| TOR1AIP1 | 1.13817866 | 19.2518537 | 1.23E-08 |
| B4GALT2 | 1.1354006 | 19.6354975 | 2.17E-06 |
| DNAJC18 | 1.13338848 | 17.8790584 | 1.04E-06 |
| GALNT16 | 1.1292336 | 18.6863023 | 7.04E-04 |
| ILDR2 | 1.12774343 | 17.3389517 | 0.00559373 |
| SDF2L1 | 1.12729366 | 21.088035 | 2.57E-05 |
| SCNM1 | 1.12242659 | 18.257964 | 3.99E-08 |
| MAPK3 | 1.12099363 | 18.3029318 | 5.96E-06 |

| | | | |
|---|---|---|---|
| GPC2 | 1.12000498 | 17.3363722 | 3.95E-04 |
| TELO2 | 1.11965082 | 20.486466 | 2.61E-04 |
| B4GALT1 | 1.11574125 | 18.3888077 | 2.32E-05 |
| BCL2L11 | 1.11521856 | 17.339071 | 1.38E-04 |
| GOLIM4 | 1.11514385 | 18.7874043 | 1.96E-07 |
| YIPF4 | 1.11453792 | 19.3869654 | 3.51E-07 |
| TSPYL1 | 1.11420877 | 21.3272608 | 2.46E-04 |
| DISP1 | 1.10795538 | 17.3323557 | 6.76E-07 |
| ZNHIT1 | 1.10759348 | 17.3322351 | 8.78E-07 |
| CRNN | 1.10707547 | 17.6893183 | 0.00456465 |
| ALG11 | 1.1054655 | 17.3816014 | 3.87E-06 |
| ZNF589 | 1.10200422 | 16.6671858 | 1.38E-06 |
| TIMP3 | 1.09922253 | 20.0420622 | 3.21E-07 |
| CACHD1 | 1.09542864 | 19.2678558 | 6.78E-06 |
| M6PR | 1.09501534 | 17.3280423 | 7.04E-06 |
| TRIM14 | 1.09433376 | 16.4157911 | 0.00416255 |
| LMAN2 | 1.09388431 | 18.986506 | 4.82E-08 |
| SPCS1 | 1.09063864 | 17.2523275 | 7.35E-05 |
| PIGW | 1.09062607 | 19.858851 | 4.31E-07 |
| PKD2 | 1.08275976 | 17.3239572 | 8.56E-07 |
| DIPK1B | 1.08189483 | 19.2688614 | 2.25E-07 |
| EPHB4 | 1.08159308 | 17.9826298 | 1.53E-06 |
| DGAT1 | 1.07990797 | 16.9985011 | 1.99E-08 |

| | | | |
|---|---|---|---|
| RAVER1 | 1.07514862 | 17.7614365 | 2.18E-05 |
| ASPH | 1.07370114 | 20.0260705 | 3.75E-07 |
| GIGYF2 | 1.07214043 | 16.5175291 | 0.06053019 |
| ATRN | 1.07156266 | 18.2708283 | 4.44E-07 |
| GOLGA5 | 1.07000174 | 15.8916596 | 9.60E-08 |
| NECTIN3 | 1.06979084 | 17.9805646 | 2.91E-06 |
| GALNT3 | 1.0692471 | 18.2034705 | 0.0018366 |
| NETO2 | 1.06552834 | 17.9570038 | 2.33E-05 |
| SPATA7 | 1.06372463 | 17.7343931 | 3.76E-07 |
| PLXNB2 | 1.06050965 | 17.2974555 | 2.40E-07 |
| UBE2I | 1.05920119 | 16.775706 | 3.20E-04 |
| MARCHF7 | 1.05698672 | 18.9701571 | 1.09E-07 |
| CYP27C1 | 1.05442204 | 17.7601336 | 9.56E-06 |
| OSTC | 1.05347507 | 19.8207137 | 2.36E-04 |
| LGALS1 | 1.05266673 | 17.3139261 | 7.14E-07 |
| LMLN | 1.05198923 | 17.3137003 | 4.87E-05 |
| NUP153 | 1.05190004 | 18.1783224 | 1.36E-07 |
| CHSY3 | 1.05013245 | 18.7529002 | 2.06E-04 |
| CELSR1 | 1.04817988 | 18.848828 | 1.89E-07 |
| GALNT7 | 1.0478137 | 19.9499282 | 1.72E-06 |
| LMF2 | 1.04540621 | 21.6119357 | 0.0160942 |
| SELENOS | 1.04324917 | 16.4562872 | 7.00E-08 |
| SCAMP4 | 1.0410829 | 19.8096506 | 1.62E-07 |

| | | | |
|---|---|---|---|
| APH1B | 1.04006306 | 16.0912426 | 5.56E-06 |
| B4GALNT1 | 1.03885497 | 16.5458049 | 1.43E-04 |
| ACP1 | 1.03095856 | 19.1171006 | 4.07E-05 |
| ANTXR1 | 1.03022328 | 18.2145639 | 9.09E-06 |
| METTL25B | 1.02890686 | 16.1031079 | 1.12E-07 |
| ADPGK | 1.0264003 | 18.3892514 | 2.07E-05 |
| SF3A2 | 1.02031664 | 17.7216584 | 0.01481717 |
| CPD | 1.01999574 | 19.5879482 | 1.02E-07 |
| DYM | 1.01772498 | 18.5298232 | 2.02E-07 |
| TXNDC16 | 1.01592518 | 18.3115861 | 2.84E-07 |
| VDAC1 | 1.01337272 | 21.7530093 | 0.00945036 |
| FAM133A;FAM133B | 1.01235598 | 21.7346322 | 0.0233101 |
| SBDS | 1.0087912 | 19.7966407 | 6.92E-06 |
| XPO4 | 1.00812615 | 18.7007941 | 3.40E-04 |
| MELK | 1.00655437 | 17.2652304 | 2.09E-05 |
| DIPK1A | 1.00579093 | 17.2719037 | 1.37E-05 |
| EMC2 | 1.00417901 | 18.5908604 | 2.34E-06 |
| IGFBPL1 | 1.00149911 | 17.2968703 | 6.78E-05 |

**Table S5. Secretory pathway markers.** List of secretory pathway antibodies used throughout the manuscript in immunofluorescence experiments. CST, Cell Signaling Technology.

| Organelle | Marker (host) | Used antibodies, dilution |
|---|---|---|
| Endoplasmic reticulum (ER) | Calnexin (rabbit)<br>Calnexin (mouse)<br>Calnexin (goat) | Abcam ab22595, 1:5000<br>Santa Cruz sc-46669, 1:100<br>SICGEN AB0037-200, 1:1500 |
| COPII vesicles | SEC31A (rabbit)<br>SEC31A (mouse) | CST 13466S, 1:100<br>Santa Cruz sc-136233, 1:100 |
| COPI vesicles | COPB2 (rabbit)<br>COPB2 (mouse) | Invitrogen PA5-96557, 1:5000<br>Santa Cruz sc-393615, 1:200 |
| cis-Golgi | GM130 (rabbit)<br>GM130 (mouse)<br>GRASP55 (rabbit) | CST 12480S, 1:500<br>Abcam, ab169276, 1:1500<br>Proteintech 10598-1-AP, 1:1000 |
| Early endosome | EEA1 (rabbit)<br>EEA1 (mouse) | CST 3288S, 1:100<br>CST 48453S, 1:100 |
| Late endosome | RAB7 (rabbit)<br>RAB7 (mouse) | CST 9367S, 1:100<br>CST 95746S, 1:100 |
| Lysosome | LAMP1 (rabbit)<br>LAMP1 (mouse)<br>LAMP2 (goat) | CST 9091S, 1:100<br>CST 15665S, 1:100<br>R&D AF6228, 1:400 |

**Table S6. IP-MS of TMED7-MYC and TMED7 with TMED5CC-MYC**

Proteins that preferentially bind TMED7 compared to TMED7 with a TMED5 CC domain.

Proteins (with gene names) are sorted with logFC value and were analyzed using the Limma package in R.

| Gene | logFC | Average Expression | Adj. p value |
|---|---|---|---|
| S100A8 | 9.13171503 | 21.72117 | 4.11E-17 |
| KRT82 | 7.44628915 | 19.6662457 | 2.43E-19 |
| ATP6V1F | 7.07738052 | 18.6549337 | 8.02E-16 |
| BPNT2 | 6.2806535 | 21.7954247 | 0.00238063 |
| IGKV3-15;IGKV3-7;IGKV3D-7 | 5.85724652 | 21.8849337 | 3.77E-15 |
| UNC5C | 5.59696536 | 20.4842958 | 8.95E-17 |
| CLTB | 5.11979263 | 19.2697212 | 7.87E-13 |
| TTLL2 | 5.06804365 | 19.1905966 | 1.67E-15 |
| BLZF1 | 4.94716913 | 20.3047035 | 9.22E-12 |
| CLTA | 4.84549306 | 20.8421005 | 2.14E-10 |
| BRI3BP | 4.51079479 | 20.8171272 | 3.17E-13 |
| KRT31 | 4.33734445 | 19.9822654 | 7.74E-07 |
| TBC1D30 | 4.23912721 | 19.9384508 | 7.83E-13 |
| CLMN | 4.16336739 | 19.1435568 | 4.38E-06 |
| RNF26 | 3.99450604 | 19.5079014 | 2.51E-10 |
| ZZZ3 | 3.86372575 | 16.0835424 | 3.56E-14 |
| ZDHHC9 | 3.84913207 | 20.7888007 | 7.70E-12 |
| PUS3 | 3.6574373 | 18.1992376 | 1.08E-05 |

| | | | |
|---|---|---|---|
| KLHL22 | 3.6317254 | 20.2233949 | 9.44E-13 |
| URB1 | 3.48878464 | 19.0112791 | 6.32E-04 |
| IAH1 | 3.40812129 | 16.1403568 | 1.32E-12 |
| BBS12 | 3.3870644 | 16.6621554 | 1.89E-13 |
| CLINT1 | 3.38704487 | 22.3071589 | 5.23E-13 |
| SMARCAL1 | 3.38374025 | 16.3053336 | 7.51E-13 |
| ZNF770 | 3.334886 | 18.6725359 | 8.03E-14 |
| MIF | 3.29779317 | 19.7139742 | 1.42E-08 |
| SLC35F5 | 3.28083262 | 20.27653 | 1.74E-09 |
| CTIF | 3.2779947 | 18.8325868 | 3.02E-14 |
| ZNF462 | 3.26561243 | 16.2693521 | 1.73E-12 |
| KRT33B | 3.23369304 | 19.3700793 | 6.59E-11 |
| SLC39A1 | 3.22480184 | 19.7209804 | 1.88E-08 |
| GRIN2D | 3.21571287 | 18.8201305 | 3.13E-14 |
| ANO6 | 3.19759927 | 18.8165078 | 1.65E-13 |
| TMC7 | 3.1898141 | 17.692606 | 0.00146695 |
| GORASP2 | 3.11595196 | 21.5483795 | 2.06E-11 |
| SLC7A3 | 3.11262495 | 21.4378546 | 6.57E-08 |
| SARAF | 3.08278025 | 19.4190099 | 9.90E-11 |
| RANBP9 | 3.05946415 | 18.5006876 | 0.00515282 |
| CLTC | 2.98100454 | 25.1459889 | 1.20E-13 |
| CC2D1B | 2.97520657 | 20.0991583 | 2.27E-08 |
| KRT86 | 2.97067601 | 20.1441005 | 1.20E-13 |

| | | | |
|---|---|---|---|
| THOC1 | 2.91994603 | 17.0125095 | 4.13E-13 |
| PRICKLE1 | 2.91818842 | 19.0297376 | 2.66E-10 |
| VAMP2;VAMP3 | 2.90364572 | 20.2629268 | 5.16E-11 |
| UBE2J1 | 2.8753765 | 22.8790157 | 1.01E-05 |
| SEC11C | 2.85772147 | 19.761293 | 4.75E-11 |
| CLTCL1 | 2.84951055 | 18.74689 | 2.55E-12 |
| PSEN2 | 2.83479649 | 18.5725944 | 1.29E-10 |
| UBR4 | 2.82387818 | 23.2095739 | 4.47E-15 |
| COL1A2 | 2.81723447 | 20.1538516 | 2.54E-14 |
| VCPIP1 | 2.79585878 | 18.7200566 | 0.02188118 |
| FZD7 | 2.74288219 | 18.7255643 | 1.58E-11 |
| CCDC66 | 2.72488649 | 16.5160336 | 2.00E-12 |
| TSPAN10 | 2.69624304 | 17.9996148 | 2.71E-09 |
| PGAP3 | 2.68141911 | 17.0461915 | 7.01E-10 |
| NTAQ1 | 2.67365356 | 18.7117186 | 5.23E-13 |
| PCNX1 | 2.63937513 | 19.2145771 | 3.17E-13 |
| ZFAND6 | 2.59122959 | 16.4684818 | 4.69E-11 |
| COG2 | 2.56325066 | 20.3099268 | 0.04451345 |
| LAMTOR2 | 2.52453569 | 19.067707 | 1.25E-08 |
| EFNA5 | 2.5232001 | 19.9648802 | 8.86E-09 |
| CCNL2 | 2.51559095 | 16.8612289 | 2.66E-08 |
| DHCR7 | 2.51305233 | 25.1167396 | 6.26E-04 |
| PATL1 | 2.48540823 | 16.3499895 | 2.81E-10 |

| | | | |
|---|---|---|---|
| HGSNAT | 2.47155008 | 17.9629589 | 3.12E-11 |
| RPL28 | 2.46850379 | 23.8600095 | 1.34E-04 |
| IDH2 | 2.46141746 | 23.5671271 | 0.02006605 |
| TEDC1 | 2.45488036 | 20.5936584 | 0.03061295 |
| VSTM4 | 2.42247424 | 18.5728338 | 2.82E-11 |
| KRT85 | 2.42167281 | 19.9195888 | 7.32E-05 |
| RHBDD3 | 2.40958769 | 19.2341868 | 2.96E-10 |
| ZNF282 | 2.40747753 | 16.3108859 | 3.82E-11 |
| SLC6A8 | 2.40259069 | 18.8380004 | 6.24E-13 |
| SH2B1 | 2.39010707 | 21.6418456 | 4.49E-10 |
| KRTAP11-1 | 2.38664101 | 19.2602389 | 3.22E-12 |
| OARD1 | 2.38171409 | 16.2152244 | 1.37E-11 |
| SLC52A2 | 2.35518307 | 17.6247577 | 1.00E-06 |
| RAB32 | 2.3411243 | 22.4573214 | 2.85E-04 |
| WDR33 | 2.3372144 | 18.7546216 | 0.00382353 |
| SFXN4 | 2.33215807 | 24.6307525 | 0.00219051 |
| ZC3H8 | 2.32019747 | 16.0259417 | 2.25E-07 |
| PHF3 | 2.31861501 | 16.9245193 | 1.24E-08 |
| HMGCS1 | 2.31533794 | 17.1265696 | 4.53E-10 |
| NBEAL2 | 2.31320221 | 18.3569584 | 0.01166043 |
| TMUB1 | 2.30744076 | 20.0461118 | 0.0076691 |
| RRP1 | 2.29646627 | 19.4658027 | 0.002793 |
| TMED3 | 2.2848021 | 23.5636362 | 1.91E-06 |

| | | | |
|---|---|---|---|
| RAB36 | 2.27496747 | 19.583969 | 1.97E-10 |
| STX8 | 2.26963661 | 19.4758161 | 5.33E-12 |
| SGPL1 | 2.26187232 | 22.9397435 | 0.04353337 |
| ATP9B | 2.2582008 | 19.6598754 | 9.20E-13 |
| IKZF1 | 2.25754743 | 16.3267357 | 2.79E-07 |
| PPP2R5E | 2.25191488 | 19.3599702 | 4.55E-07 |
| KRT36 | 2.25103242 | 20.4441319 | 4.46E-11 |
| SLC43A1 | 2.24876552 | 19.4878625 | 4.99E-10 |
| P3H2 | 2.23936088 | 19.7164414 | 6.48E-08 |
| TSPAN12 | 2.21745332 | 18.5621236 | 1.76E-10 |
| EFTUD2 | 2.21714262 | 22.1901757 | 0.01494192 |
| UBL4A | 2.20436189 | 19.3815417 | 1.64E-05 |
| VPS4B | 2.19922523 | 20.5580386 | 9.13E-12 |
| ERCC6L | 2.19283403 | 17.2271973 | 1.89E-11 |
| NDRG3 | 2.17881964 | 19.0356369 | 5.11E-05 |
| DHX29 | 2.17770592 | 23.6356467 | 0.02692623 |
| BSCL2 | 2.17700291 | 21.6770515 | 0.00828717 |
| SLC4A2 | 2.16897811 | 20.8591752 | 3.80E-11 |
| ATP11C | 2.149915 | 21.4724219 | 1.39E-10 |
| NEIL1 | 2.14064384 | 16.6958739 | 3.24E-08 |
| GLYCTK | 2.13830184 | 19.2323595 | 2.81E-10 |
| FBXW9 | 2.13684431 | 19.4454606 | 3.87E-08 |
| MOSPD2 | 2.1362744 | 20.272768 | 9.47E-10 |

| | | | |
|---|---|---|---|
| PLPP5 | 2.13079468 | 19.3952046 | 3.18E-11 |
| KCMF1 | 2.11900658 | 21.3213822 | 6.90E-08 |
| PORCN | 2.11685886 | 20.2353962 | 9.58E-08 |
| HDDC3 | 2.1148625 | 19.2241533 | 2.43E-11 |
| PRAME | 2.10216451 | 21.4642756 | 0.00271054 |
| DCXR | 2.09443369 | 19.3441914 | 6.22E-07 |
| SLC47A1 | 2.07617795 | 20.1472868 | 8.74E-12 |
| PEAK1 | 2.07215811 | 20.3222433 | 3.82E-11 |
| GASK1A | 2.04591448 | 19.087301 | 6.49E-08 |
| MEAF6 | 2.04489372 | 17.3069341 | 3.00E-09 |
| LEPROTL1 | 2.04263025 | 20.2874846 | 9.29E-09 |
| DOP1B | 2.03982277 | 19.4255469 | 0.03507099 |
| MIGA2 | 2.03777027 | 17.3959248 | 1.41E-11 |
| SLC30A9 | 2.03604734 | 22.2715156 | 1.56E-10 |
| ALX3;ALX4 | 2.03352238 | 18.9951353 | 3.56E-08 |
| YIPF6 | 2.02220495 | 22.6869448 | 7.85E-09 |
| TNRC6B | 2.02114984 | 18.2491655 | 2.80E-05 |
| SLC6A15 | 2.01825506 | 19.7375095 | 1.21E-07 |
| CHPT1 | 2.01658939 | 19.0394526 | 4.13E-10 |
| PEMT | 2.01559145 | 19.7431584 | 1.44E-10 |
| NUDT8 | 2.01380105 | 20.0091104 | 1.58E-07 |
| SLC33A1 | 2.00668226 | 19.339894 | 1.64E-04 |
| BBS1 | 2.00436864 | 19.6836548 | 1.26E-04 |

| | | | |
|---|---|---|---|
| FASTKD3 | 1.99290485 | 20.7090046 | 3.21E-06 |
| MED4 | 1.98298944 | 16.9705414 | 1.06E-10 |
| FIS1 | 1.97838964 | 19.7532583 | 1.06E-07 |
| SLC25A46 | 1.97767117 | 20.2937527 | 3.43E-07 |
| KRTCAP2 | 1.97764154 | 19.2486612 | 2.61E-10 |
| DAGLB | 1.97734896 | 19.6082902 | 6.83E-04 |
| LYPD6B | 1.96567766 | 18.5701234 | 7.63E-11 |
| CD151 | 1.96500511 | 20.5298928 | 4.91E-10 |
| SLC38A9 | 1.96381935 | 17.7739521 | 9.00E-10 |
| SETD9 | 1.95236129 | 19.7838594 | 1.60E-10 |
| SCAMP4 | 1.95225778 | 23.8735353 | 1.48E-06 |
| THBS1 | 1.94846771 | 17.7767942 | 7.39E-07 |
| SLC38A1 | 1.94676654 | 18.9288087 | 1.11E-08 |
| C1GALT1 | 1.93996968 | 20.8847941 | 0.0364733 |
| SLC12A4 | 1.93521577 | 18.3462682 | 1.88E-08 |
| TMEM115 | 1.93269158 | 22.7073898 | 0.00571273 |
| FAM83D | 1.93141048 | 23.4431914 | 0.27134969 |
| ATP1B3 | 1.92610401 | 22.9126648 | 1.56E-09 |
| PHB1 | 1.91642576 | 22.7962476 | 6.55E-06 |
| TUBB2B | 1.91545404 | 19.442103 | 0.039158 |
| SMAP2 | 1.90154576 | 19.0708454 | 2.81E-10 |
| SYNGR1 | 1.90152547 | 20.3683339 | 8.81E-08 |
| MSRB3 | 1.88102415 | 22.0567089 | 8.66E-07 |

| | | | |
|---|---|---|---|
| ACADSB | 1.87689695 | 22.8157638 | 0.00141663 |
| SLC1A3 | 1.87400339 | 21.9226543 | 2.26E-11 |
| SLC7A5 | 1.87357015 | 18.7164645 | 1.18E-10 |
| ABCG1 | 1.87116974 | 16.9598985 | 8.59E-07 |
| CTU1 | 1.86627538 | 17.8582256 | 0.00637793 |
| PKD2 | 1.8605938 | 18.5028252 | 1.14E-07 |
| TADA2B | 1.85718214 | 18.3523368 | 2.49E-07 |
| BTAF1 | 1.85050231 | 23.7484244 | 3.28E-09 |
| HNRNPH3 | 1.83829161 | 17.7245334 | 3.65E-10 |
| GOLGA2 | 1.82310634 | 22.2004089 | 0.00182219 |
| FUNDC2 | 1.80957309 | 19.4126065 | 9.90E-10 |
| ATP11A | 1.80614945 | 18.5382178 | 5.72E-10 |
| NFE2L1 | 1.80290146 | 18.5375682 | 2.36E-10 |
| TMEM147 | 1.79949873 | 21.9439071 | 7.34E-06 |
| LRRC8D | 1.79381938 | 19.4326077 | 1.37E-05 |
| LETMD1 | 1.79012034 | 20.0201575 | 7.74E-07 |
| GAK | 1.77575074 | 20.5738197 | 9.24E-10 |
| SLC23A2 | 1.77498203 | 18.3952989 | 1.10E-07 |
| STX6 | 1.77391021 | 19.8819847 | 2.04E-10 |
| RCE1 | 1.7605815 | 19.7578347 | 2.28E-09 |
| DEGS1 | 1.7600105 | 19.2465093 | 2.73E-07 |
| CDH24 | 1.75807336 | 21.291565 | 7.80E-05 |
| SLC25A25 | 1.75238793 | 20.225719 | 1.88E-06 |

| | | | |
|---|---|---|---|
| GJC1 | 1.7485889 | 18.5992926 | 3.56E-08 |
| SVEP1 | 1.74669515 | 18.1702006 | 5.23E-09 |
| CCDC167 | 1.74508094 | 20.4217382 | 6.25E-07 |
| SLC39A3 | 1.73729466 | 19.8263467 | 8.46E-05 |
| SLCO4A1 | 1.73707493 | 17.8299409 | 1.91E-08 |
| IFRD1 | 1.72351218 | 20.3066119 | 0.14318755 |
| SLC18B1 | 1.72252189 | 18.7114248 | 2.53E-07 |
| CYP20A1 | 1.72004483 | 17.7653578 | 2.66E-10 |
| FZD2 | 1.71984923 | 22.8498727 | 1.96E-09 |
| ATP2B1 | 1.71960618 | 22.1894313 | 1.11E-09 |
| RFT1 | 1.71878145 | 21.3994899 | 7.65E-08 |
| AKNA | 1.71852442 | 19.0311875 | 4.22E-08 |
| TRMT2B | 1.71762338 | 19.2194769 | 5.32E-10 |
| PIK3C2A | 1.71459186 | 21.271451 | 2.87E-09 |
| XPO4 | 1.71141762 | 21.8336628 | 2.09E-04 |
| NSMCE4A | 1.70686734 | 19.5331372 | 1.92E-06 |
| B4GALT4 | 1.70444683 | 22.33092 | 0.01261192 |
| SLC6A11 | 1.7005201 | 18.1079096 | 1.29E-10 |
| ERMP1 | 1.69294874 | 20.8585044 | 0.06379981 |
| GHITM | 1.69052769 | 21.1581522 | 1.37E-06 |
| CSTA | 1.68246002 | 21.1264446 | 0.00834502 |
| TMEM63A | 1.67985527 | 17.1535211 | 6.87E-06 |
| TRIM27 | 1.67115632 | 19.3547139 | 4.50E-07 |

| | | | |
|---|---|---|---|
| YIPF5 | 1.666898 | 22.3206891 | 2.42E-09 |
| CCM2 | 1.66507456 | 15.9384716 | 0.20165961 |
| STT3A | 1.6632095 | 26.2155965 | 3.00E-08 |
| TBC1D16 | 1.66122368 | 16.7557244 | 7.59E-08 |
| FOXP4 | 1.65584293 | 19.0324105 | 0.0155331 |
| KRT35 | 1.63788424 | 18.5045648 | 5.39E-10 |
| MFSD14B | 1.63621833 | 19.3815298 | 6.06E-09 |
| OSTC | 1.63219706 | 24.5717377 | 2.35E-07 |
| SLC44A1 | 1.62529551 | 17.5254418 | 0.00163217 |
| MT-ND1 | 1.62341838 | 19.4164353 | 1.53E-06 |
| NCAPD3 | 1.6205783 | 21.499379 | 0.00591992 |
| HACD3 | 1.61856796 | 27.4738313 | 4.62E-04 |
| THAP4 | 1.61235912 | 19.4966357 | 4.09E-08 |
| TMEM185A | 1.60773216 | 17.4975401 | 3.57E-09 |
| MBOAT1 | 1.60677002 | 20.9411635 | 4.67E-06 |
| NATD1 | 1.59641096 | 17.7729095 | 6.25E-07 |
| YIF1B | 1.5947655 | 23.267613 | 2.74E-08 |
| GEMIN8 | 1.58864866 | 19.3289264 | 1.96E-07 |
| BIRC2 | 1.58765355 | 17.8295484 | 3.66E-07 |
| RRN3 | 1.58656704 | 19.2322835 | 1.28E-05 |
| ENDOD1 | 1.58476279 | 17.715732 | 1.25E-08 |
| C12ORF57 | 1.58195636 | 17.0632633 | 5.05E-09 |
| IL13RA1 | 1.58138593 | 17.4680493 | 8.45E-07 |

| | | | |
|---|---|---|---|
| XPO1 | 1.5803863 | 25.6591337 | 0.0015878 |
| SLC30A5 | 1.57405151 | 20.9993452 | 9.63E-08 |
| JMJD8 | 1.57209908 | 20.8790063 | 2.29E-06 |
| RAP1GAP | 1.5696683 | 18.4909216 | 1.14E-09 |
| PSIP1 | 1.56957375 | 18.9764082 | 1.23E-06 |
| PPFIBP1 | 1.5670822 | 21.1771849 | 6.39E-04 |
| ARL16 | 1.56306613 | 18.9604168 | 1.92E-06 |
| BRF2 | 1.5620541 | 19.3867606 | 5.75E-06 |
| ATP11B | 1.56189493 | 17.9913388 | 8.86E-09 |
| CALML3 | 1.55951831 | 18.4888916 | 3.77E-09 |
| CIP2A | 1.55462224 | 22.8103944 | 3.58E-10 |
| CALML5 | 1.554414 | 19.8464482 | 0.11986086 |
| EXOG | 1.55380831 | 18.9708936 | 2.81E-08 |
| SLC35G2 | 1.55366681 | 22.0066503 | 1.80E-07 |
| UBE2K | 1.55171316 | 17.5196726 | 1.25E-05 |
| ABCA7 | 1.5480197 | 18.4865918 | 3.10E-08 |
| UBE2S | 1.54558435 | 16.4058655 | 0.03452031 |
| SLC35A5 | 1.54473532 | 19.506655 | 1.03E-05 |
| XPNPEP1 | 1.544329 | 17.4775161 | 2.02E-10 |
| CDC23 | 1.54305101 | 21.330724 | 8.34E-04 |
| SYF2 | 1.53580339 | 17.3228728 | 6.54E-08 |
| CKLF | 1.53189128 | 19.1470114 | 8.14E-08 |
| TMEM219 | 1.53087245 | 20.0586382 | 3.58E-06 |

| | | | |
|---|---|---|---|
| PCGF1 | 1.52935839 | 17.1858071 | 9.79E-07 |
| DES | 1.52789302 | 17.6852286 | 1.24E-04 |
| B4GALT3 | 1.52314184 | 19.8929052 | 7.18E-09 |
| EIF2A | 1.51769287 | 17.7796934 | 1.61E-09 |
| MTFR2 | 1.51607436 | 19.0873226 | 2.34E-04 |
| PIGA | 1.51349332 | 21.5061712 | 2.98E-05 |
| SCN5A | 1.51332552 | 18.9945441 | 1.97E-07 |
| GTSE1 | 1.51020525 | 22.103844 | 5.16E-07 |
| ZNF239 | 1.50967216 | 20.2073839 | 1.63E-04 |
| SGMS2 | 1.50527891 | 16.9263071 | 8.81E-09 |
| MFSD10 | 1.5046537 | 18.6824028 | 2.43E-06 |
| SERINC1 | 1.50420659 | 20.5598015 | 1.28E-07 |
| LRRC8E | 1.50262121 | 18.6727122 | 1.04E-08 |
| CLCN7 | 1.50246003 | 22.6540578 | 3.70E-06 |
| RNF145 | 1.49868313 | 21.1232588 | 1.56E-04 |
| LEPR | 1.49480483 | 20.1456202 | 0.04908064 |
| TMED9 | 1.49480049 | 26.3011035 | 2.77E-06 |
| CCDC22 | 1.49442226 | 19.1362251 | 0.14475073 |
| SEC61A1 | 1.49394397 | 26.6406205 | 2.34E-08 |
| DAD1 | 1.49351908 | 22.5115998 | 5.00E-06 |
| MON1A | 1.49316518 | 18.2039328 | 3.19E-10 |
| STARD3 | 1.4930413 | 19.0946724 | 6.95E-06 |
| PHB2 | 1.49259554 | 24.6472514 | 3.04E-09 |

| | | | |
|---|---|---|---|
| TMEM186 | 1.48858766 | 21.1869088 | 0.01329346 |
| SLC1A4 | 1.4871836 | 20.0278331 | 8.25E-07 |
| SLC12A7 | 1.48013018 | 17.7490492 | 8.03E-08 |
| GJA1 | 1.47795673 | 22.1584653 | 8.56E-08 |
| UNG | 1.47308636 | 18.9151355 | 4.26E-08 |
| ARSB | 1.46869554 | 20.6175064 | 0.00587342 |
| TIMM23 | 1.46714287 | 23.0240725 | 9.11E-06 |
| RABL3 | 1.46712285 | 21.3882801 | 1.50E-06 |
| SLC22A18 | 1.46611596 | 17.6290065 | 1.80E-08 |
| TIMMDC1 | 1.46568347 | 24.9674108 | 4.46E-05 |
| TTYH3 | 1.46500179 | 16.99203 | 1.54E-06 |
| VDAC1 | 1.45845207 | 24.9094359 | 0.00283054 |
| SPPL2A | 1.4575643 | 20.5379779 | 1.56E-09 |
| ELOVL2 | 1.45563192 | 18.2011575 | 0.00178085 |
| HM13 | 1.45345432 | 21.1140526 | 1.40E-05 |
| ATP6V0A2 | 1.44958435 | 20.4091053 | 9.56E-07 |
| FAM8A1 | 1.44630994 | 21.73232 | 2.32E-10 |
| TMEM69 | 1.44594264 | 17.1418791 | 5.44E-07 |
| LAMTOR3 | 1.44033534 | 20.505036 | 0.00990437 |
| TOLLIP | 1.43826162 | 16.0720237 | 1.45E-07 |
| SERINC5 | 1.43795906 | 18.2677295 | 1.26E-05 |
| MMGT1 | 1.43545775 | 25.8614459 | 0.0041915 |
| SEMG1 | 1.43486309 | 18.4639605 | 2.66E-08 |

| | | | |
|---|---|---|---|
| SLC39A14 | 1.4321082 | 21.1250718 | 0.00316937 |
| FLVCR1 | 1.4308264 | 21.6777136 | 1.15E-08 |
| SLC30A6 | 1.4305459 | 20.7109636 | 7.13E-05 |
| MANBAL | 1.42974566 | 20.0276543 | 1.46E-04 |
| GOLGA5 | 1.42778746 | 18.641411 | 8.53E-07 |
| MFSD1 | 1.42478107 | 19.2099405 | 1.79E-07 |
| PTGES2 | 1.42401358 | 20.5810386 | 0.00270728 |
| MARCHF6 | 1.42249775 | 18.2206839 | 7.04E-06 |
| FADS2 | 1.41798746 | 23.379155 | 3.38E-04 |
| HPX | 1.41745373 | 17.40983 | 7.10E-07 |
| SLC35C2 | 1.41606344 | 20.0554143 | 4.18E-08 |
| BORA | 1.41583891 | 20.7112382 | 1.01E-06 |
| TMEM53 | 1.40932853 | 20.2141332 | 0.01110174 |
| NSFL1C | 1.4080978 | 17.3746736 | 3.14E-07 |
| ALG9 | 1.40781971 | 20.8784971 | 6.37E-06 |
| ST7 | 1.40496288 | 21.6402839 | 0.05467024 |
| DISP2 | 1.40246497 | 17.6636719 | 3.10E-08 |
| FXR2 | 1.40118788 | 18.9684271 | 2.26E-09 |
| MAP6D1 | 1.39656696 | 18.0072134 | 9.33E-09 |
| ATP1A3 | 1.39552459 | 20.7138102 | 7.23E-07 |
| DPAGT1 | 1.39186437 | 19.7187237 | 6.51E-07 |
| CDC34;UBE2R2 | 1.38918986 | 18.6256658 | 1.41E-08 |
| G6PC3 | 1.38824301 | 24.2060455 | 1.04E-06 |

| | | | |
|---|---|---|---|
| ZNF410 | 1.3880285 | 18.1524107 | 2.13E-07 |
| ATP1A1 | 1.38704393 | 25.7908871 | 1.99E-07 |
| ALG8 | 1.38642179 | 22.0400946 | 2.37E-06 |
| SHPK | 1.38585018 | 17.6343165 | 6.43E-09 |
| SLC25A21 | 1.38496985 | 20.9499974 | 2.00E-06 |
| TMEM175 | 1.38483152 | 18.9324102 | 2.93E-04 |
| CEPT1 | 1.37808759 | 22.4170245 | 2.46E-08 |
| METTL3 | 1.3780841 | 20.4077574 | 1.36E-05 |
| EXOSC1 | 1.37670638 | 17.3238903 | 3.44E-07 |
| DDX10 | 1.37550074 | 17.698375 | 8.39E-09 |
| SLC39A8 | 1.37280331 | 20.8909375 | 1.02E-04 |
| PAQR3 | 1.37235617 | 20.6453934 | 2.06E-06 |
| CCNH | 1.37050756 | 18.7422153 | 4.01E-05 |
| ICAM4 | 1.36473109 | 20.4132741 | 5.99E-05 |
| SLC29A1 | 1.36321843 | 19.4543653 | 3.88E-06 |
| ANO5 | 1.36052557 | 20.4637952 | 0.00151224 |
| CMTM4 | 1.35987254 | 20.2873508 | 0.0083408 |
| CABLES1 | 1.35943656 | 21.1711371 | 4.20E-06 |
| SSR4 | 1.35940512 | 26.4834618 | 0.09414576 |
| SLC22A5 | 1.35900601 | 20.5269034 | 3.26E-04 |
| SLC4A7 | 1.35215528 | 17.6502084 | 1.04E-06 |
| ATP6V0A1 | 1.35153919 | 22.0056245 | 2.27E-06 |
| PIGH | 1.34881457 | 21.694277 | 1.11E-06 |

| | | | |
|---|---|---|---|
| GPR180 | 1.3477051 | 22.6512836 | 9.63E-08 |
| NR3C1 | 1.34764165 | 21.3871713 | 8.43E-04 |
| TMEM30A | 1.34666375 | 22.9791367 | 0.0364733 |
| UBIAD1 | 1.34622469 | 19.3821649 | 1.52E-04 |
| TMUB2 | 1.34374451 | 20.6854555 | 1.34E-04 |
| FBXO6 | 1.34308875 | 18.2340371 | 3.37E-07 |
| PDLIM1 | 1.34104866 | 17.5325858 | 2.52E-08 |
| TMED1 | 1.33977323 | 23.3303366 | 2.52E-09 |
| MBTPS2 | 1.339432 | 22.847334 | 2.79E-07 |
| ATP5MC2 | 1.33905272 | 21.5880697 | 0.00412684 |
| ANKRD52 | 1.33892251 | 19.9344084 | 0.01962175 |
| CBX4 | 1.33768842 | 18.6852602 | 3.42E-05 |
| C2CD2L | 1.33726278 | 20.4253963 | 5.87E-07 |
| SLC1A5 | 1.33416072 | 24.2704214 | 7.02E-07 |
| PAK2 | 1.33360655 | 18.8298099 | 6.07E-08 |
| THAP12 | 1.32731418 | 18.5250055 | 6.63E-04 |
| NISCH | 1.32317267 | 20.0588903 | 0.01104024 |
| FZD1 | 1.32205196 | 18.0824802 | 6.89E-05 |
| SLC7A2 | 1.32174465 | 18.3545145 | 1.25E-04 |
| SLC26A2 | 1.32107928 | 18.7170062 | 1.92E-06 |
| PAM | 1.32099334 | 20.6895667 | 2.04E-05 |
| WWOX | 1.32067808 | 18.9811667 | 2.50E-06 |
| ZDHHC3 | 1.32035022 | 18.7196687 | 6.12E-07 |

| | | | |
|---|---|---|---|
| ACAT1 | 1.32030852 | 19.3607835 | 5.88E-04 |
| POGLUT3 | 1.32004656 | 20.6265938 | 0.02739064 |
| KLHL13 | 1.31986641 | 19.6950815 | 4.66E-04 |
| TNRC6A | 1.31789757 | 16.7641362 | 3.54E-04 |
| SDHD | 1.31566552 | 18.5336677 | 1.59E-07 |
| DENND11 | 1.3152734 | 18.5553876 | 5.54E-08 |
| KCNG1 | 1.31203932 | 18.556592 | 3.68E-05 |
| RTL10 | 1.30952844 | 18.5721643 | 1.14E-07 |
| SLC4A11 | 1.30872512 | 18.3627821 | 4.20E-06 |
| NAAA | 1.30855927 | 16.3853452 | 5.59E-06 |
| ACOT8 | 1.30770363 | 23.0935956 | 8.54E-05 |
| TMEM161B | 1.30320633 | 18.4313388 | 4.58E-05 |
| ORC5 | 1.30319684 | 22.1775635 | 6.99E-05 |
| PIGW | 1.30132865 | 22.0489961 | 3.20E-08 |
| PEX12 | 1.30054891 | 17.4909076 | 7.21E-05 |
| FKBP5 | 1.30023467 | 18.8605629 | 2.54E-09 |
| SLC30A1 | 1.29924845 | 18.3881755 | 3.55E-08 |
| TMPPE | 1.29763889 | 18.8734499 | 0.00379213 |
| FAM210A | 1.29694576 | 21.0688029 | 8.99E-08 |
| ZFYVE27 | 1.29367798 | 16.8461537 | 4.94E-05 |
| STK36 | 1.29209169 | 18.1657391 | 7.76E-05 |
| CUL3 | 1.28382875 | 20.3560288 | 2.77E-07 |
| POM121C | 1.28361891 | 23.6302858 | 1.24E-08 |

| | | | |
|---|---|---|---|
| P2RY1 | 1.28300331 | 17.9714442 | 1.38E-06 |
| VMP1 | 1.28011001 | 23.2332364 | 0.00535631 |
| STAG2 | 1.27998331 | 19.7768424 | 1.13E-07 |
| SHPRH | 1.27947388 | 18.422688 | 2.69E-07 |
| NDUFC2 | 1.27926681 | 22.8343441 | 0.00158018 |
| TMEM181 | 1.27865217 | 23.0275421 | 3.04E-08 |
| ATP9A | 1.27800014 | 19.7981104 | 3.64E-06 |
| MAGEA3 | 1.27684982 | 16.5460134 | 0.00166011 |
| SURF4 | 1.27603746 | 24.4533742 | 3.49E-05 |
| PRR14 | 1.27388154 | 19.5953158 | 1.01E-06 |
| FZD3 | 1.27376378 | 19.3244321 | 3.64E-06 |
| SASS6 | 1.27349855 | 20.8361272 | 4.08E-05 |
| YIPF3 | 1.26936353 | 22.5012071 | 1.71E-05 |
| C15ORF61 | 1.26781648 | 18.81238 | 1.18E-06 |
| TMEM128 | 1.26607115 | 19.7058325 | 1.15E-07 |
| LAGE3 | 1.26316616 | 17.967276 | 2.87E-07 |
| ROGDI | 1.26308716 | 17.6240595 | 2.30E-04 |
| NDFIP2 | 1.26160173 | 18.5521636 | 1.21E-05 |
| NAA20 | 1.26099711 | 18.6764435 | 2.63E-08 |
| PIGG | 1.25937592 | 21.8374034 | 4.26E-04 |
| SLC43A2 | 1.25765375 | 19.2339933 | 0.00457346 |
| MKLN1 | 1.25710281 | 18.8360555 | 3.96E-05 |
| RAB34 | 1.25664689 | 19.2757994 | 2.12E-04 |

| | | | |
|---|---|---|---|
| TMEM39B | 1.25662978 | 21.1531507 | 2.40E-04 |
| AGPAT2 | 1.25457809 | 22.7709514 | 0.0012123 |
| ADGRL3 | 1.2534719 | 19.9891178 | 1.62E-05 |
| TMEM177 | 1.2531992 | 23.3222736 | 0.00557145 |
| LONRF3 | 1.25212547 | 19.430717 | 6.42E-07 |
| NPC1 | 1.25197783 | 20.4811237 | 3.63E-05 |
| NIF3L1 | 1.25092774 | 17.609986 | 1.91E-07 |
| FAM210B | 1.25067018 | 18.2283241 | 4.18E-07 |
| INTS1 | 1.24866781 | 20.8421957 | 0.09756147 |
| TIMM50 | 1.24781796 | 24.4256583 | 3.58E-06 |
| STOM | 1.24763082 | 18.7096503 | 9.52E-05 |
| TRMT1 | 1.24190485 | 19.4035161 | 3.74E-06 |
| EBP | 1.2403496 | 23.4468507 | 1.04E-05 |
| FIRRM | 1.24016953 | 20.2585688 | 4.77E-05 |
| SMO | 1.23903558 | 18.2852628 | 8.29E-07 |
| DCD | 1.23826213 | 23.0669918 | 0.08246439 |
| TMEM209 | 1.23754954 | 21.5404332 | 4.65E-04 |
| POLR2E | 1.23733365 | 17.5944541 | 2.89E-04 |
| DRAM2 | 1.23655455 | 21.2433762 | 7.21E-06 |
| FBXL12 | 1.23577499 | 20.9943676 | 4.45E-05 |
| ATP2A2 | 1.23542891 | 24.4735981 | 4.89E-08 |
| IMPACT | 1.23344629 | 18.6669073 | 1.03E-04 |
| ATP8B2 | 1.23245443 | 20.7605307 | 1.29E-06 |

| | | | |
|---|---|---|---|
| JADE3 | 1.23217024 | 18.9112401 | 6.47E-06 |
| TBC1D9B | 1.23180519 | 22.1152182 | 0.00935899 |
| PTDSS1 | 1.22936484 | 24.5884906 | 0.05699601 |
| SAAL1 | 1.22787605 | 21.4061278 | 1.44E-04 |
| SLC41A3 | 1.22718158 | 19.2217559 | 3.53E-06 |
| TMEM179B | 1.22563357 | 18.4221146 | 6.64E-08 |
| CHP1 | 1.2187103 | 22.0606866 | 4.23E-04 |
| RHBDD1 | 1.21831711 | 21.5830753 | 8.41E-08 |
| WFS1 | 1.21812276 | 21.3421203 | 8.43E-08 |
| ZDHHC12 | 1.21540481 | 18.0627527 | 1.25E-04 |
| RNF128 | 1.21523501 | 21.6625666 | 1.77E-04 |
| ZNF414 | 1.21467603 | 20.6852362 | 0.00151267 |
| CLEC16A | 1.21412218 | 18.6984623 | 9.19E-05 |
| TMEM168 | 1.21311455 | 17.9170505 | 3.23E-06 |
| RANBP6 | 1.21261218 | 18.5995053 | 0.01304812 |
| COG4 | 1.21245613 | 20.1739929 | 3.46E-09 |
| PDGFRL | 1.21184385 | 20.1527263 | 4.44E-04 |
| SLC20A1 | 1.21155912 | 21.4413643 | 6.85E-06 |
| WIZ | 1.21018851 | 19.3867662 | 6.72E-06 |
| S100A7 | 1.20824429 | 20.5325947 | 0.11234637 |
| ECH1 | 1.20705076 | 18.9089069 | 6.54E-08 |
| STX18 | 1.20640204 | 21.3414271 | 0.00320855 |
| RETREG2 | 1.20485824 | 20.3111815 | 8.68E-07 |

| | | | |
|---|---|---|---|
| ABCB8 | 1.20416223 | 19.4090779 | 2.13E-06 |
| APBB1 | 1.19868917 | 20.6689774 | 9.79E-07 |
| MFSD9 | 1.19808703 | 18.8939519 | 2.51E-04 |
| ZDHHC4 | 1.19442203 | 19.052575 | 2.73E-06 |
| KTN1 | 1.19326921 | 24.8091963 | 3.68E-05 |
| VKORC1L1 | 1.19026336 | 22.1646993 | 4.38E-06 |
| EMC6 | 1.18966975 | 24.3660747 | 1.86E-04 |
| ZFP14;ZFP30 | 1.18950374 | 18.4932941 | 1.45E-07 |
| LSS | 1.18754173 | 17.1226475 | 0.19412241 |
| SREBF1 | 1.18521764 | 21.6442067 | 3.99E-05 |
| PIGK | 1.18419386 | 23.5109474 | 4.80E-05 |
| FADS3 | 1.18146576 | 18.6269885 | 2.52E-04 |
| HMGCR | 1.17728685 | 19.5525934 | 6.49E-06 |
| SHMT2 | 1.17594357 | 22.3826957 | 7.44E-04 |
| ATG7 | 1.17521561 | 18.575982 | 3.34E-08 |
| NAF1 | 1.17379712 | 18.720482 | 0.00154476 |
| ENTREP1 | 1.17350491 | 17.5276163 | 5.27E-08 |
| SLC12A2 | 1.17319193 | 21.4289883 | 3.98E-06 |
| NELFA | 1.1716868 | 21.9451119 | 0.05174814 |
| PRELID1 | 1.16877298 | 16.3349214 | 0.02931325 |
| ALG6 | 1.16805285 | 22.3706552 | 0.00787195 |
| GPAA1 | 1.16700301 | 22.954029 | 4.31E-07 |
| TMX1 | 1.1662015 | 21.8103399 | 1.39E-05 |

| | | | |
|---|---|---|---|
| SLC25A36 | 1.16556853 | 20.8093294 | 2.29E-07 |
| MLH1 | 1.15862052 | 20.1942434 | 0.080376 |
| STEAP1 | 1.15784473 | 19.5459896 | 2.19E-06 |
| TMEM87A | 1.15632739 | 19.0500489 | 2.02E-04 |
| PSEN1 | 1.15566159 | 21.3952784 | 0.00192768 |
| TAC1 | 1.15508256 | 18.9072182 | 3.49E-05 |
| COX7B | 1.15444553 | 22.614919 | 0.0048409 |
| PSD3 | 1.15241837 | 17.9729052 | 1.12E-04 |
| RHEB | 1.15053412 | 21.1762158 | 0.00384407 |
| TUBB6 | 1.15008512 | 29.9652959 | 5.56E-07 |
| C19ORF25 | 1.14931733 | 22.4291736 | 3.75E-04 |
| SFXN3 | 1.1471092 | 20.6957828 | 0.00180628 |
| SLC16A2 | 1.14707145 | 20.7059538 | 1.21E-05 |
| TMEM187 | 1.14620121 | 18.7549962 | 3.11E-05 |
| PC | 1.14453548 | 20.4190871 | 0.0916435 |
| VAC14 | 1.14403674 | 21.6122728 | 0.00129425 |
| MRPL11 | 1.14395006 | 18.923934 | 1.42E-06 |
| TELO2 | 1.14291727 | 24.6357203 | 8.33E-05 |
| ARMCX6 | 1.14235319 | 16.3669365 | 5.06E-05 |
| SLC7A6 | 1.14109993 | 22.5816016 | 6.63E-08 |
| WLS | 1.13966683 | 24.772327 | 6.97E-05 |
| SLC35F6 | 1.13966504 | 22.7718171 | 0.00406605 |
| C5ORF34 | 1.13723258 | 19.0023899 | 3.68E-05 |

| | | | |
|---|---|---|---|
| KNSTRN | 1.13699741 | 19.0864167 | 0.00394675 |
| RNF170 | 1.13541495 | 20.6889993 | 2.01E-06 |
| ATP2C1 | 1.13435731 | 20.9907005 | 1.74E-05 |
| MFSD5 | 1.13212869 | 19.9181637 | 1.31E-04 |
| EEFSEC | 1.1273404 | 18.8343039 | 0.04271112 |
| OSBPL8 | 1.12679968 | 24.0477627 | 6.72E-07 |
| PRKDC | 1.12669552 | 27.3477618 | 2.98E-07 |
| CNIH4 | 1.12638345 | 25.6871591 | 6.14E-04 |
| PCNX3 | 1.12518055 | 20.571971 | 1.50E-08 |
| SIK2 | 1.12355223 | 19.4814122 | 6.53E-07 |
| KIAA2013 | 1.12078099 | 21.3484461 | 1.01E-05 |
| KCNG3 | 1.12047136 | 16.9796957 | 0.03239189 |
| SLC35D3 | 1.11998031 | 18.15255 | 9.79E-07 |
| TCTN2 | 1.11995409 | 19.3765728 | 7.74E-04 |
| UBXN8 | 1.11952795 | 20.8542705 | 6.92E-08 |
| SYNE1 | 1.11916859 | 18.0476179 | 8.52E-05 |
| ATP1B1 | 1.11879943 | 21.8723344 | 0.00526607 |
| NUP37 | 1.11712693 | 19.9401158 | 0.0432311 |
| ANKH | 1.1150471 | 20.4838296 | 1.87E-04 |
| PI4KA | 1.11501612 | 22.2683447 | 2.44E-04 |
| GP1BB | 1.11299482 | 20.7478586 | 5.08E-07 |
| PTMA | 1.11283067 | 17.4596842 | 3.73E-04 |
| TRPM7 | 1.1120878 | 19.8900043 | 1.52E-05 |

| | | | |
|---|---|---|---|
| CYP27C1 | 1.11085908 | 19.4584462 | 0.00605928 |
| TENT4A | 1.11074158 | 16.8265747 | 4.78E-05 |
| SEC62 | 1.11008031 | 20.520531 | 1.15E-05 |
| ZDHHC18 | 1.10852612 | 18.9633926 | 0.00461674 |
| DPY19L4 | 1.10660082 | 20.5940938 | 0.00182219 |
| CHRNA5 | 1.1040844 | 21.1163773 | 3.21E-04 |
| ICAM5 | 1.10233199 | 19.5221615 | 8.81E-06 |
| CHRFAM7A;CHRNA7 | 1.10230409 | 18.6953408 | 1.57E-05 |
| CCDC134 | 1.10155998 | 20.5229208 | 8.52E-04 |
| SLC39A7 | 1.10037205 | 23.6050454 | 4.80E-04 |
| EMC10 | 1.09941617 | 19.4181517 | 3.42E-05 |
| SOAT1 | 1.09832034 | 20.1799689 | 9.32E-06 |
| BRAT1 | 1.0961728 | 21.7651608 | 0.00147354 |
| MCM3AP | 1.09592137 | 20.291839 | 0.05184143 |
| XPO6 | 1.0958953 | 21.7676281 | 1.10E-07 |
| ZNF446 | 1.09278904 | 19.4401941 | 3.09E-06 |
| CCDC8 | 1.09146672 | 21.1564736 | 4.47E-06 |
| MTDH | 1.08977831 | 21.1641065 | 9.44E-06 |
| ALDH1B1 | 1.08899361 | 26.0479943 | 0.04156108 |
| ACADS | 1.08763249 | 17.5055652 | 4.54E-06 |
| HOOK2 | 1.0871547 | 20.1776969 | 4.50E-04 |
| EPDR1 | 1.08697495 | 19.5121803 | 6.14E-04 |
| GPR137 | 1.08500357 | 20.3834711 | 7.54E-04 |

| | | | |
|---|---|---|---|
| QTRT1 | 1.08372484 | 19.1478116 | 8.39E-06 |
| MRPL20 | 1.08310581 | 21.03133 | 0.03040921 |
| GRAMD2B | 1.08255813 | 18.5373864 | 1.74E-05 |
| GDAP2 | 1.0816474 | 16.7680239 | 1.25E-04 |
| QPRT | 1.08117053 | 18.1658907 | 0.00291568 |
| PTDSS2 | 1.08115558 | 22.6671745 | 2.13E-06 |
| NAT8L | 1.07867462 | 19.6798735 | 5.15E-04 |
| ABI1 | 1.07833202 | 17.4491447 | 0.01992355 |
| URB2 | 1.07447596 | 19.9633867 | 0.00222316 |
| TMEM231 | 1.07117896 | 21.183951 | 5.88E-05 |
| MRPL3 | 1.0672455 | 19.0477432 | 0.02377554 |
| CCDC115 | 1.06681129 | 20.5882013 | 4.08E-05 |
| ORMDL2 | 1.06643761 | 19.8989637 | 0.03977908 |
| SPPL2B | 1.066058 | 20.9858663 | 1.95E-07 |
| MCCC1 | 1.06574283 | 18.6058581 | 1.61E-05 |
| DYM | 1.06573776 | 21.2579921 | 4.32E-05 |
| SEC11A | 1.06546889 | 23.9927345 | 1.89E-05 |
| TMED7 | 1.06362855 | 31.4371682 | 0.01761828 |
| SNX19 | 1.06334249 | 17.7598347 | 2.56E-06 |
| SSR1 | 1.06298364 | 26.1441665 | 0.02280878 |
| CRTC2 | 1.06050952 | 18.4317523 | 0.04336357 |
| SLC10A7 | 1.06045974 | 18.3755042 | 6.99E-05 |
| FZD6 | 1.0602702 | 20.3397043 | 1.30E-05 |

| | | | |
|---|---|---|---|
| COX10 | 1.05917618 | 18.2413239 | 6.28E-06 |
| ANO8 | 1.05760003 | 17.8972937 | 7.10E-04 |
| APPBP2 | 1.05677158 | 21.2204933 | 3.99E-04 |
| TAP1 | 1.0558433 | 22.0666394 | 0.01537746 |
| TCEAL9 | 1.05522201 | 18.5083548 | 9.62E-05 |
| CERS5 | 1.05305145 | 22.2964319 | 2.06E-04 |
| CCT6B | 1.05292151 | 21.3199052 | 2.49E-07 |
| RLIM | 1.05291561 | 21.3673365 | 9.90E-05 |
| RCCD1 | 1.05204296 | 18.0315866 | 2.92E-05 |
| ASPHD2 | 1.05100533 | 21.7556564 | 7.10E-06 |
| TMEM208 | 1.0487301 | 21.9926276 | 1.50E-05 |
| COX18 | 1.0484167 | 22.1868078 | 1.83E-05 |
| NAV1 | 1.04799152 | 20.7423451 | 4.90E-07 |
| PLOD1 | 1.04765372 | 23.4748187 | 9.04E-04 |
| TMED4 | 1.04612328 | 24.4645578 | 1.65E-05 |
| GPR108 | 1.04565397 | 19.6261206 | 8.36E-06 |
| RHBDF1 | 1.04425358 | 19.753097 | 8.82E-06 |
| LPCAT2 | 1.04325546 | 20.1138418 | 2.20E-05 |
| GDE1 | 1.04183255 | 17.9027104 | 8.06E-05 |
| SLC16A10 | 1.04075097 | 20.5393851 | 5.38E-05 |
| DPY19L2 | 1.04020543 | 16.7840616 | 1.71E-05 |
| RPL10L | 1.03926917 | 18.6381067 | 5.45E-04 |
| COL25A1 | 1.03855003 | 20.3317582 | 1.83E-05 |

| | | | |
|---|---|---|---|
| GEMIN2 | 1.03647452 | 20.6511399 | 7.67E-05 |
| TMEM259 | 1.03458974 | 19.6478599 | 1.03E-04 |
| FXYD5 | 1.03439474 | 18.0544315 | 1.04E-05 |
| TAPT1 | 1.03336152 | 19.2473217 | 5.89E-06 |
| TMEM223 | 1.03285078 | 24.142024 | 0.00227136 |
| HTT | 1.03242318 | 20.0326766 | 2.92E-04 |
| PIGT | 1.03214328 | 24.2761741 | 2.72E-04 |
| TM9SF4 | 1.03194 | 21.6484528 | 3.18E-08 |
| DYRK1A | 1.03019029 | 19.8413729 | 0.00465896 |
| C15ORF39 | 1.03010952 | 21.5973201 | 7.92E-05 |
| MT-ND3 | 1.03001591 | 18.1385475 | 1.53E-06 |
| ATXN1L | 1.02991544 | 15.6976694 | 8.20E-05 |
| TMEM132E | 1.02867448 | 20.3569992 | 0.13942317 |
| SLC35C1 | 1.02809287 | 19.5194174 | 1.48E-04 |
| MATCAP2 | 1.02700348 | 18.7528046 | 1.99E-04 |
| SFXN2 | 1.02622939 | 22.9852082 | 1.86E-05 |
| TCF7L2 | 1.02419486 | 18.3676373 | 2.11E-06 |
| NR2F1 | 1.02352617 | 18.6900447 | 9.52E-06 |
| SLC17A5 | 1.02348968 | 18.7439808 | 1.52E-04 |
| OXA1L | 1.02247001 | 27.0869208 | 9.22E-05 |
| ATR | 1.02188893 | 21.8273434 | 1.08E-07 |
| CISD3 | 1.02170997 | 20.3614837 | 0.00221816 |
| ACTR3B | 1.02106995 | 18.5733484 | 7.22E-06 |

| | | | |
|---|---|---|---|
| BCL2L11 | 1.019641 | 17.7395374 | 7.90E-04 |
| SNRPE | 1.01932108 | 23.3865847 | 0.002336 |
| TADA1 | 1.01905783 | 19.424338 | 7.71E-04 |
| NIPA2 | 1.01838296 | 19.0917678 | 4.65E-04 |
| SLC25A29 | 1.0181818 | 21.0289247 | 1.40E-05 |
| SLC5A3 | 1.01762244 | 21.0102518 | 2.91E-05 |
| RARS2 | 1.01717862 | 23.2864467 | 6.87E-07 |
| ZBED3 | 1.01696612 | 19.846109 | 4.54E-04 |
| VPS29 | 1.01403149 | 17.803301 | 3.88E-06 |
| ELP4 | 1.01309063 | 17.6058769 | 0.07218464 |
| ZDHHC13 | 1.01304168 | 22.778945 | 7.76E-04 |
| NDUFB5 | 1.01285744 | 23.0448945 | 1.02E-04 |
| GLRX | 1.01044717 | 17.3940675 | 2.94E-07 |
| CASC3 | 1.01006402 | 18.3053945 | 4.05E-06 |
| BLTP2 | 1.00675603 | 19.1244389 | 6.78E-04 |
| TPCN2 | 1.00666667 | 18.3783212 | 1.02E-06 |
| TENT4B | 1.00504604 | 16.2972769 | 1.04E-05 |
| QSOX1 | 1.00283185 | 21.2814072 | 9.47E-05 |
| FBXO9 | 1.0018004 | 18.2034938 | 2.77E-04 |
| EVI5 | 1.00063606 | 20.7027237 | 5.71E-05 |
| ARL6IP5 | 1.00053118 | 21.361235 | 6.53E-07 |

**Table S7. TMED antibodies.**

List of antibodies detecting TMEDs used for Western blot and immunofluorescence throughout the manuscript. IF, Immunofluorescence.

| Antigen (Host) | Company, Cat# | Application | KO validated |
|---|---|---|---|
| TMED1 (rabbit)<br><br>TMED1 (mouse) | Sigma, HPA018507<br><br>Santa Cruz, sc377321 | Western, 1:1000<br>IF, 1:100<br>IF, 1:100 | Yes (Fig. S1)<br><br>Yes (not shown) |
| TMED2 (mouse)<br>TMED2 (rabbit) | Santa Cruz, sc-378459<br>Proteintech, 11981-1-AP | Western, 1:500<br>IF, 1:100 | Yes (Fig. S1)<br>Yes (not shown) |
| TMED4 (rabbit) | LSBio, LS-C783502 | Western, 1:1000 | Yes (Fig. S1) |
| TMED5 (rabbit) | Thermo, PA5-31580 | Western, 1:1000<br>IF, 1:500 | Yes (Fig. S1) |
| TMED7 (rabbit) | Atlas, HPA008960 | Western, 1:1000<br>IF, 1:500 | Yes (Fig. S1) |
| TMED9 (rabbit) | Proteintech, 21620-1-AP | Western, 1:1000<br>IF, 1:1000 | Yes (Fig. S1) |
| TMED10 (mouse)<br><br>TMED10 (rabbit) | Santa Cruz, sc137003 HRP<br>Santa Cruz, sc137003<br>Bethyl, A305-218A | Western, 1:500<br>IF, 1:100<br>IF, 1:100 | Yes (Fig. S1)<br><br>Yes (not shown) |
